# Systematic analysis reveals an Mln1-dependent trade-off between metabolic flexibility and thermotolerance in *Cryptococcus neoformans*

**DOI:** 10.64898/2026.10.30.763006

**Authors:** Seong-Ryong Yu, Minjae Lee, So-Yeon Yang, Hyunjin Cha, Doyeon Won, Dohee Kim, Jin-Tae Choi, Soojin Yu, Ran Seo, Kyung-Ah Lee, Jong-Seung Lee, Won-Jae Lee, Dong-Woo Lee, Yong-Sun Bahn

## Abstract

During infection, fungal pathogens must adapt to multiple environmental challenges, including changes in temperature, nutrient availability, and immune stress. Although transcriptional regulators are central to these adaptations, how individual regulators contribute to distinct environmental responses and how these responses are functionally interconnected remain poorly understood. Here, we investigated the functions of infection-associated transcription factors under distinct host-relevant conditions in *Cryptococcus neoformans*. Integrating in vivo expression profiling with infectivity data from signature-tagged mutagenesis, we prioritized 24 transcription factors, 12 of which were induced under host-mimicking conditions. Initial functional analyses revealed distinct roles for these transcription factors in adaptation to temperature, carbon, and nitrogen availability. Further characterization of Mln1, a fungal basic helix-loop-helix transcription factor, uncovered an unexpected connection between carbon source adaptation and thermotolerance. Mig1-dependent repression limited *MLN1* expression in the presence of glucose, whereas Mln1 accumulated in the nucleus in the presence of trehalose or ethanol, but not glucose. Loss of *MLN1* impaired carbon-starvation gene expression and growth on non-preferred carbon sources, particularly trehalose. Conversely, *mln1*Δ cells accumulated more intracellular trehalose and exhibited enhanced thermotolerance, yet showed reduced melanization, titan cell formation, virulence in *Drosophila*, and brain fungal burden in mice. Together, these results suggest that Mln1 contributes to metabolic flexibility and fungal virulence while differentially influencing thermotolerance. More broadly, these findings highlight a potential physiological trade-off between nutrient utilization and stress resistance, suggesting that these adaptive processes may be interconnected in fungal pathogens.

## Introduction

Pathogenic fungi must rapidly adapt to environmental changes as they transition from environmental reservoirs to mammalian hosts. This transition exposes them to elevated temperature, altered gas concentrations, nutrient limitation, and immune-derived stresses that can compromise their survival and pathogenicity (1–3). *Cryptococcus neoformans*, an encapsulated basidiomycete and a major cause of life-threatening fungal meningitis, has been classified as a critical-priority fungal pathogen by the World Health Organization (4, 5). Widely distributed in soil, decaying wood, and bird guano (6, 7), *C. neoformans* can enter the human host through inhalation of infectious propagules and subsequently disseminate from the lungs to the central nervous system (8–11). Successful infection therefore requires the coordinated regulation of environmental stress responses and virulence-associated traits, including capsule formation and melanization (12, 13).

Among the challenges encountered during infection, elevated temperature and variable carbon availability impose particularly important demands on fungal adaptation. Thermotolerance is a key virulence determinant, distinguishing pathogenic *Cryptococcus* species, which can grow at temperatures up to 39°C, from many closely related environmental fungi (14, 15). Meanwhile, the abundance and composition of available carbon sources vary substantially across host compartments (16). Glucose availability is relatively limited at the pulmonary airway surface (17, 18), and *C. neoformans* cells recovered during early lung infection exhibit transcriptional responses associated with alternative carbon source utilization (19). In the brain, glucose serves not only as a nutrient but also as a signal that promotes tolerance to amphotericin B through Mig1-dependent regulation, while abundant myo-inositol provides an alternative carbon source whose uptake and metabolism have been implicated in blood–brain barrier traversal (20–23). Carbon availability also influences virulence-associated processes, including melanization and the expression of antiphagocytic factors (13, 24). Adaptation to these diverse conditions involves conserved signaling pathways, including cyclic AMP–protein kinase A, mitogen-activated protein kinases, calcineurin, and Snf1, which coordinate stress responses and metabolic regulation through downstream transcriptional regulators (19, 25–28).

Trehalose metabolism provides a potential link between carbon utilization and thermal stress resistance. In fungi, intracellular trehalose serves as a mobilizable carbon reserve and helps protect cellular proteins during heat stress, while extracellular trehalose can also serve as an alternative carbon source. The degradation of intracellular trehalose following stress can facilitate cellular recovery, as demonstrated in *Saccharomyces cerevisiae* (29, 30). In *C. neoformans*, elevated temperature promotes trehalose accumulation and induces the trehalose biosynthetic genes *TPS1* and *TPS2*, both of which are required for growth at mammalian body temperature (31). Thus, trehalose metabolism must accommodate two distinct physiological demands: the utilization of trehalose as a carbon source and the maintenance of intracellular trehalose for stress protection. How these functions are coordinated across changing nutritional and thermal conditions, however, remains incompletely understood.

Despite extensive characterization of fungal stress signaling and metabolic pathways, the transcriptional regulators that respond to individual host-associated environmental cues remain poorly defined. During infection, elevated temperature and nutrient limitation occur simultaneously, making it difficult to distinguish cue-specific transcriptional responses and their functional consequences. Our previous genome-wide analysis identified 45 *C. neoformans* transcription factors (TFs) associated with virulence or infectivity (32). Together with published in vivo transcriptional profiling and signature-tagged mutagenesis datasets (32, 33), these findings provide a foundation for systematically identifying regulators of host-associated adaptation. However, whether transcriptional regulators of nutrient adaptation also influence thermal stress resistance remains largely unexplored.

Here, we integrated previously published transcriptional and infectivity datasets to prioritize 24 infection-associated TFs and systematically examined their responses to individual host-associated cues. We identified 12 TFs induced under host-mimicking conditions and resolved partially overlapping transcriptional responses to elevated temperature, carbon limitation, and nitrogen limitation. Temperature-responsive TFs supported growth at febrile temperatures, whereas the nitrogen-responsive regulator Sre1 was required for growth when amino acids served as the sole nitrogen source. The fungal-restricted, carbon-responsive regulator Mln1 promoted the utilization of selected alternative carbon sources, particularly trehalose, yet its deletion unexpectedly increased basal intracellular trehalose abundance and resistance to acute heat stress. Loss of Mln1 also reduced virulence in *Drosophila* and fungal burden in mouse brains. These findings connect Mln1-dependent carbon adaptation with trehalose homeostasis and thermoprotection, suggesting a potential trade-off between nutrient utilization and thermal stress resistance in a pathogenic fungus.

## Results

### Identification of host-responsive TFs involved in cryptococcal pathogenicity

To identify TFs that mediate cryptococcal responses to host-associated cues, we integrated two previously published datasets (Fig. 1A). The first dataset was a NanoString-based in vivo transcriptional profile measuring the expression of 180 TFs across lung, brain, kidney, and spleen at 3, 7, 14, and 21 days post-infection (33). The second dataset was an STM screen of TF mutants assessed via three independent infection routes, including intranasal (IN), intravenous (IV), and intracerebroventricular (ICV) (32, 33). Two filtering criteria were then applied. First, TFs showing ≥ 2-fold induction at least once in both the brain and lung relative to the in vitro baseline were retained, yielding 52 candidates (Supplementary Information, Fig. S1A, and Dataset S1). Second, among these, TF mutants with an STM score ≤ -2 (a fourfold or greater reduction) in at least one infection model were selected, narrowing the list to 24 candidates (Supplementary Information, Fig. S1B).

**Figure 1.**
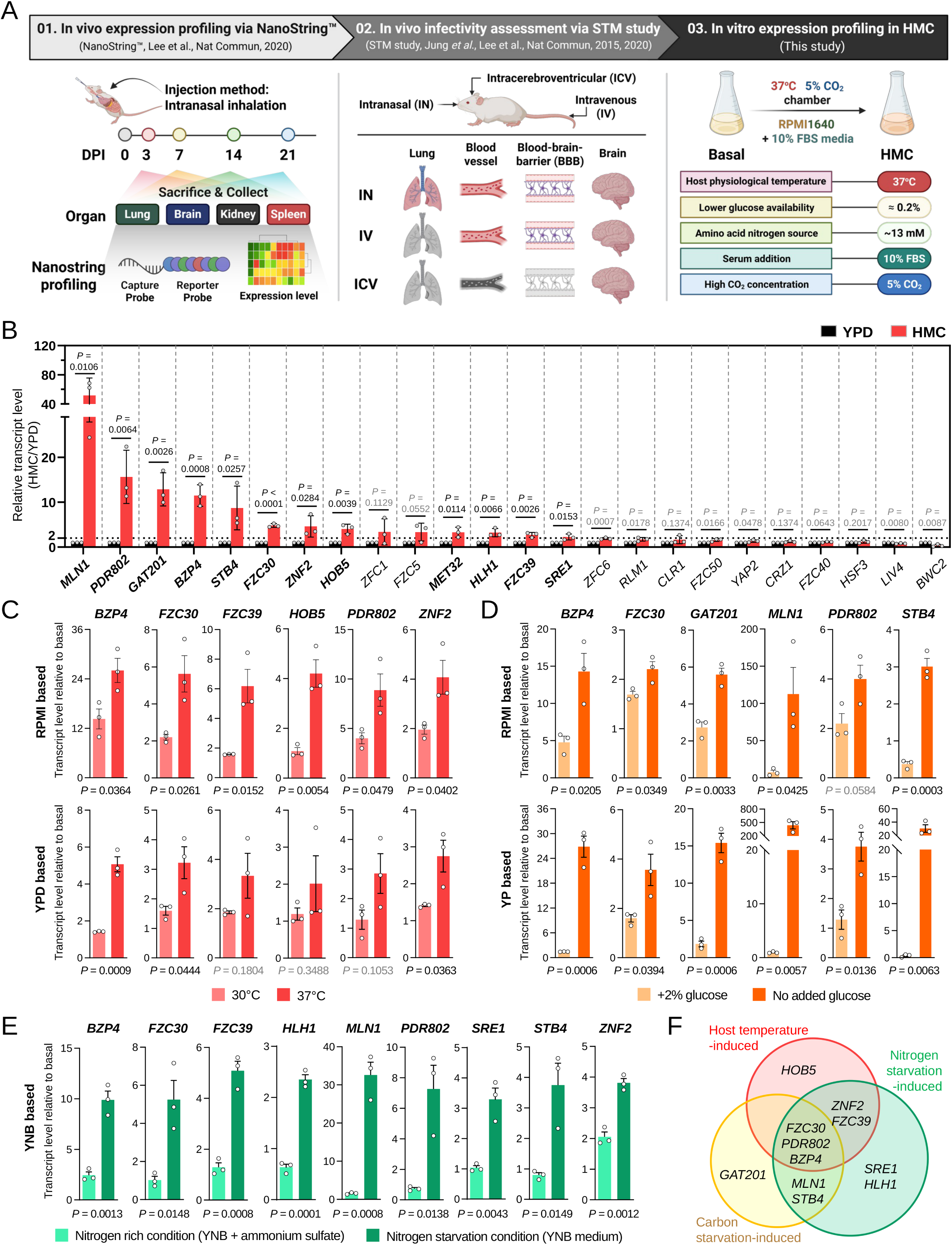
Identification and classification of host-responsive transcription factors in *Cryptococcus neoformans*. (A) Schematic overview of the stepwise selection of transcription factors (TFs) using previously published in vivo NanoString expression and signature-tagged mutagenesis (STM) infectivity data, followed by validation under host-mimicking conditions (HMC). HMC consisted of RPMI 1640 medium supplemented with 10% fetal bovine serum (FBS) at 37°C under 5% CO_2_. The schematic was created with BioRender (https://BioRender.com/45upvnl). (B) Quantitative reverse transcription PCR (qRT-PCR) analysis of 24 candidate TFs under YPD and HMC conditions. Transcript levels were normalized to *ACT1* and expressed relative to YPD. Genes showing >2-fold induction and *P* < 0.05 are indicated by bold gene names and black *P* values. Data represent the mean ± SD of three biological replicates. Statistical significance was determined using a one-tailed Student’s t-test. (C–E) qRT-PCR analysis of the 12 HMC-induced TFs under individual host-associated cues. Each panel shows TFs meeting the induction criteria. (C) Temperature responses at 30°C and 37°C in RPMI and YPD media. (D) Glucose responses in RPMI with or without an additional 2% glucose and in YP versus YPD at 30°C. (E) Nitrogen responses in YNB containing 2% glucose without amino acids, with or without ammonium sulfate. Transcript levels were normalized to *ACT1* and expressed relative to the pre-shift basal sample. Data represent the mean ± SD of three biological replicates, each with three technical replicates. Dots indicate individual biological replicates. Statistical significance was determined using unpaired two-tailed Student’s t-tests. Primers and media compositions are listed in Supplementary Information, Tables S2 and S3, respectively. (F) Venn diagram showing the overlap among TFs induced by elevated temperature, carbon starvation, and nitrogen starvation. Induction was defined as >2-fold expression relative to basal levels with *P* < 0.05 versus the matched control. For temperature and carbon responses tested in two medium backgrounds, these criteria were required in at least one medium without an opposing directional response in the other.

We next asked which of these 24 candidates respond transcriptionally to host-like conditions in vitro. To approximate the host environment, cells were cultured in RPMI medium supplemented with 10% fetal bovine serum (FBS) at 37°C under 5% CO_2_ (hereafter host-mimicking conditions, HMC) (33). As a control, cells werse grown in yeast extract-peptone-dextrose (YPD) medium at 30°C under ambient air. Twelve TFs, including *MLN1*, *PDR802*, *GAT201*, *BZP4*, *STB4*, *FZC30*, *ZNF2*, *HOB5*, *MET32*, *HLH1*, *FZC39*, and *SRE1*, showed significant induction under HMC, with a fold change > 2 and *P* < 0.05 (Fig. 1B).

### Temperature and nutrient availability regulate pathogenicity-associated TFs

To determine how the 12 HMC-responsive TFs respond to individual host-associated cues, we examined their expression under conditions that varied temperature, CO_2_ concentration, serum supplementation, glucose availability, and nitrogen availability (Supplementary Information, Fig. S2A). Among the cues tested, temperature and nutrient availability produced the strongest transcriptional responses. In contrast, 5% CO_2_ and 10% FBS alone had relatively minor effects on the expression of these TFs (Supplementary Information, Fig. S2B and C). TFs were considered induced when their mean transcript level under the test condition exceeded twofold relative to the pre-shift basal sample and differed significantly from the matched control condition. For temperature and carbon responses, which were evaluated in two medium backgrounds, these criteria had to be met in at least one medium, with no opposing directional response in the other.

A temperature shift from 30°C to 37°C significantly induced six TFs (*BZP4*, *FZC30*, *FZC39*, *HOB5*, *PDR802*, and *ZNF2*) (Fig. 1C), while reduced glucose availability induced six TFs (*BZP4*, *FZC30*, *GAT201*, *MLN1*, *PDR802*, and *STB4*) (Fig. 1D). In RPMI, expression was compared in the presence or absence of an additional 2% glucose, whereas yeast extract-peptone (YP) medium was compared with YPD. Among these, *MLN1* and *STB4* showed the strongest responses under carbon deprivation in YP medium, reaching approximately 439 and 30 times their basal expression levels, respectively (Fig. 1D). Similarly, nitrogen depletion significantly upregulated nine TFs (*BZP4*, *FZC30*, *FZC39*, *HLH1*, *MLN1*, *PDR802*, *SRE1*, *STB4*, and *ZNF2*), with *MLN1* again showing the strongest response, reaching approximately 33 times its basal expression level (Fig. 1E). The strong responsiveness of *MLN1* to both carbon and nitrogen limitation suggests that Mln1 may coordinate metabolic adaptation to nutrient starvation in *C. neoformans*.

Notably, *MET32* was induced under HMC but did not respond to any individual cue tested, suggesting that its induction may depend on combined environmental cues or other features of HMC. *BZP4*, *FZC30*, and *PDR802* responded broadly to all three conditions, whereas other TFs displayed more condition-specific expression patterns: *HOB5* was selectively induced by elevated temperature, *GAT201* responded primarily to carbon limitation, and *SRE1* and *HLH1* were transcriptionally induced only under nitrogen starvation (Fig. 1F).

### Host-responsive TF modules differentially regulate adaptation to temperature and nutrient availability

Having categorized the 12 host-responsive TFs into temperature-, nitrogen-, and carbon-responsive modules, we next examined whether TFs within each module functionally contribute to adaptation to the corresponding host-associated conditions. Adaptation to host temperature is a prerequisite for cryptococcal pathogenesis. Six TFs, *BZP4, FZC30, FZC39, HOB5, PDR802,* and *ZNF2*, were significantly induced at elevated temperature in our screen. Previous phenotypic screening implicated Fzc30 in high-temperature growth (32), but the relative contributions of these TFs across increasing temperatures had not been systematically compared.

To compare their growth across temperatures, we monitored growth of the corresponding single-deletion mutants in a multi-channel bioreactor under controlled temperature conditions. At 37°C, all six mutants grew comparably to wild type (WT) (Fig. 2A), indicating that loss of any one of these temperature-responsive TFs was insufficient to impair growth under the conditions tested. This may reflect functional buffering by other transcriptional regulators or compensatory pathways that maintain growth at host physiological temperature. A graded thermosensitivity emerged as temperature increased. At 38°C, *bzp4*Δ*, fzc30*Δ*, fzc39*Δ, and *hob5*Δ showed impaired growth, whereas *pdr802*Δ and *znf2*Δ remained relatively WT-like. At 39°C, all six mutants exhibited reduced growth to varying degrees (Fig. 2A). These results indicate that the contribution of these temperature-responsive TFs becomes more apparent as thermal stress increases above host physiological temperature.

**Figure 2.**
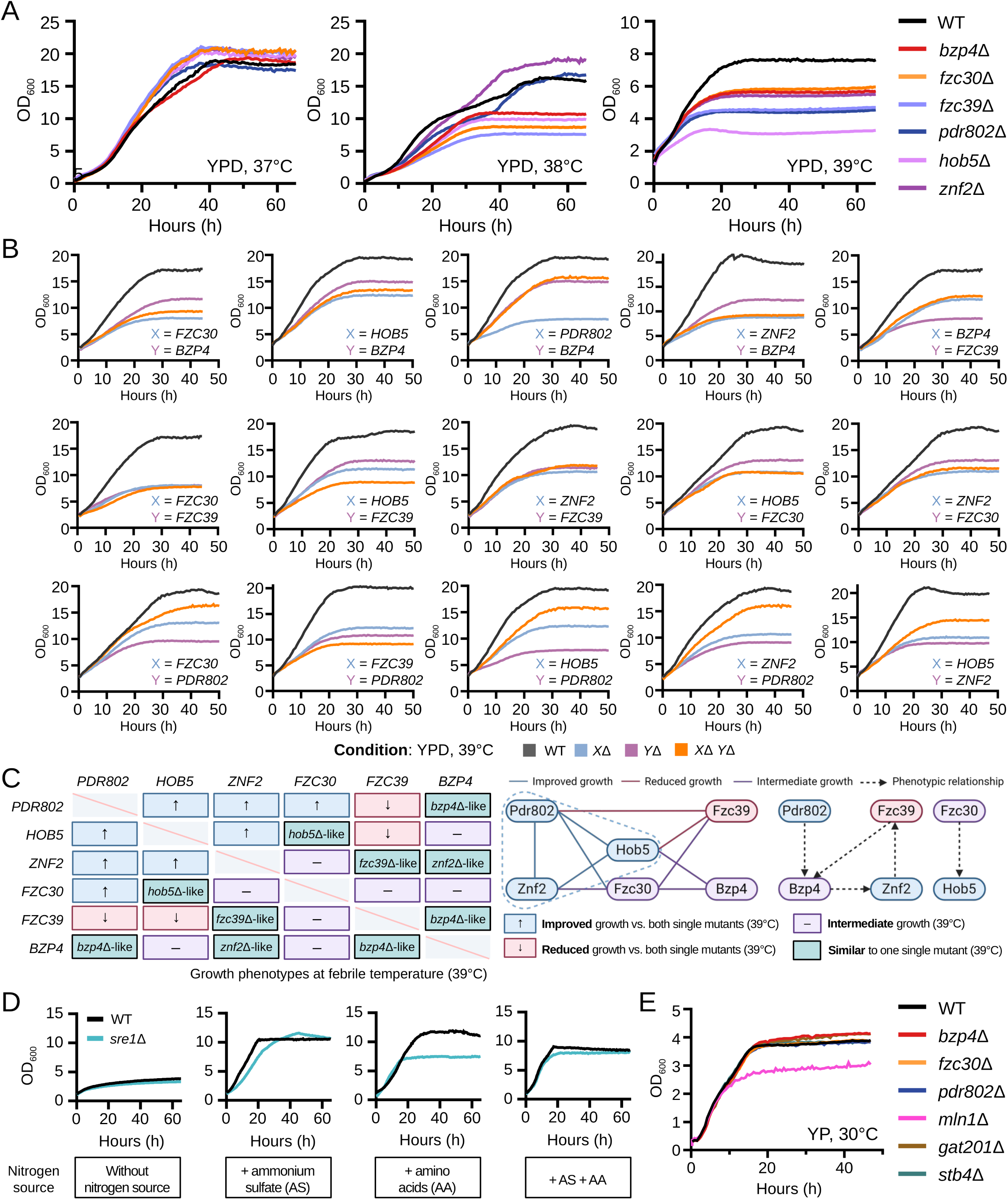
Functional analysis of host-cue-responsive TF modules under temperature, nitrogen, and carbon stress conditions. (A) Growth of WT (H99) and six temperature-responsive TF deletion mutants (*bzp4*Δ, *fzc30*Δ, *fzc39*Δ, *hob5*Δ, *pdr802*Δ, and *znf2*Δ) in YPD medium at 37°C, 38°C, and 39°C. Growth was monitored by measuring OD_600_ using a multi-channel bioreactor. Curves from one biological experiment are shown. (B) Growth kinetics of WT, the indicated single-deletion mutants, and all 15 pairwise double-deletion mutants at 39°C in YPD medium, starting at an OD_600_ of 2.0. In each panel, *A* and *B* represent the indicated TF genes. WT, *X*Δ, *Y*Δ, and *X*Δ *Y*Δ are shown in black, light blue, purple, and orange, respectively. Identical WT or single-mutant control curves are reproduced across panels when measured in the same experiment. (C) Summary of pairwise deletion mutant growth phenotypes at 39°C, a febrile temperature. Matrix and network summarizing the growth phenotypes of the 15 double mutants relative to their corresponding single mutants at 39°C. Blue cells with upward arrows indicate improved growth relative to both single mutants, pink cells with downward arrows indicate reduced growth, purple cells with dashes indicate intermediate growth, and light cyan cells indicate single-mutant-like growth. Diagonal cells are excluded. Network connections use corresponding colors, and node colors indicate the most frequent phenotype category, excluding single-mutant-like relationships. The dashed blue outline highlights Pdr802, Hob5, and Znf2, whose pairwise double deletions showed improved growth relative to both single mutants. Black dashed arrows point toward the TF whose single-deletion phenotype is resembled by the double mutant. The schematic was created with BioRender (https://BioRender.com/45upvnl). (D) Growth of WT and *sre1*Δ in YNB medium containing 2% glucose without amino acids or ammonium sulfate, or supplemented with ammonium sulfate (AS), amino acids (AA), or both. Growth was monitored by OD_600_. Curves from one biological experiment are shown. (E) Growth of WT and six carbon-responsive TF deletion mutants (*bzp4*Δ, *fzc30*Δ, *pdr802*Δ, *mln1*Δ, *gat201*Δ, and *stb4*Δ) in glucose-free YP medium at 30°C, monitored by OD_600_. Curves from one biological experiment are shown.

To determine whether the thermosensitivity of these mutants was associated with impaired induction of the trehalose biosynthetic pathway, we examined the expression of *TPS1* and *TPS2* following a temperature upshift from 30°C to 39°C for 1 h. Both genes were significantly induced after temperature upshift in the WT and in all six deletion mutants (Supplementary Information, Fig. S3), with no significant differences in either basal or 39°C expression levels between the WT and any of the mutants. These results indicate that none of the six temperature-responsive TFs is individually required for temperature-induced transcription of *TPS1* or *TPS2*.

Because functional redundancy could mask thermotolerance defects at 37°C, we generated all 15 pairwise double-deletion strains among the six TFs (Supplementary Information, Fig. S4) and monitored their growth under the same condition. None of the double mutants showed appreciable thermosensitivity at 37°C (Fig. S5). Interestingly, at 39°C, the double mutants exhibited diverse growth patterns depending on the TF combination (Fig. 2B and C). Notably, all three pairwise double mutants among *PDR802*, *HOB5*, and *ZNF2* (*hob5*Δ *pdr802*Δ, *znf2*Δ *pdr802*Δ, and *hob5*Δ *znf2*Δ) showed improved growth compared with either corresponding single mutant. Similar growth improvement was observed in the *fzc30*Δ *pdr802*Δ. In contrast, simultaneous deletion of *FZC39* with either *PDR802* or *HOB5* resulted in more severe growth defects than those observed in the corresponding single mutants. The remaining double mutants exhibited intermediate growth or resembled one of the corresponding single mutants. In particular, *pdr802*Δ *bzp4*Δ and *bzp4*Δ *fzc39*Δ showed growth patterns similar to that of *bzp4*Δ. Together, these findings suggest that the six temperature-responsive TFs have distinct genetic relationships during high-temperature adaptation rather than simply functioning redundantly.

We next examined the nitrogen-responsive module. Nitrogen depletion significantly induced nine TFs in our expression profiling, including *BZP4, FZC30, FZC39, HLH1, MLN1, PDR802, SRE1, STB4,* and *ZNF2*. To determine whether any of these TFs are required for growth on different nitrogen sources, we assessed growth of the corresponding deletion mutants on yeast nitrogen base (YNB) medium containing 2% glucose and supplemented with an amino acid mixture, ammonium sulfate, or both. Among the mutants tested, only *sre1*Δ displayed a nitrogen source-specific defect, whereas the remaining eight mutants grew comparably to WT on all nitrogen sources tested (Supplementary Information, Fig. S6). In medium containing ammonium sulfate as the sole nitrogen source, *sre1*Δ grew comparably to WT, whereas growth was markedly reduced when amino acids were provided as the sole nitrogen source (Fig. 2D). This defect was largely rescued by the addition of ammonium sulfate, indicating that *sre1*Δ is specifically impaired under conditions in which amino acids provide the available nitrogen. These findings provide functional support for the nitrogen-responsive TF classification and suggest an additional role for Sre1 beyond its established functions in sterol regulation and hypoxia adaptation (34).

Finally, we examined the carbon-responsive module. Six TFs, *BZP4, FZC30, GAT201, MLN1, PDR802,* and *STB4*, were induced under reduced glucose availability in our screen. Because carbon-free YNB did not support sufficient growth for quantitative comparison, we assessed the corresponding deletion mutants in glucose-free YP medium, which provided a carbon-limited condition that still permitted measurable growth. Among these mutants, *mln1*Δ showed the most pronounced growth defect, whereas the remaining mutants grew comparably to WT (Fig. 2E). Notably, *MLN1* also showed exceptionally strong transcriptional induction under carbon limitation, reaching approximately 439-fold in glucose-free YP medium (Fig. 1D). Together, these transcriptional and phenotypic findings led us to focus subsequent analyses on Mln1 as a key regulator of carbon adaptation.

### Mln1 is a taxonomically restricted fungal bHLH regulator with broad stress-associated phenotypes

Having identified Mln1 as the major functional hit within the carbon-responsive TF module, we next examined the molecular features and broader phenotypic profile of this regulator. *MLN1* (CNAG_04837) encodes a 551-amino-acid putative TF containing a predicted basic helix-loop-helix (bHLH) DNA-binding domain spanning residues 280-366 (Supplementary Information, Fig. S7A). bHLH TFs typically function as homo- or heterodimers, with the basic region mediating DNA binding and the HLH region mediating dimerization (35, 36). AlphaFold 3 modeling of the Mln1 bHLH domain predicted a helix-rich structure and generated a possible dimeric arrangement of two identical bHLH domains (Supplementary Information, Fig. S7B) (37). However, the predicted dimer showed relatively low inter-chain confidence (ipTM = 0.56; pTM = 0.57), preventing a firm conclusion that Mln1 forms a homodimer. Although the bHLH domain is broadly conserved among eukaryotic TFs, Mln1 itself shows limited sequence conservation and has no clear ortholog in *S. cerevisiae*. The gene was therefore originally designated *MLN1* based on its melanization phenotype (38). Consistent with this limited conservation, comparative BLASTp analysis of the 12 host-cue-responsive TFs showed that Mln1 was poorly conserved across fungi relative to the other candidate TFs (Supplementary Information, Fig. S7C), and EggNOG-based ortholog analysis identified candidate Mln1 orthologs in 104 species, predominantly within Ascomycota, Basidiomycota, and Mucoromycota (Supplementary Information, Fig. S7D and E) (39, 40). Thus, Mln1 represents a taxonomically restricted fungal bHLH regulator.

Our previous large-scale phenotypic analysis further suggested that Mln1 affects multiple stress-associated traits (32). The *mln1*Δ mutant exhibited enhanced sensitivity to oxidative stress and heavy metal stresses, as well as to amphotericin B, fludioxonil, and fluconazole (Supplementary Information, Fig. S7F). Notably, *mln1*Δ showed enhanced rather than reduced growth at 39°C (Supplementary Information, Fig. S7G and H), even though *MLN1* was not significantly induced by elevated temperature in our host-cue expression screen. Beyond these stress-associated phenotypes, we also examined virulence-associated traits previously linked to Mln1. Although an initial niger seed agar assay did not reveal a clear melanization defect (32), *mln1*Δ produced visibly reduced melanin on L-3,4-dihydroxyphenylalanine (L-DOPA) medium (Supplementary Information, Fig. S7I), consistent with the original naming of *MLN1* based on its melanization phenotype (38). The mutant also showed modestly decreased capsule formation under minimal medium conditions (Supplementary Information, Fig. S7J), in contrast to the increased capsule phenotype reported in our previous screen using a different inducing medium (32). All these phenotypes were restored to WT levels by ectopic complementation of *mln1*Δ with a WT copy of *MLN1* (Supplementary Information, Fig. S7 and S8A). Together with the strong induction of *MLN1* under carbon starvation, these stress- and virulence-associated phenotypes prompted us to investigate how Mln1 contributes to carbon-limited growth and nutrient adaptation.

### Mln1 supports condition-dependent utilization of selected non-preferred carbon sources

To determine whether the growth defect of *mln1*Δ under carbon-limited conditions reflects a general defect in carbon adaptation or a specific inability to utilize particular carbon sources, we examined growth of WT, *mln1*Δ, and complemented strains on a panel of carbon sources. These included monosaccharides, disaccharides, sugar alcohols, and non-fermentable carbon sources, tested in both YP- and YNB-based media. Because carbon-source phenotypes can differ between surface-associated and submerged growth, each condition was assessed in parallel by spotting assays on solid media and continuous growth monitoring in liquid cultures.

Across this carbon-source panel, *mln1*Δ exhibited a selective rather than global carbon-utilization defect (Fig. 3A). Growth on monosaccharides, including glucose, fructose, galactose, and xylose, was largely comparable between WT and *mln1*Δ in both YP and YNB media. Similarly, most sugar alcohols and several non-fermentable carbon sources supported near-WT growth of the mutant. In contrast, growth defects became apparent without added glucose and on selected non-preferred carbon sources, most notably disaccharides. The mutant showed impaired growth on maltose, sucrose, and trehalose in a condition-dependent manner, with the strongest and most consistent defect observed on trehalose. The corresponding growth defects observed on solid media were rescued in the complemented strain, confirming that these defects were attributable to loss of *MLN1*.

**Figure 3.**
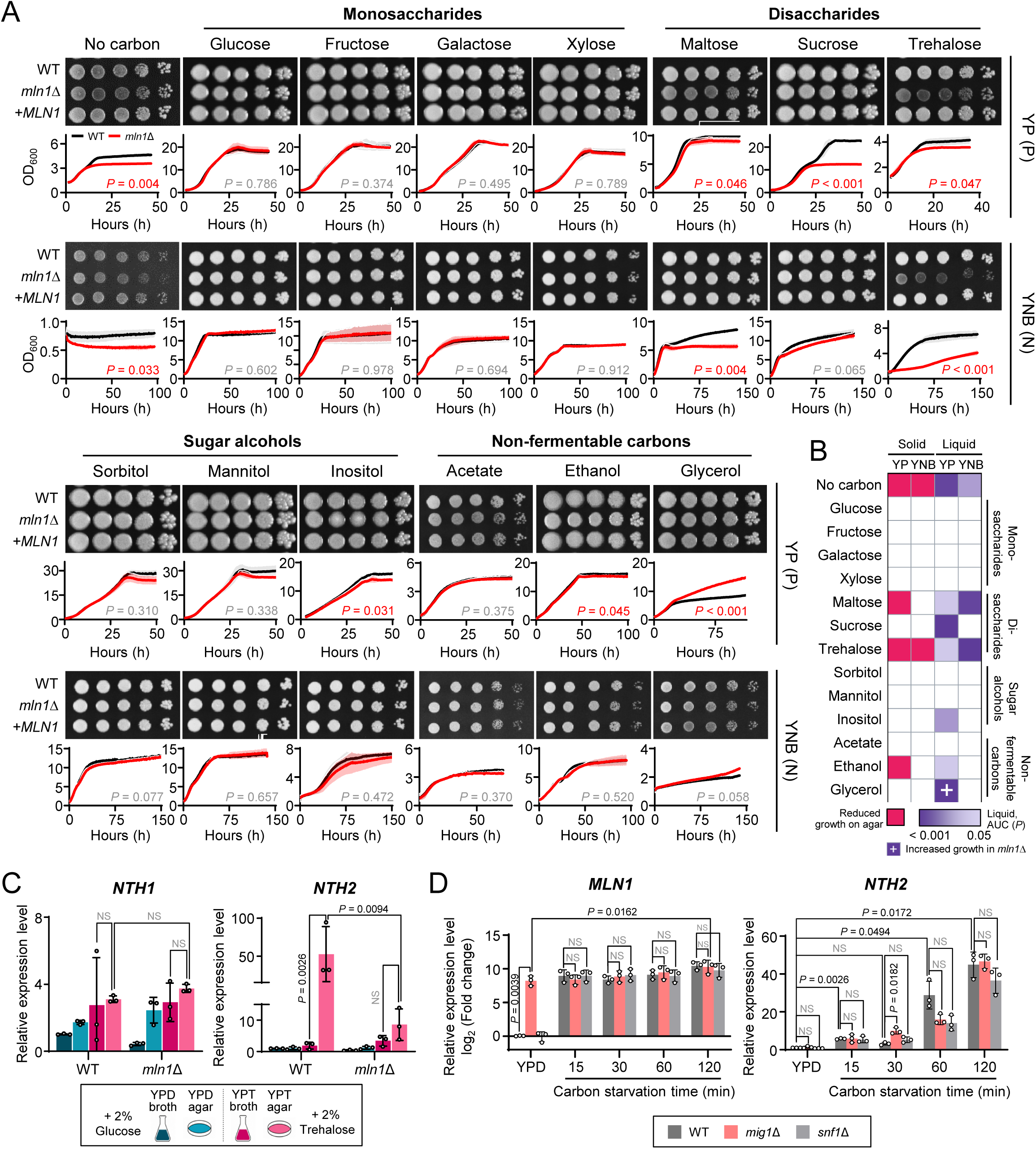
Mln1 supports condition-dependent utilization of selected non-preferred carbon sources in *C. neoformans*. (A) Carbon-source utilization of WT (H99), *mln1*Δ (YSB1172), and the complemented strain *mln1*Δ+*MLN1* (YSB11310) in YP- or YNB-based media without added carbon or supplemented with 2% of the indicated carbon source. Growth was assessed at 30°C by 10-fold serial-dilution spotting assays for all three strains and liquid OD_600_ growth curves for WT and *mln1*Δ. Growth curves represent the mean ± SD of three biological replicates. *P* values indicate comparisons between WT and *mln1*Δ based on the area under the curve (AUC), with significant values (*P* < 0.05) shown in red. (B) Heatmap summarizing Mln1-dependent growth phenotypes across carbon sources, media, and growth formats. Carbon sources follow the order in panel A. Pink boxes indicate reduced growth of *mln1*Δ on solid media, whereas color intensity for liquid cultures represents AUC-based *P* values, with darker colors indicating smaller *P* values. A plus sign (+) indicates significantly enhanced growth of *mln1*Δ relative to WT. (C) qRT-PCR analysis of *NTH1* and *NTH2* expression in WT and *mln1*Δ cells grown in YP medium supplemented with 2% glucose or trehalose under liquid and agar conditions. Transcript levels were normalized to *ACT1*. Bars represent the mean ± SD of three biological replicates. Statistical significance was assessed by three-way ANOVA with strain, carbon source, and growth format as factors, followed by Šídák’s multiple-comparisons test. (D) Time-course qRT-PCR analysis of *MLN1* and *NTH2* expression in WT, *mig1*Δ (YSB9271), and *snf1*Δ (YSB2372) following carbon starvation. Transcript levels were normalized to *ACT1* and expressed relative to the basal WT condition, with *MLN1* shown as log2 fold changes and *NTH2* as relative expression levels. Bars represent the mean ± SD of three biological replicates, with dots indicating individual replicates. Statistical significance was assessed by two-way ANOVA followed by Dunnett’s multiple-comparisons tests, comparing mutants with WT at each time point and starvation time points with basal YPD within each strain. Basal-to-120-min comparisons were significant in all three strains (WT, *P* = 0.0023; *mig1*Δ, *P* = 0.0162; *snf1*Δ, *P* = 0.0009), with only the *mig1*Δ comparison displayed for clarity. (C, D) Exact adjusted *P* values are shown for significant comparisons, whereas nonsignificant comparisons are indicated by gray “NS”.

The requirement for Mln1 was strongly influenced by both medium composition and growth format. In YP-based media, *mln1*Δ showed significant growth defects without added glucose and on several non-preferred carbon sources, including maltose, sucrose, trehalose, inositol, and ethanol. In contrast, glycerol supported significantly enhanced growth of *mln1*Δ relative to WT in liquid YP medium. In YNB-based media, the mutant defects were most evident under carbon starvation and on maltose and trehalose, whereas several other carbon sources produced weaker or no reproducible differences. These results indicate that Mln1-dependent carbon utilization is not simply determined by the chemical identity of the supplied carbon source, but also by the nutritional background and whether cells grow on a surface or in liquid culture.

Trehalose showed the most striking context-dependent behavior. On solid media, WT cells utilized trehalose efficiently, whereas *mln1*Δ showed a pronounced growth defect. In liquid culture, trehalose utilization differed between media: YP liquid supported little growth even in WT cells, whereas YNB liquid allowed more robust WT growth and revealed a strong defect in *mln1*Δ. Thus, Mln1 is required for trehalose utilization under conditions in which *C. neoformans* can effectively use trehalose as a carbon source. To summarize these complex condition-dependent phenotypes, we generated a heatmap integrating visible growth phenotypes on solid media with statistical differences in liquid growth assays across YP- and YNB-based conditions (Fig. 3B). This summary highlighted disaccharides, particularly trehalose and maltose, as carbon sources with recurrent Mln1-dependent growth defects.

Because WT cells utilized trehalose efficiently on YP agar supplemented with 2% trehalose (YPT agar) but showed little growth in the corresponding liquid medium (YPT broth), we asked whether this growth-format-dependent phenotype was associated with differential induction of trehalase genes. WT and *mln1*Δ cells were cultured in YP medium supplemented with either 2% glucose (YPD) or 2% trehalose (YPT) under liquid and agar conditions, and *NTH1* (neutral trehalase 1, CNAG_03785) and *NTH2* (neutral trehalase 2, CNAG_03525) expression was measured by quantitative reverse transcription PCR (qRT-PCR) (Fig. 3C). *NTH1* expression remained relatively stable across conditions. By contrast, *NTH2* was strongly induced in WT cells on YPT agar, whereas only weak induction was observed in YPT broth. Moreover, *NTH2* induction on YPT agar was markedly reduced in *mln1*Δ cells. These results suggest that surface-associated induction of *NTH2* is linked to efficient trehalose utilization in YP-based medium and that Mln1 is required for full activation of this response.

Because Mig1 (multicopy inhibitor of *GAL* gene expression 1) and Snf1 are well-established regulators of glucose repression and carbon source adaptation in fungi (19, 41, 42), we next asked whether they regulate *MLN1* expression in response to carbon availability. For Mig1, we constructed a strain in which the genomic region encoding residues 61-362, which overlaps the first C2H2 zinc-finger motif (residues 60-87) and encompasses the second motif (residues 88-112), was replaced by the *NAT* (nourseothricin N-acetyltransferase)-STM cassette (hereafter *mig1*Δ, Supplementary Information, Fig. S8F). To verify disruption of the *MIG1* zinc-finger region, we further examined *MIG1* transcripts using three qRT-PCR amplicons targeting the 5′ region, the deleted C2H2 zinc-finger region, and the 3′ region (Supplementary Information, Fig. S8G). In *mig1*Δ cells, the 5′ amplicon remained at WT-like levels, the amplicon within the deleted region was undetectable, and the 3′ amplicon remained detectable at a reduced level. These results confirmed the expected internal deletion of the *MIG1* DNA-binding region.

Having verified this allele, we grew WT, *mig1*Δ, and *snf1*Δ cells under glucose-replete conditions, shifted them to carbon starvation, and monitored *MLN1* transcript levels over time (Fig. 3D). Because basal *MLN1* expression in WT was very low, *MLN1* expression is presented as log_2_ fold changes relative to WT under the basal condition. Under basal conditions, *MLN1* expression was strongly derepressed in *mig1*Δ cells relative to WT, whereas *snf1*Δ had no significant effect. Notably, reanalysis of a previously published RNA-seq dataset generated using an independently constructed complete *mig1*Δ strain revealed similarly strong derepression of *MLN1* under glucose-containing conditions (20). Following carbon starvation, *MLN1* expression increased over time within each of the three strains, including *mig1*Δ, in which expression increased further from its already elevated basal level. When the strains were compared at the corresponding post-starvation time points, *MLN1* expression did not differ significantly among WT, *mig1*Δ, and *snf1*Δ cells despite the marked basal derepression observed in *mig1*Δ. Thus, although Mig1 is a major determinant of basal *MLN1* repression, the additional induction of *MLN1* during carbon starvation can occur in the absence of either Mig1 or Snf1 under the conditions tested.

To determine whether the strong basal derepression of *MLN1* in *mig1*Δ cells was sufficient to activate an Mln1-dependent downstream response, we also measured *NTH2* expression in WT, *mig1*Δ, and *snf1*Δ cells. Despite the strong derepression of *MLN1* in *mig1*Δ cells, *NTH2* transcript levels were not significantly different from those in WT cells under glucose-replete conditions (Fig. 3D). This result suggests that increased *MLN1* transcript abundance alone is insufficient to induce *NTH2* under glucose-replete conditions and that additional carbon-state-dependent regulatory signals may be required to activate Mln1-dependent downstream responses.

### Mln1 abundance and nuclear localization are differentially regulated by carbon availability

Because Mln1-dependent growth defects were most evident under carbon starvation and on selected non-preferred carbon sources, we next asked whether Mln1 responds to these conditions by altering its subcellular localization. For localization analysis, we generated a strain expressing Mln1-mRuby3 from a construct integrated at the *MLN1* locus and confirmed that the tagged protein retained function (Supplementary Information, Fig. S8B-D). We first examined Mln1 localization over time under nutrient-replete conditions, carbon starvation, nitrogen starvation, combined carbon and nitrogen starvation, glucose-free YP medium, and HMC (Fig. 4A and B).

**Figure 4.**
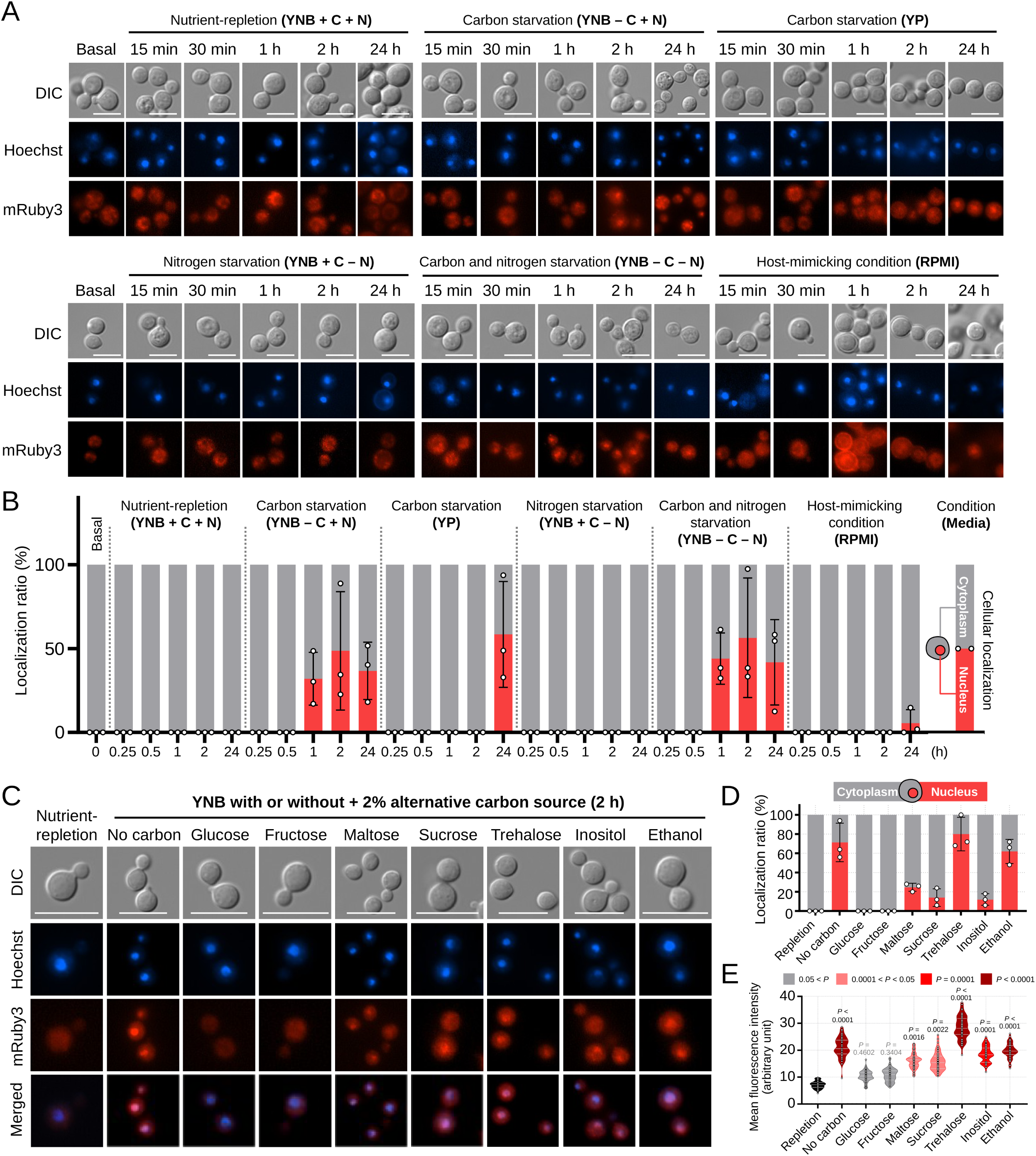
Mln1 abundance and nuclear localization respond differently to carbon availability. (A) Time-course fluorescence microscopy of Mln1-mRuby3 (YSB10818), expressed from a construct integrated at the *MLN1* locus, under nutrient-replete, carbon-depleted, nitrogen-depleted, or combined carbon/nitrogen-depleted YNB conditions, glucose-free YP medium, and HMC (RPMI 1640 with 10% FBS at 37°C under 5% CO_2_). Cells were transferred from YPD cultures grown at 30°C to the indicated media and sampled over time. DIC, Hoechst-stained nuclei, and mRuby3 fluorescence images are shown. Scale bars, 5 μm. (B) Quantification of cytoplasmic and nuclear-enriched Mln1-mRuby3 localization, classified by overlap with Hoechst-stained nuclei. At least 50 cells per condition and time point were analyzed in each of three independent biological replicates. Bars show mean percentages, and dots indicate biological replicates. (C) Representative DIC, Hoechst, mRuby3, and merged images after 2 h in YNB medium without added carbon or supplemented with the indicated carbon sources. Scale bars, 5 μm. (D) Quantification of Mln1-mRuby3 localization under the conditions in panel C, using the classification and presentation described in panel B. At least 50 cells per condition were analyzed in each of three independent biological replicates. (E) Single-cell quantification of mean whole-cell Mln1-mRuby3 fluorescence intensity under the conditions in panel C. Violin plots show individual-cell measurements from at least 50 cells per condition in each of three biological replicates. Statistical significance was assessed by nested one-way ANOVA, with cells nested within biological replicates, followed by Dunnett’s multiple-comparisons test against the nutrient-replete condition.

Under nutrient-replete YNB conditions, Mln1-mRuby3 showed predominantly diffuse cytoplasmic localization across all time points. In contrast, carbon starvation promoted nuclear accumulation of Mln1, with the strongest response observed at 2 h. Nuclear accumulation was also observed in glucose-free YP medium, but only after prolonged incubation (24 h). Nitrogen starvation alone did not induce detectable nuclear accumulation, indicating that transcriptional induction of *MLN1* by nitrogen limitation is not sufficient to drive Mln1 nuclear localization. Simultaneous carbon and nitrogen starvation also induced nuclear accumulation, further supporting carbon deprivation as the primary cue for Mln1 nuclear localization. Under HMC, which contains ∼0.2% glucose, Mln1 showed little nuclear accumulation, with only a modest increase at 24 h. Together, these data indicate that Mln1 nuclear localization is driven primarily by carbon limitation rather than by nitrogen starvation.

We next asked whether Mln1 nuclear accumulation reflects complete carbon depletion or whether it can also be triggered by specific non-preferred carbon sources. Based on the carbon-source utilization phenotypes, we examined Mln1 localization after 2 h in YNB medium lacking carbon or supplemented with individual carbon sources, including glucose and fructose as preferred or readily metabolized monosaccharide controls, maltose, sucrose, and trehalose as disaccharides, and inositol and ethanol as additional non-preferred carbon sources (Fig. 4C-E). Carbon starvation and trehalose induced the strongest nuclear accumulation, whereas ethanol also triggered robust nuclear enrichment despite producing a relatively modest growth phenotype (Fig. 4D). Maltose and sucrose produced intermediate responses, and inositol induced a weaker but detectable increase in nuclear localization. In contrast, glucose and fructose failed to induce substantial nuclear accumulation above the basal level. We next asked whether these localization changes were accompanied by changes in overall Mln1 abundance. Whole-cell Mln1-mRuby3 fluorescence increased under several non-preferred carbon conditions (Fig. 4E), consistent with increased Mln1 abundance. Inositol increased whole-cell Mln1-mRuby3 fluorescence but induced only modest nuclear accumulation (Fig. 4D and E), further indicating that Mln1 abundance and nuclear localization are not strictly coupled.

Together, these analyses show that Mln1 regulation involves separable changes in whole-cell fluorescence and carbon-dependent nuclear localization. The lack of nuclear accumulation during nitrogen starvation, a condition that strongly induces *MLN1*, further indicates that transcriptional induction alone is not sufficient to drive Mln1 nuclear localization. These findings identify subcellular localization as an additional layer of Mln1 regulation under carbon-limited and non-preferred carbon conditions.

### Mln1 regulates a carbon-state-dependent transcriptional program for alternative carbon utilization

Because Mln1 accumulated in the nucleus specifically under carbon-limited and selected non-preferred carbon conditions, we next sought to define the transcriptional program regulated by Mln1 across different nutrient conditions. We performed RNA sequencing (RNA-seq) analysis of WT and *mln1*Δ cells under nutrient-replete, carbon-starved, and nitrogen-starved conditions (Fig. 5A and Supplementary Information, Dataset S2). Principal-component analysis showed separation primarily by nutrient condition, with genotype-dependent separation most apparent under carbon starvation, whereas hierarchical clustering showed predominant nutrient-condition-associated clustering with intermingling of WT and *mln1*Δ replicates within conditions (Supplementary Information, Fig. S9A and S9B). In addition, comparison of WT cells under nutrient-replete and starvation conditions largely recapitulated the carbon- and nitrogen-responsive TF signatures identified in our qRT-PCR-based host-cue screen, although the nitrogen response of *ZNF2* was not reproduced in the RNA-seq dataset (Supplementary Information, Fig. S9C-F).

**Figure 5.**
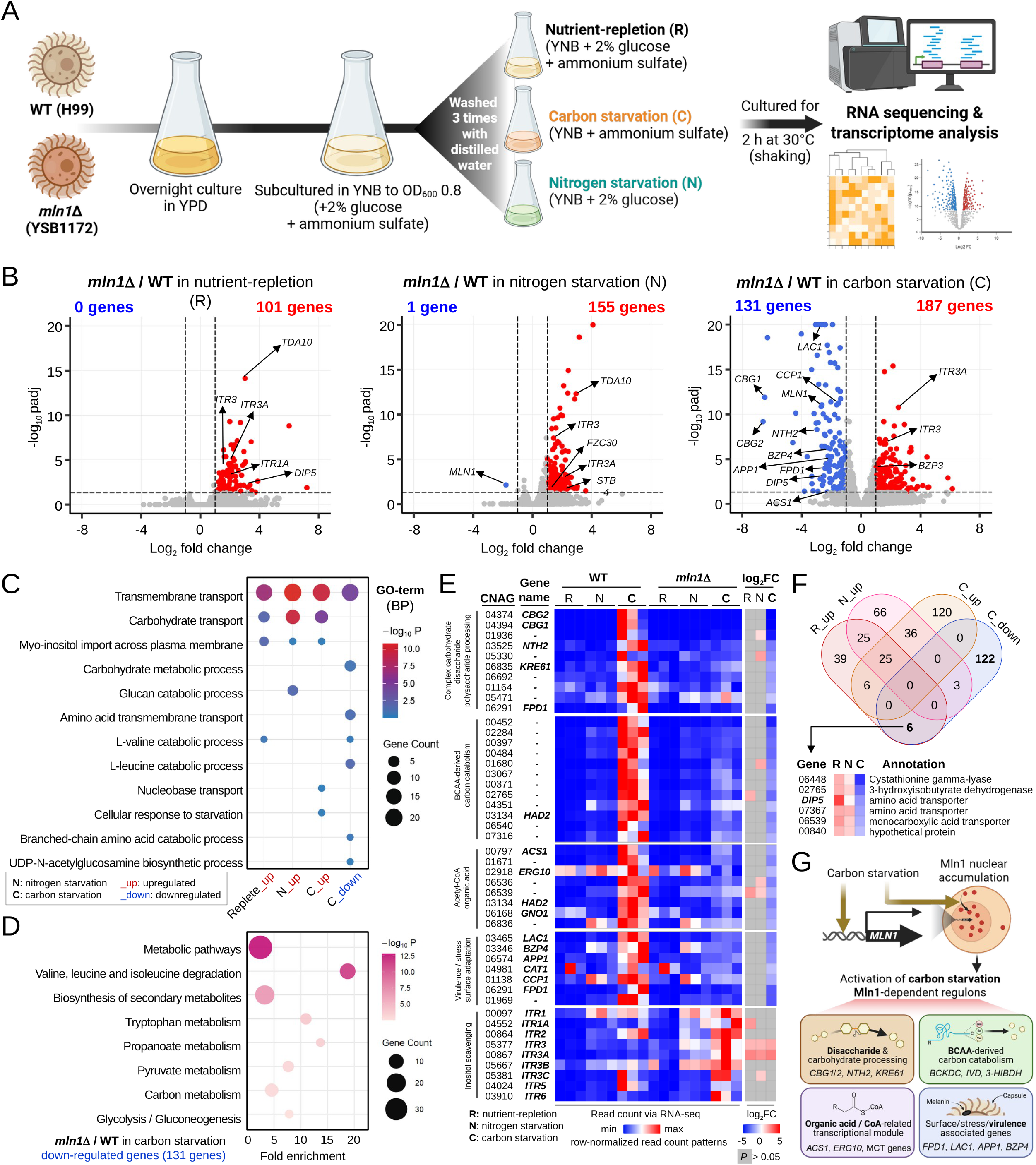
RNA-seq reveals a carbon-state-dependent transcriptional program controlled by Mln1. (A) Schematic overview of the RNA-seq experimental design. WT (H99) and *mln1*Δ (YSB1172) cells were grown in YNB medium containing 2% glucose and ammonium sulfate until OD_600_ reached 0.8, then transferred to nutrient-replete, carbon-starved, or nitrogen-starved conditions for 2 h. Glucose or ammonium sulfate was omitted to induce carbon or nitrogen starvation, respectively, and all media lacked amino acids. (B) Volcano plots showing differentially expressed genes (DEGs) in *mln1*Δ relative to WT under each nutrient condition. Red, blue, and gray dots indicate significantly upregulated, downregulated, and nonsignificant genes, respectively (|log_2_ fold change| > 1 and adjusted *P* < 0.05). DEG counts and representative genes are indicated. Full results are provided in Supplementary Information, Dataset S2. (C) Gene Ontology (GO) enrichment analysis of DEGs under each nutrient condition. Dot size indicates gene count, and color intensity indicates −log_10_ *P* value. (D) Top eight enriched KEGG pathways among genes downregulated in *mln1*Δ during carbon starvation. Dot size and color intensity represent gene count and −log_10_ *P* value, respectively. (E) Heatmap of row-wise z-scores calculated from raw read counts of selected genes grouped by functional category. Blue, white, and red indicate relatively low, intermediate, and high expression within each gene, respectively. The rightmost column shows log_2_ fold changes in *mln1*Δ relative to WT under each condition, with nonsignificant values (adjusted *P* ≥ 0.05) shown in gray. Genes belonging to multiple categories appear in each corresponding block. (F) Venn diagram showing overlap among genes upregulated under nutrient-replete (R_up), nitrogen-starved (N_up), and carbon-starved (C_up) conditions or downregulated under carbon starvation (C_down). The accompanying heatmap highlights representative genes with carbon-state-dependent expression patterns. (G) Summary of the Mln1-dependent carbon-starvation transcriptional program, including *MLN1* induction, Mln1 nuclear accumulation, and regulation of genes involved in carbohydrate processing, branched-chain amino acid (BCAA) catabolism, organic acid/CoA metabolism, and surface- or stress-associated functions. Schematics in A and G were created with BioRender (https://BioRender.com/45upvnl).

Deletion of *MLN1* produced a strongly condition-dependent transcriptional effect (Fig. 5B). Under nutrient-replete conditions, 101 genes were upregulated in *mln1*Δ cells, whereas no genes were significantly reduced. Basal *MLN1* expression in WT cells was extremely low under this condition, and its loss was therefore not detected as a significant downregulation in the RNA-seq analysis. Nitrogen starvation produced a similar pattern, with 155 genes upregulated and only *MLN1* itself significantly reduced, reflecting its strong induction in WT cells under nitrogen starvation. In contrast, carbon starvation revealed a broad bidirectional response, with 187 genes upregulated and 131 genes downregulated in *mln1*Δ cells. This pattern indicates that Mln1 supports a distinct transcriptional program specifically during carbon starvation.

Gene Ontology (GO) analysis of genes upregulated in *mln1*Δ cells revealed enrichment for transporter-related functions across nutrient-replete, carbon-starved, and nitrogen-starved conditions (Fig. 5C). Under nutrient-replete conditions, upregulated genes were associated with transmembrane and carbohydrate transport. Similar enrichment was observed during carbon and nitrogen starvation, including genes involved in carbohydrate and amino acid transport (Supplementary Information, Dataset S2). Among these transporter-related genes, two myo-inositol transporters, *ITR3* and *ITR3A*, were consistently upregulated in *mln1*Δ cells across all three nutrient conditions (Fig. 5E). Given the abundance of inositol in brain tissue and its importance during cryptococcal CNS infection (21–23), this transcriptional pattern may be relevant to CNS-associated adaptation.

In contrast, genes downregulated in *mln1*Δ cells during carbon starvation were enriched for branched-chain amino acid (BCAA) catabolism and central carbon metabolic pathways (Fig. 5C and D). *CBG2* and *CBG1*, which encode predicted β-glucuronidases (23), were among the most strongly downregulated genes, with log_2_ fold changes of approximately -6.6 and -6.5, respectively. *NTH2, KRE61*, α-glucosidase (CNAG_05471), α-glucoside symporters, sugar transporters, and enzymes involved in amino sugar metabolism were also reduced. Together, these transcriptional changes provide a potential molecular basis for the growth defects of *mln1*Δ on trehalose, maltose, sucrose, and other non-preferred carbon sources.

Notably, multiple enzymatic steps of BCAA catabolism were coordinately downregulated in *mln1*Δ during carbon starvation (Fig. 5E). These included components of the branched-chain α-keto acid dehydrogenase complex (BCKDC), isovaleryl-CoA dehydrogenase (IVD), 3-methylcrotonyl-CoA carboxylase, and 3-hydroxyisobutyrate dehydrogenase (3-HIBDH). This broader metabolic signature also extended to genes associated with organic acid and CoA-linked metabolism, including *ACS1, ERG10,* and monocarboxylate transporters (MCTs). Beyond these metabolic functions, the Mln1-dependent carbon-starvation gene set also included genes associated with cell-surface remodeling, host adaptation, and stress responses, including *FPD1, LAC1, BZP4, APP1, CAT1,* and *CCP1*.

Overlap analysis further showed that the set of genes downregulated during carbon starvation was largely distinct from genes upregulated in *mln1*Δ cells under nutrient-replete or nitrogen-starved conditions. Interestingly, six genes were upregulated in *mln1*Δ under nutrient-replete conditions but downregulated during carbon starvation (Fig. 5F). These included *DIP5*, another amino acid transporter, a monocarboxylic acid transporter (MCT), cystathionine gamma-lyase, 3-hydroxyisobutyrate dehydrogenase, and one uncharacterized protein, further illustrating that the transcriptional consequences of *MLN1* deletion depend strongly on carbon state. Collectively, these RNA-seq data show that Mln1 supports a carbon-state-dependent transcriptional program involving alternative carbon catabolism, trehalose metabolism, and broader metabolic and stress-associated functions during carbon starvation (Fig. 5G).

### Mln1 links trehalose homeostasis to carbon adaptation and acute heat-shock resistance

Our transcriptomic analysis revealed Mln1-dependent changes in genes involved in alternative carbon metabolism during carbon starvation, prompting us to examine whether these transcriptional changes were accompanied by alterations in intracellular metabolite abundance during carbon limitation. In parallel, the defects in trehalose utilization and *NTH2* expression observed above, together with the enhanced growth of *mln1*Δ at elevated temperature, led us to hypothesize that loss of *MLN1* may increase intracellular trehalose levels. We therefore examined trehalose abundance during both carbon starvation and thermal adaptation. For carbon-starvation profiling, cells were grown to exponential phase in YNB medium containing 2% glucose and ammonium sulfate, sampled under this nutrient-replete condition, and then transferred to YNB medium lacking glucose. For heat-stress profiling, cells grown in YPD at 30°C were sampled before transfer to 39°C. Intracellular metabolites were extracted from cell pellets collected at the indicated time points and analyzed by gas chromatography-mass spectrometry (GC-MS) (Fig. 6A).

**Figure 6.**
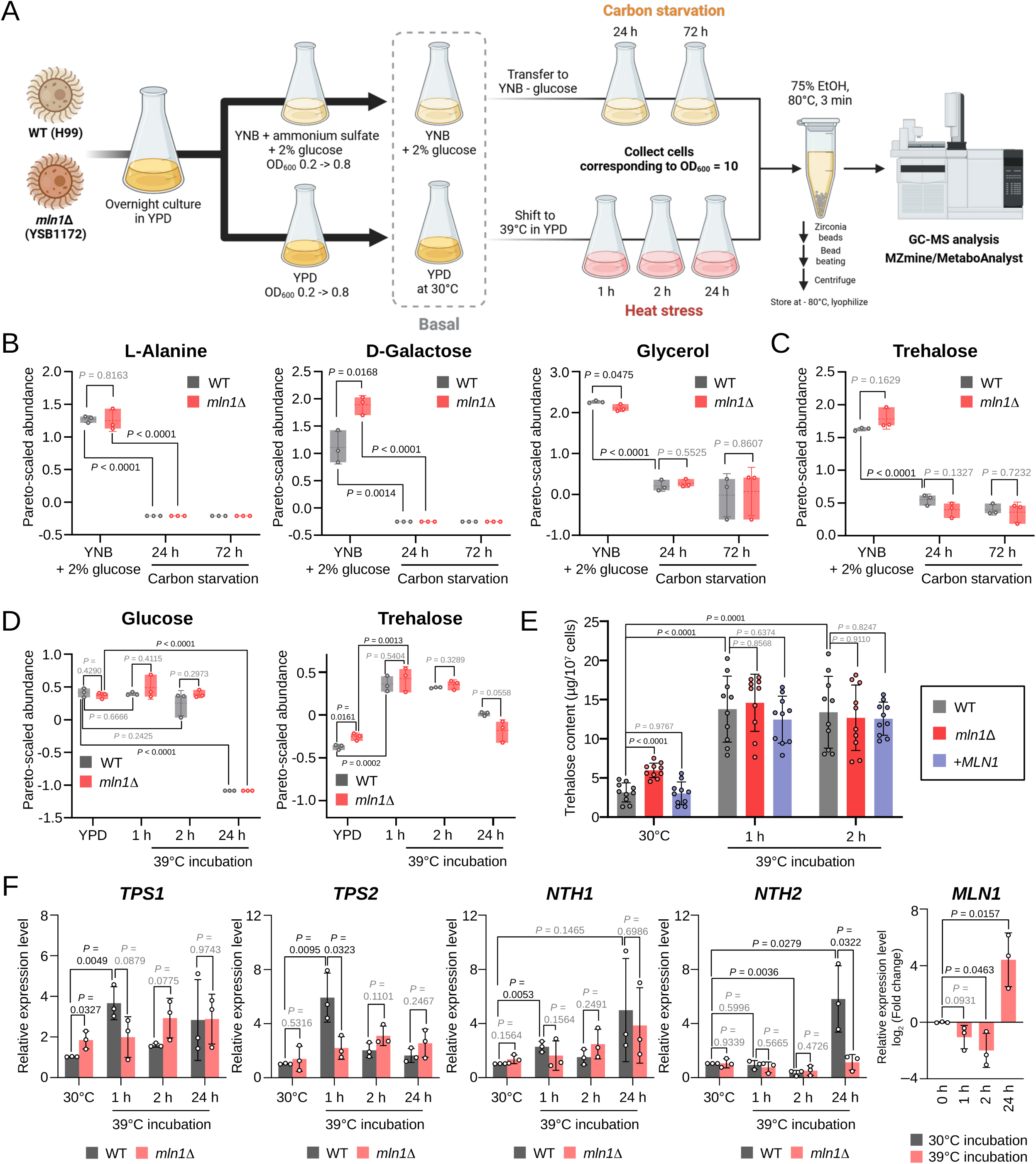
Loss of *MLN1* alters trehalose homeostasis. (A) Schematic overview of metabolite and transcriptional analyses in WT and *mln1*Δ cells under carbon starvation and elevated temperature. Cells grown in nutrient-replete YNB medium were transferred to carbon starvation, whereas cells grown in YPD at 30°C were shifted to 39°C. Intracellular metabolites were analyzed by GC-MS, with parallel heat-stress samples collected for qRT-PCR. The schematic was created with BioRender (https://BioRender.com/45upvnl). (B) Intracellular abundances of L-alanine, D-galactose, and glycerol before and during carbon starvation. (C) Intracellular trehalose abundance before and during prolonged carbon starvation. (D) Intracellular trehalose and glucose abundances before and after temperature upshift from 30°C to 39°C in YPD medium. (B–D) Metabolite abundances were normalized as described in Materials and Methods. Floating bars indicate minimum and maximum values, bars represent the mean ± SD of three biological replicates, and dots indicate individual replicates. Statistical significance was assessed using unpaired two-tailed Student’s *t*-tests for the indicated comparisons, with WT and *mln1*Δ compared at each time point in panel D. *P* values are shown in the panels. (E) Enzymatic quantification of intracellular trehalose in WT, *mln1*Δ, and the *MLN1*-complemented strain at 30°C and after 1 or 2 h at 39°C. Trehalose content was expressed as μg per 10^7^ cells. Bars represent the mean ± SD, and dots indicate biological replicates. Statistical significance was assessed by two-way ANOVA followed by Dunnett’s multiple-comparisons test against WT within each condition and Tukey’s test across temperature/time conditions within each strain. *P* values are shown above the corresponding brackets. (F) qRT-PCR analysis of *TPS1*, *TPS2*, *NTH1*, *NTH2*, and *MLN1* during incubation at 39°C. Transcript levels were normalized to *ACT1* and expressed relative to the WT basal condition, except *MLN1*, which is shown for WT only as log_2_ fold changes. Bars represent the mean ± SD of three biological replicates, with dots indicating individual replicates. Statistical significance was assessed using unpaired two-tailed Student’s *t*-tests, with *P* values shown in the panels.

Under nutrient-replete YNB conditions, L-alanine, D-galactose, and glycerol were readily detected in the intracellular metabolite pool. Following carbon starvation, L-alanine and D-galactose decreased to below the detection limit by 24 h and remained undetectable thereafter, whereas glycerol also decreased markedly but remained detectable (Fig. 6B). D-galactose abundance was higher in *mln1*Δ cells than in WT cells under nutrient-replete conditions, but this difference disappeared following carbon starvation (Fig. 6B). This increase may be associated with the enrichment of carbohydrate transport-related genes among those upregulated in *mln1*Δ cells under nutrient-replete conditions (Fig. 5C). In contrast to the marked depletion of L-alanine, D-galactose, and glycerol during carbon starvation, trehalose remained readily detectable throughout prolonged starvation (Fig. 6C). This persistence is consistent with its dual role as both a mobilizable carbon reserve and a stress-protective metabolite in fungi (29–31). However, intracellular trehalose abundance did not differ significantly between WT and *mln1*Δ cells during carbon starvation (Fig. 6C). Thus, despite the Mln1-dependent regulation of *NTH2*, we did not detect a significant effect of *MLN1* deletion on the bulk intracellular trehalose pool under carbon-starved conditions.

We next examined intracellular metabolites during adaptation to elevated temperature. Under basal YPD conditions before heat exposure, intracellular trehalose abundance was significantly higher in *mln1*Δ cells than in WT cells (Fig. 6D), revealing an altered basal trehalose state in the mutant. Following a shift to 39°C, trehalose abundance increased in both WT and *mln1*Δ cells at 1 h, whereas intracellular glucose did not show a comparable increase (Fig. 6D). Trehalose levels subsequently declined toward those observed under basal conditions by 24 h. The early increase in trehalose abundance is consistent with the well-established accumulation of trehalose during thermal stress in fungi (43). Given the protective function of trehalose under thermal stress, the elevated trehalose pool in *mln1*Δ cells prior to heat exposure suggested that altered trehalose homeostasis may contribute to their enhanced thermotolerance.

To independently validate the altered trehalose abundance observed in the metabolite analysis, intracellular trehalose content was quantified using an enzymatic trehalose assay in WT, *mln1*Δ, and the *MLN1*-complemented strain (Fig. 6E). Under basal conditions at 30°C, *mln1*Δ cells contained significantly more trehalose than WT cells, whereas trehalose content in the complemented strain did not differ significantly from that in WT. Following the shift to 39°C, trehalose content increased markedly in WT cells by 1 h and remained elevated at 2 h. In contrast to the basal condition, no significant differences were observed between *mln1*Δ and WT at either 1 h or 2 h, and the complemented strain likewise did not differ significantly from WT. These results independently confirm that loss of *MLN1* elevates the basal intracellular trehalose pool, while acute heat exposure induces trehalose accumulation across all three genotypes.

To determine whether altered trehalose abundance was accompanied by changes in trehalose-metabolic gene expression, parallel aliquots from the metabolite cultures were analyzed by qRT-PCR for *TPS1, TPS2, NTH1, NTH2,* and *MLN1* (Fig. 6F). Under basal conditions, *TPS1* expression was significantly higher in *mln1*Δ cells than in WT cells, consistent with the elevated basal trehalose abundance in the mutant. Following the shift to 39°C, *TPS1* and *TPS2* were rapidly induced in WT cells, peaking at 1 h and declining by 2 h, whereas both genes increased more gradually in *mln1*Δ cells and continued to rise through 2 h. These results indicate that loss of *MLN1* alters the early transcriptional dynamics of trehalose biosynthesis during thermal adaptation.

Trehalase expression also diverged during thermal adaptation: *NTH1* was transiently and significantly induced in WT cells at 1 h, whereas *NTH2* decreased after the temperature shift, reaching a significant reduction at 2 h, and then increased significantly by 24 h. *MLN1* showed a similar temporal shift, with transcript levels decreasing at 1 and 2 h, then increasing and reaching a level above the basal level at 24 h. Because both *NTH2* and *MLN1* are strongly induced by carbon limitation, the late increase in *NTH2*, together with the upward trend in *MLN1*, may reflect the emergence of nutrient limitation rather than a direct response to elevated temperature.

Because *mln1*Δ cells contained an elevated trehalose pool before heat exposure, we next asked whether this altered basal metabolic state was associated with increased resistance to acute lethal heat stress by exposing cells to 55°C for 15-30 min. Following heat shock, *mln1*Δ cells showed greater resistance than WT cells, whereas complementation of *MLN1* restored heat-shock sensitivity to WT levels (Supplementary Information, Fig. S10). Given the established role of trehalose in protecting yeast cells from severe heat stress (44), these findings are consistent with the possibility that the elevated basal trehalose pool in *mln1*Δ contributes to its enhanced heat-shock resistance.

Together, these results link Mln1 to two distinct aspects of trehalose biology. Mln1 contributes to basal trehalose homeostasis under glucose-replete YPD conditions and supports efficient growth when trehalose is used as an alternative carbon source, in parallel with Mln1-dependent induction of *NTH2*. Loss of *MLN1* is therefore associated with elevated basal trehalose accumulation under YPD conditions and altered regulation of trehalose biosynthetic genes, which may contribute to enhanced thermotolerance, while simultaneously impairing the efficient utilization of trehalose as a carbon source.

### *MLN1* deletion impairs host-associated fitness and titan cell formation

We next examined whether *MLN1* deletion alters fungal fitness during mammalian infection. C57BL/6 mice were intranasally infected with WT, *mln1*Δ, or the complemented strain and monitored for survival (Fig. 7A and B). Overall survival of mice infected with *mln1*Δ was not significantly different from that of mice infected with WT (Fig. 7B). In a separate cohort of infected mice, analysis of fungal burden in organs revealed lower fungal burden in the brain. The difference in lung fungal burden between WT- and *mln1*Δ-infected mice was not statistically significant, whereas brain fungal burden was significantly reduced in the mutant and restored to WT levels in the complemented strain (Fig. 7C). These findings indicate that Mln1 contributes to CNS-associated fungal fitness without a detectable effect on lung fungal burden at the sampled time point or on overall murine survival.

**Figure 7.**
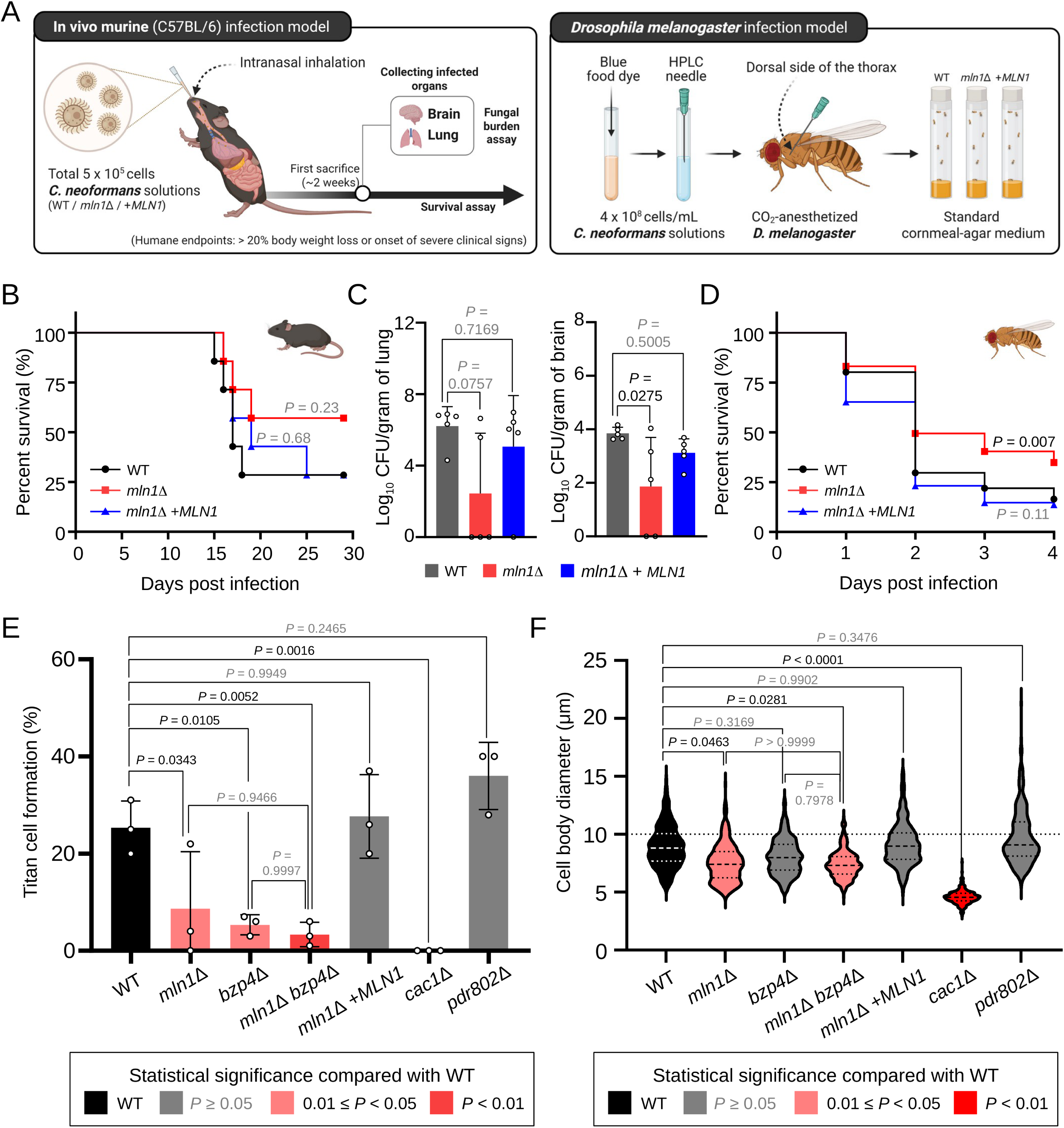
*MLN1* deletion impairs titan cell formation and host-associated fitness. (A) Schematic overview of murine and *Drosophila melanogaster* infection models. C57BL/6 mice were intranasally infected with 5 × 10^5^ cells of WT (H99), *mln1*Δ (YSB1172), or *mln1*Δ+*MLN1* (YSB11310) for survival and organ fungal burden analyses. Flies were injected with the same strains and maintained at 30°C for survival analysis. The schematic was created with BioRender (https://BioRender.com/45upvnl). (B) Survival of infected mice (n = 7 per strain), analyzed using the log-rank (Mantel-Cox) test. *P* values indicate comparisons with WT. (C) Lung and brain fungal burden at day 16 post-infection (n = 5 per strain), determined by colony-forming unit (CFU) counts. Dots represent individual mice, and bars indicate mean ± SD. Statistical significance was assessed by ordinary one-way ANOVA followed by Dunnett’s multiple-comparisons test against WT. (D) Survival of infected *Drosophila*, analyzed using the log-rank (Mantel-Cox) test, with *P* values indicating comparisons with WT. (E) Titan cell formation in WT, *mln1*Δ, *bzp4*Δ, *mln1*Δ *bzp4*Δ, *mln1*Δ+*MLN1*, *cac1*Δ, and *pdr802*Δ. Titan cells were defined by a cell body diameter ≥10 μm. Bars represent mean ± SD, and dots indicate biological replicates. Statistical significance was assessed by ordinary one-way ANOVA. (F) Distribution of cell body diameters under titan-inducing conditions. Violin plots show individual-cell measurements, and the dashed line indicates the 10 μm titan cell threshold. The *cac1*Δ and *pdr802*Δ strains served as controls for impaired and enhanced titan cell formation, respectively. Statistical significance was assessed by nested one-way ANOVA, with cells nested within biological replicates. (E, F) ANOVA was followed by Dunnett’s multiple-comparisons test against WT and Tukey’s test among non-WT strains. Exact adjusted *P* values are shown, with significant (*P* < 0.05) and nonsignificant values indicated in black and gray, respectively. Plot colors indicate significance relative to WT, as shown in the key.

Because loss of *MLN1* impairs carbon adaptation while enhancing growth at febrile temperatures, interpretation of its role in mammalian virulence could be confounded by the opposing effects of these phenotypes. We therefore assessed the virulence of the *mln1*Δ mutant in a *Drosophila melanogaster* infection model maintained at 30°C (Fig. 7A and D), where the mutant shows no growth advantage over WT (Supplementary Information, Fig. S7G). Flies infected with the *mln1*Δ mutant survived significantly longer than those infected with WT, whereas the complemented strain restored virulence to near-WT levels (Fig. 7D). These results show that loss of *MLN1* compromises virulence in a host model in which its high-temperature growth advantage is not evident.

We next asked whether Mln1 also contributes to virulence by regulating titan cell formation, as nutrient limitation and metabolic adaptation to nutrient-poor conditions are known to promote titan cell development (45), and this morphological transition promotes immune evasion and dissemination during infection (46). We therefore assessed titan cell formation using the previously established minimal-medium condition containing 15 mM glucose (45). Under titan-inducing conditions, *mln1*Δ cells produced fewer titan cells than WT, whereas complementation of *MLN1* restored titan cell formation to a WT-like level (Fig. 7E). Consistent with this defect, the cell body diameter distribution of *mln1*Δ cells was shifted toward smaller sizes, whereas the complemented strain resembled WT (Fig. 7F).

In addition to *MLN1*, *BZP4* is strongly induced by carbon starvation (Fig. 1D), and its expression was reduced in *mln1*Δ cells under carbon-starvation conditions (Fig. 5E). We therefore generated an *mln1*Δ *bzp4*Δ double mutant (Supplementary Information, Fig. S8E) to examine the individual and combined contributions of Mln1 and Bzp4 to titan cell formation (Fig. 7E and F). Similar to *mln1*Δ, the *bzp4*Δ mutant exhibited reduced titan cell formation, and the *mln1*Δ *bzp4*Δ double mutant also showed a pronounced defect. Neither titan cell formation frequency nor cell body diameter in the double mutant differed significantly from those of either single mutant (Fig. 7E and F). These results indicate that both Mln1 and Bzp4 contribute to titan cell development, but no additional effect of their combined loss was detected under these conditions.

Together with the reduced brain fungal burden in mice and the attenuated virulence observed in the *Drosophila* infection model, these findings indicate that Mln1 promotes host-associated fitness in a context-dependent manner, with contributions to titan cell formation and CNS-associated adaptation. Notably, the reduced brain fungal burden of the *mln1*Δ mutant is consistent with its diminished brain infectivity observed in our previous STM analysis (Supplementary Information, Fig. S1B), further supporting a role for Mln1 in CNS-associated fitness across these assays.

## Discussion

Our study reveals an unexpected relationship between carbon adaptation and thermal stress resistance in the human fungal pathogen *C. neoformans*. Through systematic analysis of host-responsive TFs, we identified Mln1 as a carbon-responsive regulator whose loss impairs alternative carbon utilization but enhances thermotolerance. This contrasting phenotype suggests that metabolic adaptation and stress resistance can be differentially affected by a single transcriptional regulator. Notably, the increased thermal resistance of *mln1*Δ emerged despite the absence of a prominent temperature-induced transcriptional response of *MLN1*, highlighting the importance of nutrient-responsive regulatory pathways in shaping thermal stress phenotypes.

The evolutionary position of Mln1 provides a useful context for understanding its role in carbon adaptation. Mln1 is a fungal-restricted bHLH TF with no clear ortholog in *S. cerevisiae*, although Rtg3 shows the closest sequence similarity among budding-yeast proteins (47, 48). In *S. cerevisiae*, Rtg3 forms a heterodimer with Rtg1 and mediates mitochondrial retrograde (RTG) signaling, linking mitochondrial status to transcriptional control of genes involved in the tricarboxylic acid (TCA) cycle and nitrogen metabolism (49, 50). Several features distinguish Mln1 from the canonical *S. cerevisiae* RTG pathway, as *C. neoformans* lacks a clear Rtg1 ortholog and deletion of *MLN1* did not substantially affect canonical RTG-responsive genes such as *ACO1*, *IDH1,* and *IDH2* under the nutrient conditions examined. A recent cross-species regulatory-network comparison further separated the two regulators, associating Mln1 primarily with carbohydrate transport, whereas Rtg3 was linked more strongly to TCA-cycle, amino acid, and nitrogen metabolism (48). Such functional diversification is also evident in other fungi. In *Candida albicans*, the Rtg1–Rtg3 complex regulates a distinct set of target genes compared with S. cerevisiae, whereas in *Komagataella phaffii*, Rtg1 partners with the divergent Rtg3-related protein RtgX to regulate carbon metabolism at both transcriptional and post-transcriptional levels (51, 52). Consistent with this cross-species diversification, our carbon-starvation transcriptome places Mln1 within a similarly distinct program centered on carbohydrate processing and alternative carbon catabolism, raising the possibility that Mln1 represents a divergent metabolic regulator with a role in carbon adaptation in *C. neoformans*.

The upstream regulation of *MLN1* further distinguishes basal glucose repression from the response to carbon limitation. In budding yeast, Snf1-dependent phosphorylation of Mig1 relieves glucose repression (41). In *C. neoformans*, the strong basal derepression of *MLN1* in *mig1*Δ cells identifies Mig1-dependent regulation as a major mechanism maintaining low *MLN1* expression under glucose-replete conditions, a relationship independently reproduced in a published *mig1*Δ RNA-seq dataset (20). Carbon starvation, however, induced *MLN1* in WT and *snf1*Δ cells and further increased its expression in *mig1*Δ cells despite the strong basal derepression caused by loss of Mig1. These results indicate that Snf1 is dispensable for starvation-induced *MLN1* expression under the conditions tested and that carbon starvation provides a regulatory input that remains active even when Mig1-dependent basal repression is absent. The downstream response further indicates that *MLN1* transcript abundance alone is insufficient to determine Mln1 activity. Despite strong basal *MLN1* derepression in glucose-grown *mig1*Δ cells, *NTH2* expression remained low. Likewise, nitrogen starvation strongly induced *MLN1* transcription without promoting Mln1 nuclear accumulation, whereas carbon starvation induced both *MLN1* expression and nuclear accumulation. Together, these observations support a layered regulatory model in which nutrient conditions control *MLN1* transcription, while carbon status provides an additional level of regulation through Mln1 nuclear localization.

The pronounced dependence of *CBG1* and *CBG2* on Mln1 provides a prominent example of the carbon-processing program controlled by Mln1. Both genes were strongly induced by carbon starvation in WT cells, whereas this response was markedly reduced in *mln1*Δ, placing *CBG1* and *CBG2* among the most strongly affected transcripts in the mutant. *CBG1* and *CBG2* encode predicted β-glucuronidases and are also induced by inositol together with *MIO1* and *MIO2* (23). Fungal β-glucuronidases can release glucuronic acid residues from complex glycans (53), suggesting that Mln1-dependent regulation of *CBG1* and *CBG2* may extend carbon adaptation beyond the utilization of readily available sugars.

Mln1-dependent carbon adaptation was not limited to carbohydrate-processing genes. Transcripts encoding multiple steps of BCAA catabolism were coordinately reduced in *mln1*Δ, together with *ACS1*, *ERG10*, and other genes involved in CoA-linked metabolism. Because BCAA degradation generates CoA-linked intermediates that feed into central carbon metabolism, these coordinated transcriptional changes suggest that Mln1 may support the utilization of both carbohydrate- and amino-acid-derived carbon during carbon limitation. In contrast, transporter-related genes were broadly increased in *mln1*Δ across nutrient conditions rather than specifically during carbon starvation. This pattern may reflect an indirect response to impaired carbon metabolism rather than direct negative regulation by Mln1. Mln1 thus appears to support the downstream processing of alternative carbon sources when preferred carbon becomes limiting.

Trehalose provides a functional connection between the Mln1-dependent transcriptional program and the utilization of an alternative carbon source. In YP medium, strong *NTH2* induction accompanied efficient trehalose-supported growth on agar, whereas both responses were weak in broth. Loss of *MLN1* reduced *NTH2* induction and impaired growth on trehalose agar, linking Mln1-dependent *NTH2* expression to conditions that support efficient trehalose utilization. Previous work in the JEC20/JEC21 background predicted Nth1 to be cytosolic and Nth2 to be secreted and showed overlapping contributions of the two trehalases to growth on trehalose (54). The predicted extracellular localization of Nth2 is particularly relevant to the utilization of supplied trehalose, because extracellular hydrolysis would make glucose available for uptake. Interestingly, acid-induced autolysis of *C. neoformans* has been shown to release trehalose into the extracellular environment (55), suggesting that autolyzing neighboring cells could provide an additional local source of trehalose for Nth2-dependent utilization. Because this release was demonstrated under acidic stationary-phase conditions, whether the same process contributes under carbon-limited conditions remains to be determined. The strong growth-format dependence of *NTH2* induction therefore provides a plausible explanation for why trehalose utilization differs between surface and liquid growth. These findings place Mln1 upstream of an *NTH2*-associated response that supports the utilization of extracellular trehalose.

In contrast to its role in the utilization of extracellular trehalose, Mln1 also influences intracellular trehalose homeostasis before heat exposure. Before heat exposure, *mln1*Δ cells contained more trehalose than WT cells and showed increased basal *TPS1* expression, indicating that loss of Mln1 shifts trehalose homeostasis toward accumulation under glucose-replete conditions. This pre-existing difference may be relevant to acute heat stress, as trehalose protects proteins from heat-induced damage in yeast (29, 44), and Tps1- and Tps2-dependent trehalose biosynthesis is required for cryptococcal growth at mammalian temperature (31). The two trehalase genes also showed distinct thermal responses. In WT cells, *NTH1* was transiently induced after the temperature shift, whereas *NTH2* decreased during the same early period. Heat-responsive induction of *NTH1* has also been reported in *S. cerevisiae* (56), whereas the carbon- and trehalose-responsive behavior of *NTH2* is more consistent with a role in nutrient utilization. Elevated basal trehalose may therefore contribute to the enhanced thermal resistance of *mln1*Δ cells, together with other Mln1-dependent metabolic changes.

The infection phenotypes further indicate that the contribution of Mln1 depends on host context. In the murine model, *MLN1* deletion reduced brain fungal burden without significantly affecting lung burden at the sampled time point, and no significant difference in host survival was observed. Notably, this reduction in brain fungal burden is consistent with the reduced brain infectivity of the *mln1*Δ observed in the previously published STM screen used for our initial TF selection (Supplementary Information, Fig. S1B). At the same time, the enhanced resistance of *mln1*Δ cells at 39°C suggests that the mutant may retain a relative advantage over WT at febrile temperatures, potentially influencing its behavior in the mammalian host. This consideration is relevant to the *Drosophila* model, in which *mln1*Δ exhibited attenuated virulence at 30°C, a temperature at which it shows no growth advantage over WT. This result indicates that the contribution of Mln1 to pathogenic fitness extends beyond its role in thermal adaptation. These models support a context-dependent contribution of Mln1 to pathogenic fitness, although differences in host biology and inoculation route prevent the effects of temperature from being isolated.

Several observations connect this CNS phenotype to the nutrient-responsive functions of Mln1. *MLN1* is induced in cryptococcal cells recovered from human cerebrospinal fluid and after ingestion by amoebae and macrophages (57, 58), consistent with its activation under diverse nutrient- and stress-associated conditions encountered during host and predator interactions. The brain is also enriched in inositol, which promotes cryptococcal BBB traversal and induces a transcriptional program that includes inositol transporters, *MIO1*/*MIO2*, and *CBG1*/*CBG2* (23). In carbon-starved *mln1*Δ cells, *CBG1* and *CBG2* expression was strongly reduced despite increased expression of several inositol transporter genes. This pattern suggests that nutrient uptake alone may be insufficient when downstream carbon-processing responses are impaired. Such a defect could be particularly relevant in the inositol-rich CNS, where efficient inositol acquisition and catabolism have been linked to cryptococcal virulence (22, 23).

Integrating these findings, we propose that Mln1 links carbon availability to the balance between nutrient utilization and trehalose-associated thermoprotection (Fig. 8). Under glucose-replete conditions, Mig1-dependent regulation maintains low basal *MLN1* expression and Mln1 remains predominantly outside the nucleus, whereas carbon limitation induces *MLN1* expression and promotes Mln1 nuclear accumulation. This carbon-responsive regulation supports the utilization of trehalose and other non-preferred carbon sources. Loss of *MLN1*, however, produces contrasting effects across environmental conditions. It impairs adaptation to carbon limitation and alternative carbon sources, yet increases basal trehalose abundance under glucose-replete conditions and enhances resistance to acute heat stress. These opposing phenotypes illustrate how loss of a carbon-responsive regulator can compromise nutrient adaptation while simultaneously increasing resistance to thermal stress. The emergence of this thermal phenotype from the carbon-responsive rather than the temperature-responsive module further highlights the contribution of metabolic state to thermotolerance. Nevertheless, several mechanistic questions remain unresolved. The association between elevated basal trehalose and enhanced heat resistance does not establish that trehalose accumulation causes the thermal phenotype. Direct promoter binding by Mln1 or Mig1 was not tested, and transcript abundance does not establish metabolic flux. In addition, lower brain fungal burden after intranasal infection could reflect altered dissemination, brain entry, or persistence within the brain. Resolving these alternatives will require additional targeted genetic and infection experiments in future studies. In conclusion, these findings place Mln1 at the interface between carbon adaptation, trehalose homeostasis, and host-associated fitness of *C. neoformans*.

**Figure 8.**
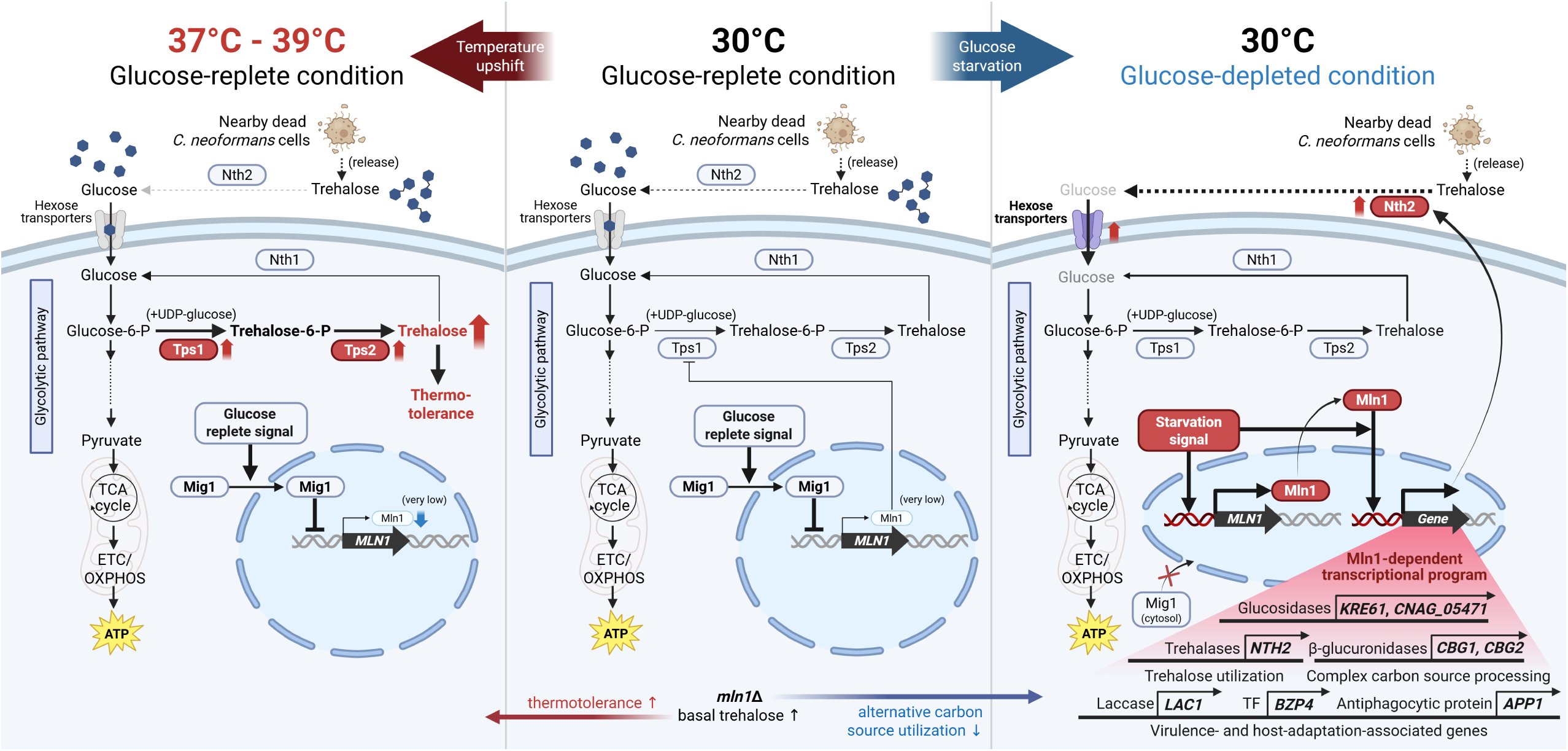
Proposed model for Mln1-mediated coordination of alternative carbon utilization and trehalose-associated thermoprotection in *C. neoformans*. The model summarizes Mln1 regulation under glucose-replete conditions at 30°C and elevated temperature and under glucose depletion at 30°C. Under glucose-replete conditions, Mig1-dependent regulation represses *MLN1*, and Mln1 remains predominantly outside the nucleus. Glucose depletion induces *MLN1* expression and Mln1 nuclear accumulation, supporting an alternative carbon-utilization program involving *NTH2*. Nth2 is predicted to act extracellularly, where trehalose hydrolysis may provide glucose for uptake. At elevated temperature, trehalose biosynthetic genes are induced and intracellular trehalose accumulates. The bottom panel summarizes *mln1*Δ phenotypes, including elevated basal intracellular trehalose under glucose-replete YPD conditions, altered *TPS1* and *TPS2* expression, and enhanced thermotolerance and acute heat-shock resistance. These findings suggest distinct roles for Mln1 in extracellular trehalose utilization and intracellular trehalose homeostasis, with a potential contribution of the latter to thermal protection. Arrows indicate regulatory or metabolic relationships, blunt-ended lines indicate inhibition, and dashed lines indicate proposed or indirect relationships. Red arrows and highlights indicate increased expression, abundance, or activation, gray components indicate decreases relative to the 30°C glucose-replete condition, and red crosses indicate inhibition. The schematic was created with BioRender (https://BioRender.com/45upvnl).

## Materials and Methods

### Ethics statement

Animal care and all experiments were conducted in accordance with the ethical guidelines of the Institutional Animal Care and Use Committee (IACUC) of AmtixBio. The AmtixBio IACUC approved all the vertebrate studies (ATBIACUC2023-07-21). All efforts were made to minimize animal suffering and to reduce the number of animals used.

### Media and growth conditions

*C. neoformans* strains were grown at 30°C in yeast extract-peptone-dextrose (YPD) medium. To mimic host conditions in vitro, we used the following medium and culture conditions: RPMI 1640 cell culture medium supplemented with 10% fetal bovine serum and incubation at 37°C in a 5% CO_2_ incubator (33). To further dissect medium-dependent factors, 9 different media conditions were used (Supplementary Information, Fig. S2A and Table S3). Cultures in each medium were incubated at 30°C, 37°C, or 37°C under 5% CO_2_. For temperature-upshift experiments, the WT and TF deletion mutants (*bzp4*Δ, *fzc30*Δ, *fzc39*Δ, *hob5*Δ, *pdr802*Δ, and *znf2*Δ) were inoculated at an optical density at 600 nm (OD_600_) of 0.2 and cultured at 30°C until the OD_600_ reached approximately 0.8. An aliquot of each culture was harvested as the basal sample. The remaining culture in the same flask was then shifted to 39°C and incubated for 1 h before harvesting the temperature-upshift sample. Three independent biological replicates were performed for each strain. Expression of candidate TFs was quantified by qRT-PCR following incubation under host-mimicking conditions or individual host-associated cues, whereas selected genes were further examined during carbon starvation and thermal adaptation, using *ACT1* as the reference gene (Supplementary Information, Materials and Methods). Mutant strains were constructed and validated by diagnostic PCR and Southern blot analysis (Supplementary Information, Materials and Methods).

### Selection of 24 pathogenicity-related TFs

To identify TFs associated with cryptococcal pathogenicity, data from two previously reported studies were utilized as references (32, 33). Initially, we analyzed existing in vivo transcriptional data obtained using NanoString methodology, as previously described (59). Briefly, this approach quantified the expression of 180 TFs across four infected tissues at multiple time points post-infection: early (day 3), intermediate (day 7), and late (days 14 and 21) stages. The in vivo expression of each TF was compared to its basal in vitro expression (YPD medium at 30°C). Expression data were normalized, converted into log_2_-based fold changes, and subsequently clustered using Morpheus software (http://software.broadinstitute.org/morpheus), developed by the Broad Institute (Supplementary Information, Fig. S1A; data were reorganized based on the original results from a previously published paper (33)). Given that lungs and the brain represent critical infection sites in cryptococcosis, TFs exhibiting at least a 2-fold increase in expression within both these organs were selected. Additionally, we incorporated data from previously conducted signature-tagged mutagenesis (STM)-based murine infection assays (32, 33). STM infectivity scores from intranasal (IN), intravenous (IV), and intracerebroventricular (ICV) infections were compiled from published datasets. Among the expression-filtered candidates, TFs whose mutants had an STM score ≤ -2 in at least one infection model were retained. Based on these criteria, we ultimately selected 24 TFs potentially linked to pathogenicity and regulated in vivo.

### Quantitative growth and nitrogen-source utilization assays

WT and mutant strains were cultured overnight in YPD medium at 30°C. To examine thermotolerance, 10-fold serial dilutions were spotted onto YPD agar and incubated at 30°C or 39°C for 6 days. For quantitative growth analysis, OD₆₀₀ was monitored continuously at 37°C, 38°C, or 39°C using a multi-channel bioreactor (RTS-8, Biosan). For the 39°C growth analysis of pairwise double-deletion mutants and their corresponding WT and single-deletion controls (Fig. 2B), cultures were inoculated at an initial OD_600_ of 2.0. Nitrogen source utilization was assessed in YNB medium lacking amino acids and ammonium sulfate, or supplemented with 5 g/L ammonium sulfate, a 1× amino acid mixture, or both (Supplementary Information, Materials and Methods). All growth assays were independently performed with three biological replicates. Chemical susceptibility was evaluated by serial-dilution spotting on drug-containing media (Supplementary Information, Materials and Methods).

### Growth curve analysis and carbon source utilization assay

To assess growth kinetics, each *C. neoformans* strain was cultured in 50 mL of YPD broth at 30°C for 16 h. Cells were harvested and resuspended in 10 mL of the indicated medium at an OD_600_ of 0.2, except for YNB without an added carbon source, for which the initial OD_600_ was 0.8. Growth was monitored every 30 min using a multi-channel bioreactor (RTS-8, Biosan) equipped with a reverse-spin mixing system at 30°C. For carbon source utilization assays, pre-cultured cells were resuspended in 10 mL of YP- or YNB-based medium without an added carbon source or supplemented with 2% glucose, fructose, galactose, xylose, maltose, sucrose, trehalose, sodium acetate, inositol, sorbitol, mannitol, glycerol, or ethanol. For carbon source utilization assays on solid medium, cells were washed three times with distilled water, resuspended in sterile water, serially diluted, and 3 μL of each dilution was spotted onto YP or YNB agar plates supplemented with 2% of the indicated carbon source. Plates were incubated at 30°C and colony growth was documented after four days. Brightness and contrast were adjusted uniformly across all images. To summarize the carbon-source growth phenotypes shown in Fig 3B, solid-medium phenotypes were scored qualitatively by visual comparison of serial-dilution growth between WT and *mln1*Δ cells, with conditions showing visible growth impairment indicated in pink. For liquid-culture conditions, heatmap values were based on the *P* values obtained from unpaired two-tailed Student’s *t*-tests comparing area-under-the-curve (AUC) values from three biological replicates of WT and *mln1*Δ, using the same statistical analysis as that applied to the growth curves in Fig 3A. Smaller *P* values were represented by greater color intensity. Conditions in which *mln1*Δ showed significantly enhanced growth relative to WT were indicated by a plus sign (+).

### Imaging of mRuby3-tagged proteins under different nutrient and host-mimicking conditions

Strains expressing mRuby3-tagged fusion proteins were initially cultured overnight in YPD medium at 30°C. Subsequently, cells were transferred to fresh YPD broth and cultured further until reaching an OD_600_ of 0.8. Cultured cells were harvested, washed three times with distilled water, and transferred into various experimental media for further analysis. To examine the effects of different nutrient conditions, cells were incubated in defined YNB-based media with or without 2% glucose and with or without ammonium sulfate. Additionally, cells were incubated in yeast extract-peptone (YP) medium lacking glucose, and RPMI 1640 medium supplemented with 10% FBS. All cultures were incubated at 30°C, except for RPMI 1640 medium, which was incubated at 37°C with 5% CO_2_ to simulate HMC. Cells were collected at indicated time points and fixed using 4% paraformaldehyde supplemented with 3.4% sucrose for 15 min. For individual carbon-source experiments, cells were transferred to YNB medium lacking an added carbon source or supplemented with 2% glucose, fructose, maltose, sucrose, trehalose, inositol, or ethanol and incubated for 2 h at 30°C before fixation. After fixation, nuclei were stained with 10 μg/mL Hoechst 33342 (Thermo Fisher) for 30 min in the dark. Prepared samples were examined and imaged using DIC and fluorescence microscopy (Nikon Eclipse, Tokyo, Japan) equipped with a digital camera (DS-Qi2). Fluorescence images were analyzed using NIS-Elements software (Nikon). Nuclei were identified based on Hoechst 33342 fluorescence. For quantification of Mln1-mRuby3 subcellular localization (Fig. 4B and D), cells showing visible enrichment of the mRuby3 signal overlapping the Hoechst-defined nuclear region were classified as nuclear-enriched, whereas cells lacking such nuclear enrichment were classified as predominantly cytoplasmic. For single-cell quantification of whole-cell Mln1-mRuby3 fluorescence (Fig. 4E), mean mRuby3 fluorescence intensity across the whole-cell area was quantified using NIS-Elements. All images used for fluorescence quantification were acquired using identical microscope settings with a 2-s exposure in the TxRED channel. Background fluorescence was measured under the same imaging conditions and subtracted from the fluorescence intensity measured for each cell. For both localization and fluorescence-intensity analyses, at least 50 cells per condition were analyzed in each of three independent biological replicates, with each time point analyzed separately for the time-course experiment.

### Total RNA isolation

WT and *mln1*Δ (YSB1172) strains were inoculated into 50 mL of YPD broth and cultured overnight at 30°C with shaking. Cells were washed three times with sterile distilled water and subcultured in 120 mL of fresh YNB-based basal medium until the OD_600_ reached approximately 0.8. The basal medium consisted of yeast nitrogen base containing ammonium sulfate but lacking amino acids, supplemented with 2% glucose. Cells were then harvested, washed three times with sterile distilled water, and divided equally into three conditions: nutrient repletion medium, carbon starvation medium, and nitrogen starvation medium. The nutrient repletion medium consisted of YNB supplemented with 2% glucose and ammonium sulfate but without amino acids. For carbon starvation, glucose was omitted while ammonium sulfate was retained, whereas for nitrogen starvation, ammonium sulfate was omitted while glucose was retained. Amino acids were absent from all three media to prevent their utilization as alternative carbon or nitrogen sources. Cells were incubated for an additional 2 h at 30°C with shaking, rapidly harvested, snap-frozen in liquid nitrogen, and stored at -80°C until RNA extraction. Total RNA was isolated by TRIzol extraction (Easy-BLUE, iNtRON).

### RNA-seq library preparation and sequencing

Total RNA samples were submitted to Macrogen Inc. (Seoul, Republic of Korea) for RNA-seq library preparation and sequencing. Sequencing libraries were prepared using a TruSeq RNA library preparation kit (Illumina, San Diego, CA, USA) according to the manufacturer’s instructions and sequenced on an Illumina HiSeq X Ten platform to generate 150-bp paired-end reads. Three independent biological replicates were sequenced for each combination of strain and nutrient condition.

### Transcriptome analysis

RNA-seq reads were aligned to the Cryptococcus neoformans H99 reference genome using HISAT2 v2.2.1. Gene-level expression data were analyzed using DESeq2 v1.24 (60). Differentially expressed genes (DEGs) were identified using thresholds of |log_2_FC| > 1 and adjusted *P* < 0.05, and genes upregulated or downregulated in *mln1*Δ cells were analyzed separately under each nutrient condition. Functional enrichment analysis was performed using DAVID Bioinformatics Resources (61), with all annotated genes of *C. neoformans* H99 as the background gene set. Enrichment significance was assessed using the EASE score with a threshold of 0.1 and a minimum gene count of 2. Benjamini–Hochberg-adjusted values and full enrichment results are provided in Supplementary Information, Dataset S2. Gene-level raw read counts were standardized by row using Z-scores to visualize relative expression patterns (Supplementary Information, Materials and Methods).

### Preparation of metabolite samples for GC-MS analysis

For metabolite profiling under carbon starvation, WT and *mln1*Δ cells were cultured in YNB medium containing 2% glucose and ammonium sulfate at 30°C until the OD_600_ reached approximately 0.8. Basal samples were collected before transfer to YNB medium lacking glucose, and additional samples were collected at the indicated time points after carbon starvation. For heat-stress profiling, cells grown in YPD at 30°C were sampled before transfer to 39°C and after 1, 2, and 24 h of incubation. Cell pellets corresponding to a total OD_600_ of 10 were collected at each time point. Intracellular metabolites were extracted and analyzed by GC-MS (Supplementary Information, Materials and Methods).

### Enzymatic quantification of intracellular trehalose

WT, *mln1*Δ, and the *MLN1*-complemented strain were cultured in YPD at 30°C and collected before or after incubation at 39°C for 1 or 2 h. Cells corresponding to 5 OD_600_ units (approximately 8.5 × 10^7^ cells) were harvested, and intracellular metabolites were extracted using boiling distilled water. Trehalose content was quantified using a Trehalose Assay Kit (Megazyme, K-TREH) by measuring the trehalase-dependent change in absorbance at 340 nm relative to a glucose standard, with correction for blank absorbance. Trehalose amounts were normalized to cell number and expressed as μg per 10^7^ cells (Supplementary Information, Materials and Methods).

### Titan cell formation assay

Titan cell formation was induced in vitro using the strains listed in this section. Overnight cultures were grown in 2 mL of YPD medium at 30°C. Cells were harvested by centrifugation at 3,000 rpm for 3 min, and the supernatant was discarded. Cell pellets were washed twice with 1 mL of minimal medium (MM), followed by centrifugation and removal of the supernatant. The MM used consisted of 15 mM glucose, 10 mM MgSO_4_, 29.4 mM KH_2_PO_4_, 13 mM glycine, and 3.0 μM thiamine, adjusted to pH 5.5. Cells were then resuspended in 1 mL of MM and adjusted to a final concentration of 1 × 10^6^ cells/mL. Cell density was determined either by direct counting using a hemocytometer (after 1:100 dilution). One milliliter of the adjusted cell suspension was transferred to Eppendorf tubes, which were placed upright in a shaking incubator and cultured at 30°C for 3 days. Tubes were oriented vertically to ensure proper aeration, and positioning was monitored daily. After incubation, cells were gently resuspended in 50-100 μL of the remaining medium and examined by DIC microscopy. The assay was performed with WT (H99), *mln1*Δ (YSB1172), *bzp4*Δ (YSB1894), the *mln1*Δ *bzp4*Δ double mutant (YSB7525), the complemented strain *mln1*Δ+*MLN1* (YSB11310), and *cac1*Δ (YSB42) and *pdr802*Δ (YSB2387). Cell body diameters were measured from DIC images using NIS-Elements v4.5, excluding the capsule. Cells with a cell body diameter of 10 μm or greater were scored as titan cells. At least 100 cells were measured per strain in each of three independent biological replicates.

### In vivo assessment of virulence and fungal burden in a murine model

To evaluate the virulence of *C. neoformans* strains in vivo, a murine inhalation model was employed. Wild-type (WT), *mln1*Δ, and *mln1*Δ+*MLN1* strains were cultured overnight in liquid YPD medium at 30°C, harvested, and washed thoroughly with PBS. Cell suspensions were adjusted to a concentration of 2.5 × 10^7^ cells/mL. Six-week-old female C57BL/6 mice (obtained from JA BIO, Suwon, Republic of Korea) were acclimatized for one week before infection. Mice were anesthetized with intraperitoneally administered Avertin (2,2,2-tribromoethanol), and 20 μL of yeast suspension (equivalent to 5 × 10^5^ cells) was administered intranasally to each mouse. For each strain, seven mice were used for survival analysis and monitored daily until predefined humane endpoints were reached. Mice were euthanized when body weight loss reached ≥ 20% of the pre-infection body weight or when severe clinical signs meeting the humane-endpoint criteria specified in the approved IACUC protocol were observed. Mice reaching these criteria were considered to have reached the survival endpoint. Survival data were analyzed using the Kaplan-Meier method and statistically evaluated with the log-rank (Mantel-Cox) test using GraphPad Prism software (version 10.4.0). For fungal burden analysis, five additional mice per strain were euthanized on day 16 post-infection. The lungs and brain were aseptically harvested, homogenized in 5 mL PBS, and plated on YPD agar containing 100 μg/mL chloramphenicol. Plates were incubated at 30°C for 3 days, after which colony-forming units (CFU) were counted and reported as CFU per gram of tissue.

### In vivo assessment of virulence in a *Drosophila melanogaster* model

The *Drosophila melanogaster w*^1118^ strain was used for the infection assays. Flies were reared at 25°C under a 12 h light/12 h dark cycle on standard cornmeal-agar medium obtained from the Bloomington Stock Center. WT and mutant *C. neoformans* strains were cultured overnight in YPD medium at 30°C, washed three times with PBS, and adjusted to a final concentration of 4 × 10^8^ cells/mL. Following centrifugation, the cell pellets were resuspended in PBS containing 1% (w/v) FD&C Blue No. 1 to facilitate visualization during injection. For the survival assay, 4-5-day-old adult female flies were anesthetized with CO_2_ and injected in the dorsal thorax with 0.1 μL of the fungal suspension, corresponding to approximately 4 × 10^4^ cells per fly, using a 33-gauge, 10 mm HPLC needle. Successful injection was confirmed by visible blue coloration of the thorax/body, whereas flies showing blue dye extending into the head were excluded from analysis. A total of 91, 89, and 95 flies were used for the WT, *mln1*Δ, and *mln1*Δ+*MLN1* strains, respectively. Flies were divided into groups of 10, and transferred to individual vials containing standard cornmeal-agar medium. Infected flies were maintained at 30°C, and survival was recorded daily for up to 4 days post-infection.

### Statistical analysis

All statistical analyses were performed using GraphPad Prism version 10.4.0. Comparisons between two groups were performed using unpaired two-tailed Student’s t-tests. Unpaired one-tailed Student’s t-tests were used in Fig. 1*B*, Supplementary Information, Fig. S9D and F, where the analysis tested a prespecified directional hypothesis of increased expression under the test condition relative to the control condition. Multiple-group comparisons were analyzed by ordinary one-way ANOVA followed by Dunnett’s or Tukey’s multiple-comparisons test, as indicated in the corresponding figure legends. Experiments involving two or three factors were analyzed by two-way or three-way ANOVA, respectively, with repeated-measures analysis used where appropriate, followed by Dunnett’s, Šídák’s, or false discovery rate (FDR) multiple-comparisons procedures as indicated in the corresponding figure legends. For single-cell measurements, individual cells were treated as nested observations within independent biological replicates and analyzed using nested one-way ANOVA, with biological replicates serving as the experimental units, as indicated in the corresponding figure legends. For quantitative growth assays, AUC was calculated for each biological replicate and compared between strains using unpaired two-tailed Student’s *t*-tests. Survival curves were compared using the log-rank (Mantel-Cox) test. *P* values < 0.05 or FDR-adjusted *q* values < 0.05 were considered statistically significant. Unless otherwise indicated, experiments were performed with at least three independent biological replicates.

## Data availability

Data supporting the findings are available in the manuscript, Supporting information files, and repositories listed below. Original uncropped images underlying the blot and gel results, together with other supporting raw data files, are available in Figshare (https://doi.org/10.6084/m9.figshare.33963328). The RNA-seq data were deposited in the Gene Expression Omnibus (GEO) database under accession number GSE343021.

## Acknowledgments

We thank Dr. Kwang-Woo Jung (Korea Atomic Energy Research Institute) and Prof. Kyung-Tae Lee (Jeonbuk National University) for generating the previously published NanoString-based transcriptional profiling and signature-tagged mutagenesis datasets that were reanalyzed to identify transcription factors examined in this study. This work was supported by grants from the National Research Foundation of Korea, funded by the Ministry of Science and ICT, Republic of Korea (RS-2025-18362970, RS-2025-02215093, and RS-2025-00555365 to Y.-S.B.); the Korea Health Technology Research and Development Project through the Korea Health Industry Development Institute, funded by the Ministry of Health and Welfare, Republic of Korea (RS-2026-25518347 to Y.-S.B.); and the Korea Institute for Advancement of Technology, funded by the Ministry of Trade, Industry and Energy, Republic of Korea (RS-2024-00418203 to Y.-S.B.). The funders had no role in study design, data collection and analysis, decision to publish, or preparation of the manuscript. During manuscript preparation, the authors used Claude (Anthropic; Opus 5) and ChatGPT (OpenAI; GPT-6 Astra) for grammar checking and language editing. All suggested revisions were reviewed and verified by the authors, who take full responsibility for the accuracy and integrity of the manuscript.

## Author Contributions

S.-R.Y., M.L., and Y.-S.B. designed research; S.-R.Y., M.L., S.-Y.Y., H.C., D.W., D.K., J.-T.C., S.Y., R.S., K.-A.L., J.-S.L., W.-J.L., and D.-W.L. performed research; J.-T.C. contributed new reagents; S.-R.Y. analyzed data; Y.-S.B. supervised research and acquired funding; S.-R.Y. and M.L. wrote the original draft; and all authors reviewed and edited the manuscript.

## Supporting information

### Supporting Materials and Methods

#### Gene expression analysis by qRT-PCR

To assess the expression of 24 pathogenicity-associated TFs, we performed quantitative reverse transcription PCR (qRT-PCR) using *C. neoformans* H99 cells grown under HMC. The H99 strain was first cultured in 50 mL of YPD broth at 30°C for 16 h with agitation. Cells were then sub-cultured into fresh YPD and incubated until the OD_600_ reached approximately 0.8. At this point, a 25 mL aliquot was harvested by centrifugation, lyophilized, and stored as the basal control sample. The remaining cells were washed three times with distilled water, resuspended in 25 mL RPMI medium supplemented with 10% fetal bovine serum (FBS), and incubated at 37°C for 3 h in a 5% CO_2_ shaking incubator (120 rpm). Following incubation, cells were collected by centrifugation and lyophilized overnight. Total RNA was extracted using the TRIzol-based Easy-BLUE reagent (iNtRON Biotechnology) and quantified with a NanoDrop spectrophotometer (Thermo Scientific). Complementary DNA (cDNA) was synthesized using a reverse transcription kit (Thermo Scientific), and qRT-PCR was performed with gene-specific primer pairs (Supporting Information, Table S2). The expression levels of target genes were normalized to that of *ACT1*, and fold changes were calculated relative to basal expression levels observed in the WT strain. For the subset of 12 TFs that showed upregulation under HMC, we further investigated their expression profiles under various environmental conditions to determine the specific cues responsible for induction. The H99 strain was again grown in YPD for 16 h at 30°C, sub-cultured to reach an OD_600_ of 0.8, washed three times with distilled water, and resuspended in 25 mL of the media (Supporting Information, Table S3). Each culture was incubated for 3 h under one of the following conditions: 30°C, 37°C, or 37°C with 5% CO_2_. After incubation, cells were lyophilized and processed for RNA extraction and cDNA synthesis as described above.

#### Carbon starvation- and heat stress-responsive gene expression analysis

To examine the regulation of *MLN1* during carbon starvation, WT (H99), *mig1*Δ (YSB9271), and *snf1*Δ (YSB2372) strains were cultured overnight in YPD medium at 30°C and subcultured into fresh YPD medium at an initial OD_600_ of 0.2. The *mig1*Δ mutant carries an internal deletion of the *MIG1* genomic region encoding amino acids 61-362, encompassing the C2H2-type zinc-finger DNA-binding region, which was replaced by the *NAT*-STM cassette. Correct disruption of the locus was verified by Southern blot analysis (Supporting Information, Fig. S8*F*). Cells were grown at 30°C until the OD_600_ reached approximately 0.8, and an aliquot of each culture was collected as the basal sample. The remaining cells were harvested and transferred to YNB medium without amino acids and without added glucose to induce carbon starvation. Samples were collected at 15, 30, 60, and 120 min after transfer. Total RNA extraction, cDNA synthesis, and qRT-PCR were performed as described above using the gene-specific primers listed in Supporting Information, Table S2. *MLN1* and *NTH2* transcript levels were normalized to *ACT1*, and relative expression levels were calculated using the WT basal sample as the reference.

To examine transcriptional responses during thermal adaptation, WT (H99) and *mln1*Δ (YSB1172) cells were cultured overnight in YPD medium at 30°C and subcultured into fresh YPD medium at an initial OD_600_ of 0.2. Cultures were grown at 30°C until the OD_600_ reached approximately 0.8, and an aliquot was collected as the basal sample. The remaining cultures were directly shifted to 39°C and incubated for 1, 2, and 24 h. At each time point, aliquots were collected in parallel with those used for metabolite profiling, rapidly harvested, and stored frozen until RNA extraction. Total RNA extraction, cDNA synthesis, and qRT-PCR were performed as described above. Transcript levels of *TPS1*, *TPS2*, *NTH1*, *NTH2*, and *MLN1* were measured using the primers listed in Supporting Information, Table S2 and normalized to *ACT1*. Relative expression levels were calculated using the WT basal sample as the reference.

#### Protein domain analysis and structural modeling

The Mln1 amino acid sequence (CNAG_04837) was analyzed using InterPro to identify conserved protein domains [1]. The predicted basic helix-loop-helix (bHLH) domain spanning residues 280-366 was subjected to structural modeling using the AlphaFold 3 Server with default settings [2]. For monomeric structure prediction, the Mln1 bHLH domain sequence was submitted as a single protein chain. To examine a possible dimeric arrangement, two identical copies of the same sequence were submitted as separate protein chains. Model confidence was evaluated using predicted template modeling (pTM), interface predicted template modeling (ipTM), and predicted aligned error (PAE).

#### Comparative conservation analysis of transcription factors and ortholog classification

To examine the conservation of *C. neoformans* TFs across eukaryotes, we adapted the previously published BLASTp-based conservation matrix from Jung et al. [3] for the 12 host-cue-responsive TFs. The taxonomic classification scheme was retained from the original study. For orthology-based analysis of *MLN1*, EggNOG-mapper v2.1.12 was used with DIAMOND-based search against the EggNOG v5 database [4, 5]. The DIAMOND search was performed in sensitive mode with an E-value threshold of 0.001. No additional sequence identity or query-coverage cutoff was applied. The --target_orthologs all and --report_orthologs options were used to retrieve ortholog candidates across available taxa. Species information was extracted and converted into full taxonomic lineages using TaxonKit [6]. Parsed taxonomy data were used to tabulate the number of orthologs assigned to each rank from kingdom to species. In-house bash scripts were used to automate lineage parsing and frequency counting. Protein sequences of the identified Mln1 orthologs were aligned using MUSCLE implemented in MEGA 11. The best-fit amino acid substitution model was determined by maximum-likelihood model selection in MEGA 11. A maximum-likelihood tree was generated in MEGA 11 under the Jones-Taylor-Thornton (JTT) model with gamma-distributed rate variation and invariant sites (JTT+G+I). The tree was midpoint-rooted and visualized using MEGA 11 [7]. Representative orthologs from major fungal phyla including Basidiomycota, Mucoromycota, and filamentous Ascomycota (e.g., *Aspergillus fumigatus*, *Botrytis cinerea*, *Fusarium oxysporum*) were included. Species with multiple orthologs retained all matching sequences in the tree. The tree was used to visualize the taxonomic distribution and sequence relationships of Mln1 orthologs across fungal lineages.

#### Construction of double knockout strains

The strains and primers used in this study are listed in the supporting tables (Supporting Information, Tables S1 and S2), respectively. Strains from the previously published TF mutant library were used as parental strains to construct double mutants [3]. *C. neoformans* double knockout mutants were generated in the H99 strain background by performing double joint PCR followed by biolistic transformation. The 5’ and 3’ regions of *PDR802*, *BZP4*, *ZNF2*, *FZC39*, and *FZC30* were PCR-amplified using primer pairs listed in Supporting Information, Table S2. The G418 sulfate/neomycin resistance (*NEO*) marker was used as a selection marker, amplified with M13 primer pairs from the pNEO plasmid. PCR products for the flanking regions and selection marker were purified together. The gene disruption cassette was constructed using overlap PCR. For biolistic transformation, the parental strain was cultured in YPD broth overnight at 30°C in a shaking incubator, spun down, spread on a YPD agar plate supplemented with 1 M sorbitol, and incubated at 30°C for 3 h. The purified overlap PCR product was precipitated onto 0.6 μm gold beads and delivered into the parental strain on a YPD+1 M sorbitol plate using a particle delivery system (PDS-1000, Bio-Rad). After transformation, cells were incubated for a further 4 h, recovered using a cell scraper, and transferred onto a YPD agar plate containing 50 μg/mL G418 (YPD+G418). For mutant screening, G418-resistant positive transformants were picked, re-streaked on YPD+G418 plates and used for genomic DNA preparation. Positive transformants were first screened by diagnostic PCR using screening primer pairs listed in Supporting Information, Table S2 and subsequently verified by Southern blot analysis (Supporting Information, Fig. S4). The original uncropped blot and gel images are deposited in Figshare as described in the Data availability section. To confirm the reproducibility of the observed phenotypes and reduce the likelihood of effects from unintended mutations, at least two independently generated deletion mutants were constructed for each targeted gene.

#### Genotyping by Southern blot analysis

To confirm successful gene disruption in the selected mutants, both diagnostic PCR and Southern blot analyses were performed. PCR-positive transformants were initially cultured in 50 mL of YPD broth at 30°C for 16 h with shaking. The cells were collected by centrifugation and lyophilized overnight. Genomic DNA was isolated using the cetyl trimethylammonium bromide (CTAB) extraction method and dissolved in TE buffer. For Southern blot analysis, genomic DNA from WT and mutant strains was digested with the restriction enzymes indicated in the corresponding figures and separated on a 1% agarose gel by electrophoresis. The gel was incubated in denaturation buffer (1.5 M NaCl, 0.5 M NaOH) for 45 min, followed by neutralization in buffer containing 1.5 M NaCl and 0.5 M Tris-HCl (pH 8.0) for an additional 45 min. DNA fragments were transferred to a nitrocellulose membrane using an upward capillary blotting method over 16 h. The membrane was subsequently crosslinked by ultraviolet irradiation at a dose of 120 mJ/cm^2^. A gene-specific hybridization probe was generated by PCR using PO primers (listed in Supporting Information, Table S2), and radiolabeled with [α-^32^P]dCTP. Hybridization was carried out overnight at 65°C in Church buffer (1 mM EDTA, 0.25 M Na_2_HPO_4_, 0.17% H_3_PO_4_, 1% hydrolyzed casein, 7% SDS). After hybridization, the membrane was washed with SSC buffer to remove nonspecific binding. Radioactive signals were visualized using the Amersham Typhoon Biomolecular Imager (Cytiva, UK), enabling confirmation of gene replacement events in the transformants.

#### Chemical susceptibility assays and amino acid mixture composition

Each mutant strain and the WT control were cultured overnight at 30°C in YPD medium and diluted to an initial OD_600_ of 0.2. To examine growth under chemical stress, 10-fold serial dilutions were spotted onto YPD plates containing fludioxonil (FDX), fluconazole (FLC), CdSO_4_, amphotericin B (AMB), or diamide (DIA) at the concentrations indicated in the figures, and plates were incubated at 30°C for up to 3 days with daily imaging. For nitrogen source utilization assays, the 1× amino acid mixture was prepared as previously described [8] and contained 30 mg/L L-isoleucine, 150 mg/L L-valine, 20 mg/L adenine sulfate, 20 mg/L L-histidine-HCl, 100 mg/L L-leucine, 30 mg/L L-lysine, 20 mg/L L-methionine, 50 mg/L L-phenylalanine, 20 mg/L L-tryptophan, 30 mg/L L-tyrosine, 20 mg/L uracil, 400 mg/L L-serine, 100 mg/L L-glutamate, 200 mg/L L-threonine, and 100 mg/L L-aspartate.

#### Acute heat-shock assay

WT, *mln1*Δ, and *mln1*Δ+*MLN1* strains were cultured overnight in YPD medium at 30°C and subcultured into fresh YPD medium at an initial OD_600_ of 0.2. Cells were grown at 30°C until the OD_600_ reached approximately 0.8. Cells corresponding to a total OD_600_ of 5.0 were harvested and resuspended in 500 μL of sterile distilled water. Cell suspensions were serially diluted, and 3 μL of each dilution was spotted onto YPD agar plates. Plates representing the 0 min control were transferred directly to 30°C, whereas the remaining plates were placed in an incubator preheated to 55°C for 15, 20, 25, or 30 min to impose acute heat shock. Following heat treatment, the plates were immediately transferred to 30°C and incubated for 3 days before imaging.

#### Melanin production assay

To evaluate melanin production, overnight cultures were harvested, washed twice with phosphate-buffered saline (PBS), and spotted onto L-3,4-dihydroxyphenylalanine (L-DOPA) agar plates. The L-DOPA medium comprised 1 g/L L-asparagine, 3 g/L KH_2_PO_4_, 250 mg/L MgSO_4_, 1 mg/L thiamine, 5 μg/L biotin, 100 mg/L L-DOPA, and 0.1% glucose. L-DOPA plates were incubated at 37°C and imaged after 2 days.

#### Capsule production assay

Capsule formation was induced under minimal-medium conditions. Cells were grown overnight in YPD at 30°C, harvested, washed twice with PBS, and resuspended in minimal medium containing 10.0 mM MgSO_4_, 29.4 mM KH_2_PO_4_, 13.0 mM glycine, and 3.0 μM thiamine, adjusted to pH 5.5, and supplemented with the indicated carbon source at 7.5 mM [9]. Cultures were incubated at 37°C for 3 days with shaking at 200 rpm. Capsules were visualized by counterstaining with India ink and imaged by differential interference contrast microscopy (DIC) (Nikon Eclipse) equipped with a DS-Qi2 digital camera. Capsule thickness was calculated as (total cell diameter - cell body diameter) / 2, measured using NIS-Elements v4.5.

#### Trehalase gene expression under glucose- and trehalose-containing conditions

WT and *mln1*Δ cells were cultured overnight in YPD medium at 30°C and subcultured into fresh YPD medium at an initial OD_600_ of 0.2. Cells were grown to an OD_600_ of approximately 0.8, harvested, washed, and transferred to YP medium supplemented with either 2% glucose or 2% trehalose prepared in liquid or agar format. Cells were incubated at 30°C for 24 h. Liquid cultures were harvested by centrifugation, whereas cells grown on agar were collected by scraping the plate surface. Total RNA extraction, cDNA synthesis, and qRT-PCR were performed as described above. *NTH1* and *NTH2* transcript levels were normalized to *ACT1* and expressed relative to WT cells grown in liquid YPD medium.

#### Construction of *MLN1*-mRuby3 and complemented strains

To visualize Mln1 localization, a construct expressing an Mln1-mRuby3 fusion protein under the control of the native *MLN1* promoter was generated. A 4,906 bp fragment spanning the *MLN1* promoter and the complete coding region, excluding the stop codon, was amplified from H99 genomic DNA using the primers listed in Supporting Information, Table S2 and assembled into the NotI-linearized p*NEO*-mRuby3 vector using the NEBuilder HiFi DNA Assembly Cloning Kit (#E5520, New England Biolabs) to yield the 11,841 bp plasmid p*NEO*-*MLN1*-mRuby3. The assembled plasmid was verified by restriction digestion and by Sanger sequencing across the assembly junctions and the entire *MLN1* coding region. The plasmid was linearized with StuI within the *MLN1* promoter region and introduced into the *mln1*Δ mutant (YSB1172) by biolistic transformation as described above. Transformants were selected on YPD agar containing 50 μg/mL G418, and integration at the *MLN1* locus was confirmed by diagnostic PCR using the genomic primer CSO (B11420), located outside the transformed construct, together with the construct-internal primer PO (B1764), yielding the expected 2,298 bp product. A single verified transformant (YSB10818) was used for all localization experiments.

For complementation, the *MLN1* gene, including its native promoter and terminator regions, was amplified from H99 genomic DNA and fused to the *NEO* marker by Gibson assembly using the primers listed in Supporting Information, Table S2. The resulting cassette was introduced ectopically into the *mln1*Δ mutant (YSB1172) by biolistic transformation, and transformants were selected on YPD agar containing 50 μg/mL G418. The presence of the introduced *MLN1*-*NEO* cassette was confirmed by diagnostic PCR, and one verified transformant (YSB11310) was designated *mln1*Δ+*MLN1* and used for all subsequent analyses.

#### RNA-seq processing and computational parameters

Raw sequencing reads were subjected to quality control and adapter trimming using Cutadapt v2.4 in Python 3.7.4, employing the specified adapter sequence [10]. Subsequent read processing followed previously established protocols [11]. Cleaned reads were aligned to the *C. neoformans* H99 reference genome using HISAT2 v2.2.1, a splice-aware aligner. Genome annotation files were retrieved from the NCBI FTP repository. HISAT2 was run with -p 30 and --dta, with paired-end reads supplied using -1 and -2. The resulting alignment files were converted and sorted using SAMtools v0.1.19, applying “-Sb -@ 8” for file conversion and “-@ 20 -m 2000000000” for sorting operations [12]. Gene-level expression matrices were generated and further analyzed using the R package IsoformSwitchAnalyzeR [13]. Data quality was evaluated with DEBrowser [14], and differential expression results were visualized using the EnhancedVolcano package in R v4.1.0 [15]. For functional enrichment analysis, gene lists were submitted to DAVID as CNAG locus tags, with *C. neoformans* var. grubii H99 selected as the species. The annotation categories GOTERM_BP_DIRECT, GOTERM_MF_DIRECT, GOTERM_CC_DIRECT, and KEGG_PATHWAY were selected. Enrichment results were visualized as dot plots in which dot size indicates the number of genes assigned to each term and color intensity indicates −log_10_ *P* value. Heatmaps were generated using Morpheus (Broad Institute) to visualize relative expression patterns across samples.

#### Metabolite extraction, derivatization, and untargeted GC-MS analysis

For metabolite extraction, each collected cell pellet was resuspended in 1 mL of 75% ethanol preheated to 80°C and transferred to a screw-cap tube containing zirconia beads. Samples were incubated in an 80°C water bath for 3 min and immediately cooled on ice for 3 min. Cells were then disrupted by bead beating for 30 s, followed by cooling on ice for 3 min. This bead-beating and cooling cycle was repeated twice. The resulting extracts were centrifuged at 13,000 rpm for 5 min at 4°C, and 700 μL of the cleared extract was transferred to a 15-mL conical tube. Samples were stored at −80°C, lyophilized, and subsequently subjected to mass spectrometry-based metabolite profiling. Lyophilized metabolite extracts were reconstituted in 1 mL of 80% methanol. After addition of 10 mM heptadecanoic acid as an internal standard, the samples were thoroughly vortexed and centrifuged at 13,000 rpm for 10 min to remove insoluble materials. A 160-μL aliquot of each supernatant was vacuum-dried at 40°C and subjected to a two-step derivatization process. First, the dried samples were treated with 60 μL of methoxyamine hydrochloride in pyridine (MOX) and incubated for 2 h (1500 rpm, 37°C) using a micromixer to minimize formation of multiple derivatives. Subsequently, the samples were derivatized with 100 μL N-methyl-N-(trimethylsilyl)trifluoroacetamide containing 1% trimethylchlorosilane (MSTFA + 1% TMCS) for 20 min (200 rpm, 70°C). The derivatized samples were analyzed by GC-MS as previously described [16]. Briefly, the autosampler injected 1 μL of each sample in splitless mode into an Agilent 7890A gas chromatograph equipped with an HP-5MS capillary column (30 m × 0.250 mm i.d.; 0.25-μm film thickness; Agilent J&W Scientific). The flow rate of helium gas was 1.1 mL/min. The oven temperature was held at 75°C for 4 min, then increased to 280°C at a rate of 4°C/min, and held for 1.56 min. The mass spectrum was recorded over an m/z range of 40–600 with a scan time of 0.2 s. Raw data were processed through peak detection, quantification, alignment, and annotation using MZmine v4.10.6 software [17]. Metabolite abundances were normalized to the internal standard, and features with more than 50% missing values among samples were excluded. The remaining missing values were imputed with 1/5 of the minimally detected peak area, followed by log_10_ transformation and Pareto scaling prior to downstream analysis. Metabolomic data preprocessing and scaling were performed using MetaboAnalyst v6.0 [18]

#### Trehalose assay reaction conditions and calculations

Cells corresponding to 5 OD_600_ units were harvested and washed twice with ice-cold distilled water. Intracellular metabolites were extracted by resuspending the cell pellet in 500 μL of boiling distilled water and heating at 100°C for 15 min with intermittent vortexing. Cell debris was removed by centrifugation, and the resulting supernatant was used for trehalose measurement.

For each reaction, 20 μL of sample extract was mixed with 200 μL of distilled water, 20 μL of Bottle 1 reagent, 10 μL of reconstituted Bottle 2 reagent, and 2 μL of Bottle 3 reagent. After incubation for 5 min at approximately 25°C, absorbance at 340 nm was measured (A1). Trehalase (2 μL; Bottle 4) was then added, and samples were incubated for 10 min before a second measurement at 340 nm (A2). Blank reactions and the supplied glucose standard (0.4 mg/mL) were analyzed in parallel. The trehalase-dependent absorbance change was calculated as ΔA = A2 − A1 and corrected using the corresponding blank. Trehalose concentration was calculated relative to the glucose standard as follows: Trehalose (mg/mL) = (ΔA sample / ΔA glucose standard) × 0.4 × 0.950. The factor 0.950 represents the molecular-weight conversion factor for converting the glucose equivalent to trehalose [342.30/(2 × 180.16)]. Trehalose amounts were calculated from the extract volume and normalized to the number of cells analyzed.

### Supplementary figures and figure legends

**Figure S1.**
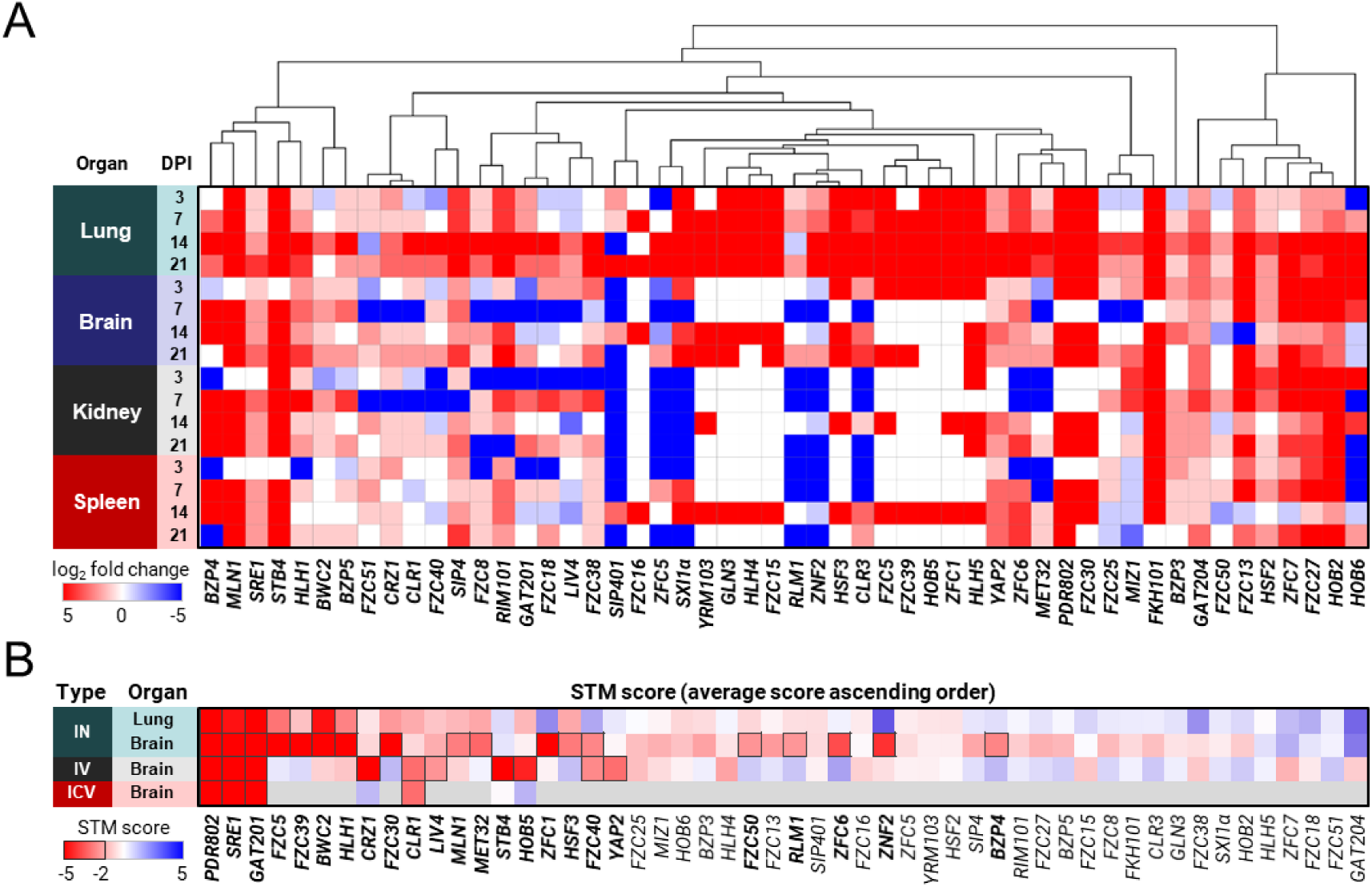
Evaluation of NanoString and STM data highlights TFs critical for in vivo adaptation. (A) In vivo expression profiling was performed using NanoString technology. A heatmap shows 52 TFs upregulated by at least 2-fold in both the lungs and brain during infection, relative to in vitro levels. (B) Signature-tagged mutagenesis (STM) analysis was conducted using three infection routes: intranasal (IN), intravenous (IV), and intracerebroventricular (ICV). TF mutants exhibiting reduced infectivity (STM score ≤ −2) in at least one infection model were selected for further analysis. A total of 24 TF mutants met this criterion and are indicated by black outlines.

**Figure S2.**
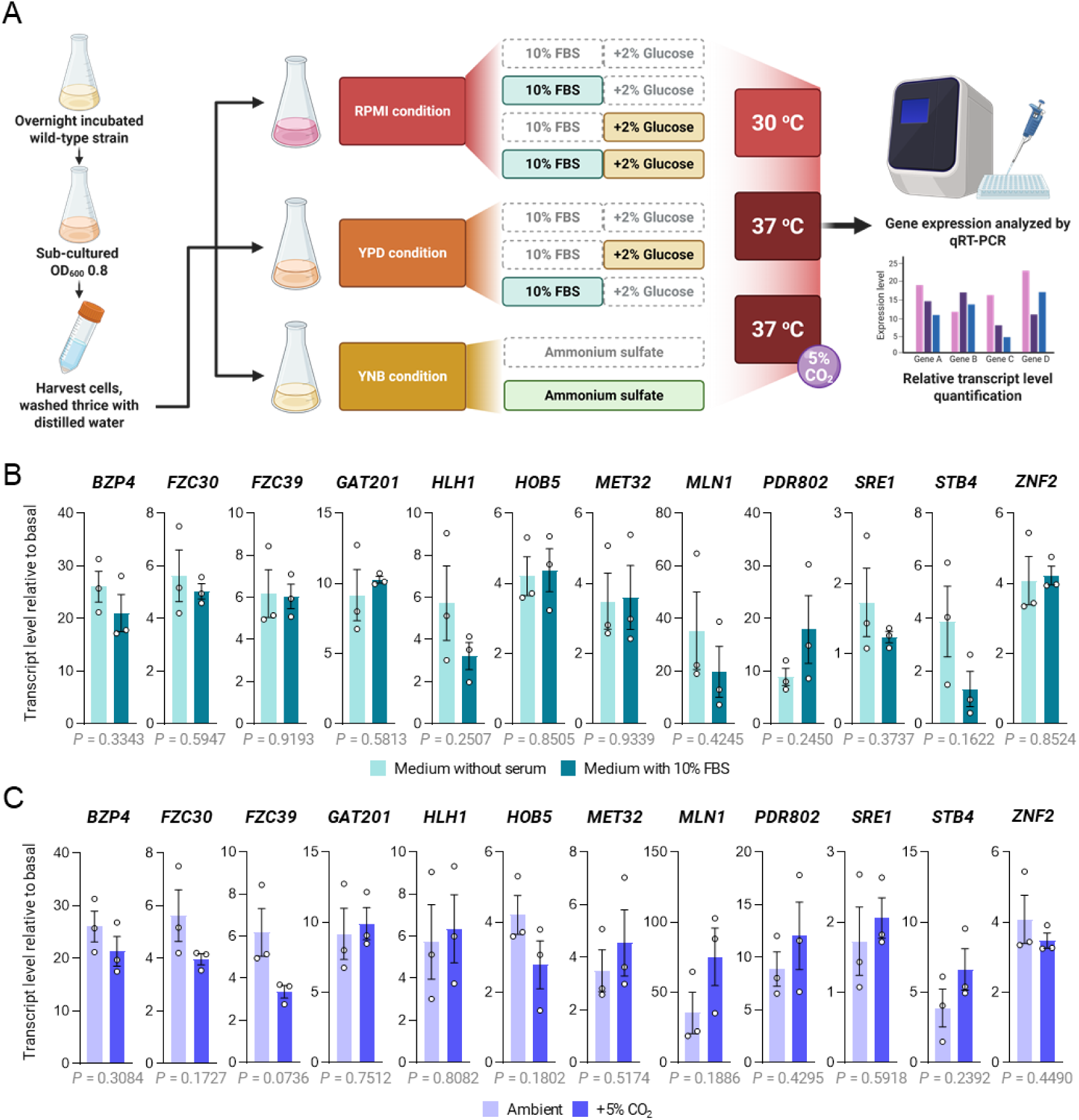
Expression analysis of 12 in vivo-regulated pathogenicity-related transcription factors under additional host-mimicking conditions. (A) Schematic overview of the experimental design used to evaluate transcription factor (TF) expression under host-mimicking conditions. Wild-type *C. neoformans* cells were grown overnight, subcultured to OD_600_ 0.8, washed, and transferred to RPMI, YPD, or YNB-based media designed to assess responses to host-relevant cues, including elevated temperature, serum supplementation, high CO_2_, glucose availability, and nitrogen availability. The schematic diagram was created with BioRender (https://BioRender.com/45upvnl). (B-C) Expression levels of selected pathogenicity-related transcription factors were quantified by quantitative RT-PCR using cDNA synthesized from cells grown under indicated conditions. Gene-specific primers listed in Supporting Information, Table S2 were used. All experiments were conducted with three independent biological replicates, each including three technical replicates. Fold changes were calculated relative to basal expression levels in RPMI medium at 30°C. (B) Comparison of gene expression at 37°C in RPMI medium with or without supplementation of 10% fetal bovine serum (FBS). (C) Comparison of gene expression at 37°C in RPMI medium under ambient air or 5% CO_2_ atmosphere.

**Figure S3.**
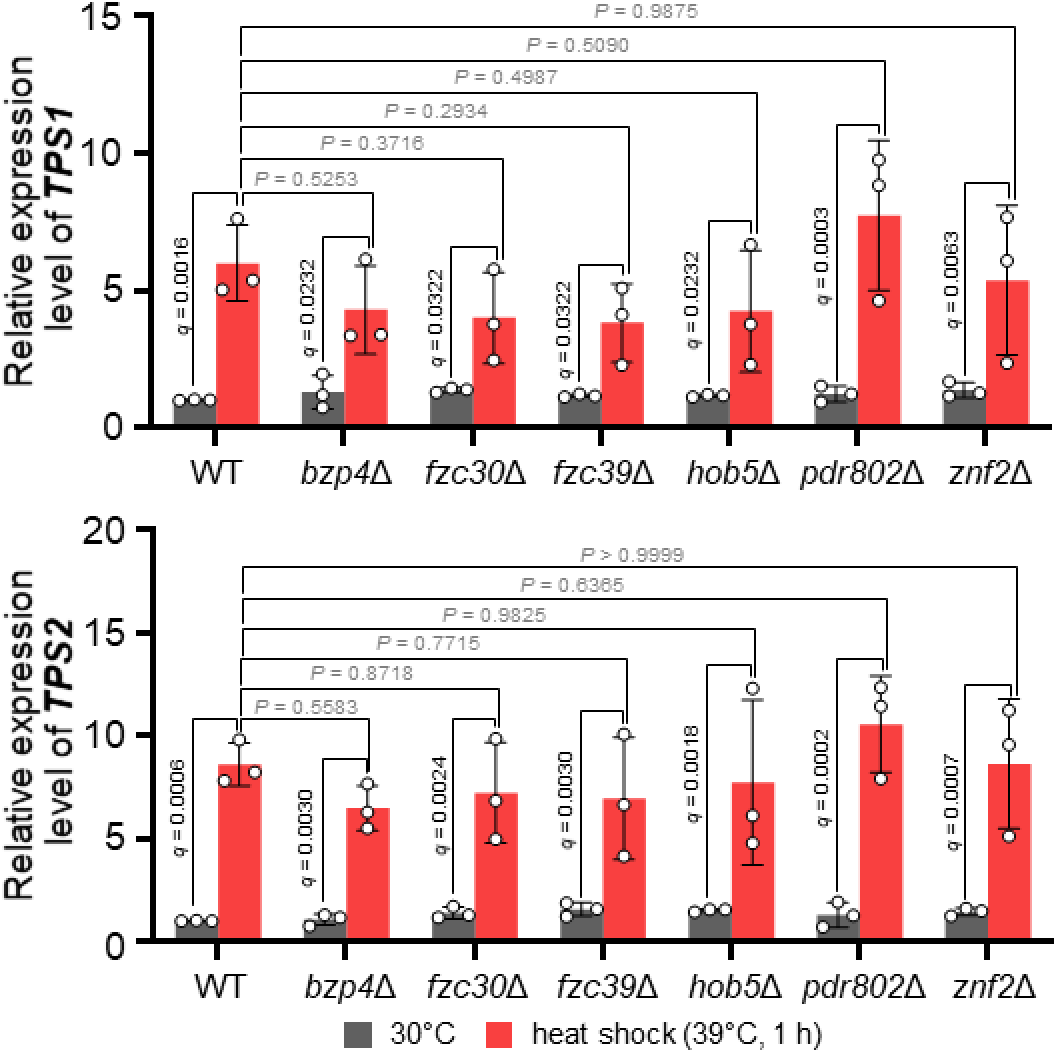
Temperature-induced expression of trehalose biosynthetic genes is maintained in temperature-responsive TF deletion mutants. Relative expression levels of *TPS1* and *TPS2* were determined in the WT and the indicated TF deletion mutants under basal conditions and following temperature upshift to 39°C for 1 h. Data represent the mean ± SD of three biological replicates. Statistical significance was assessed by two-way repeated-measures ANOVA. Comparisons between basal and 39°C conditions within each strain were performed using the two-stage linear step-up procedure of Benjamini, Krieger, and Yekutieli with a false discovery rate (FDR) of 5% (*Q* = 0.05). Comparisons between the WT and each mutant within the same condition were performed using Dunnett’s multiple comparisons test. Exact *q* values are shown for basal versus 39°C comparisons, whereas exact *P* values are shown for comparisons with the WT. Statistically significant *q* or *P* values are shown in black, whereas nonsignificant values are shown in gray.

**Figure S4.**
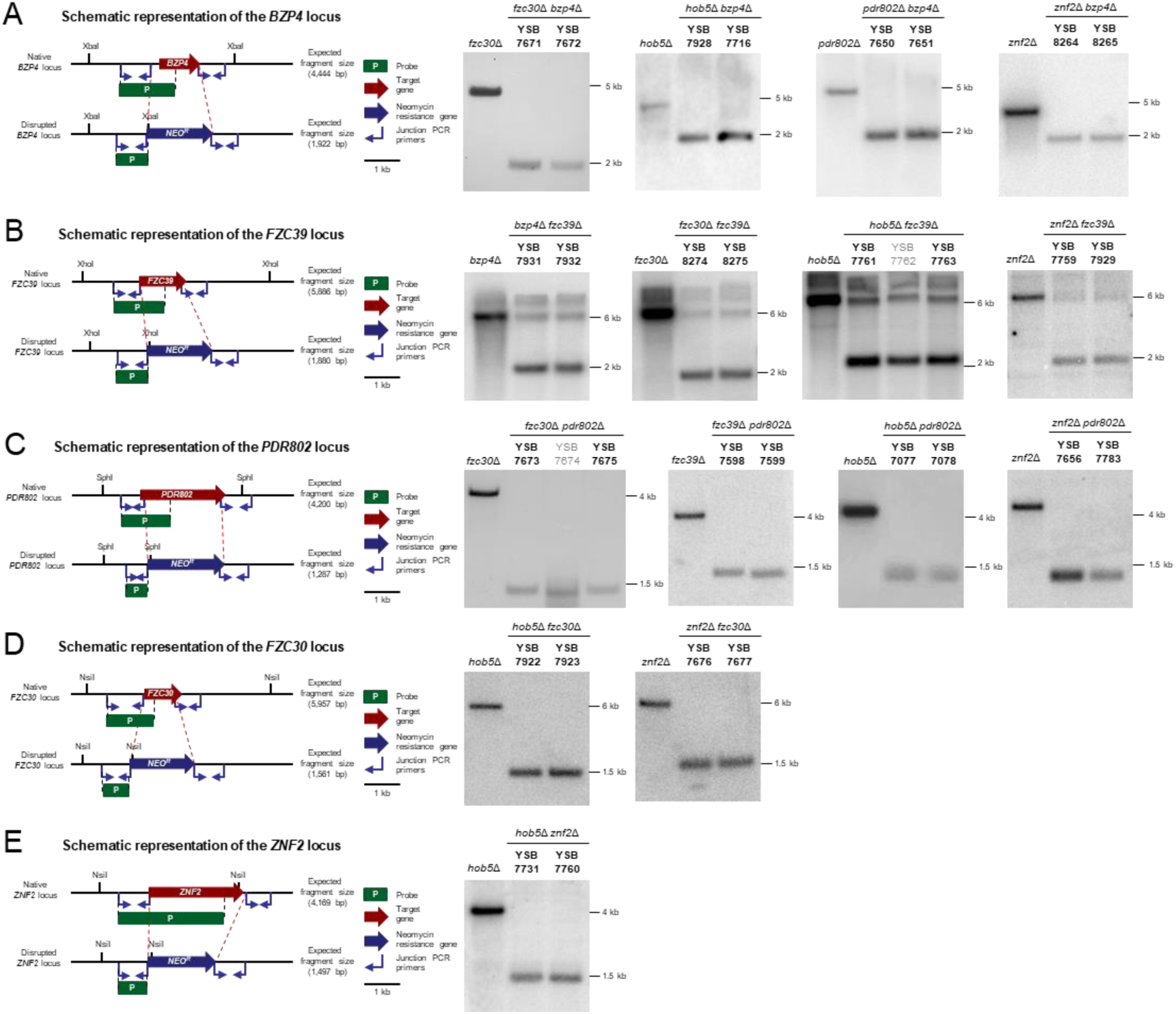
Construction and validation of pairwise double-knockout mutants among six host-temperature-responsive transcription factors. All 15 pairwise double-knockout combinations were generated from six transcription factor genes, *BZP4*, *FZC39*, *PDR802*, *FZC30*, *ZNF2*, and *HOB5*. Schematic diagrams show the native loci and the corresponding disrupted alleles, in which each target gene was replaced with the neomycin-resistance marker (*NEO*) by homologous recombination. Red dashed lines indicate the homologous recombination regions. Green bars indicate the probes used for Southern blot analysis, and blue arrows indicate the junction PCR primers. Representative Southern blot results confirming the expected genomic replacement patterns are shown for each double-knockout strain. Each blot includes the parental or control strain and independently generated mutant strains, as indicated above the lanes. Gray labels indicate transformants that failed Southern blot validation and were excluded from subsequent analyses.

**Figure S5.**
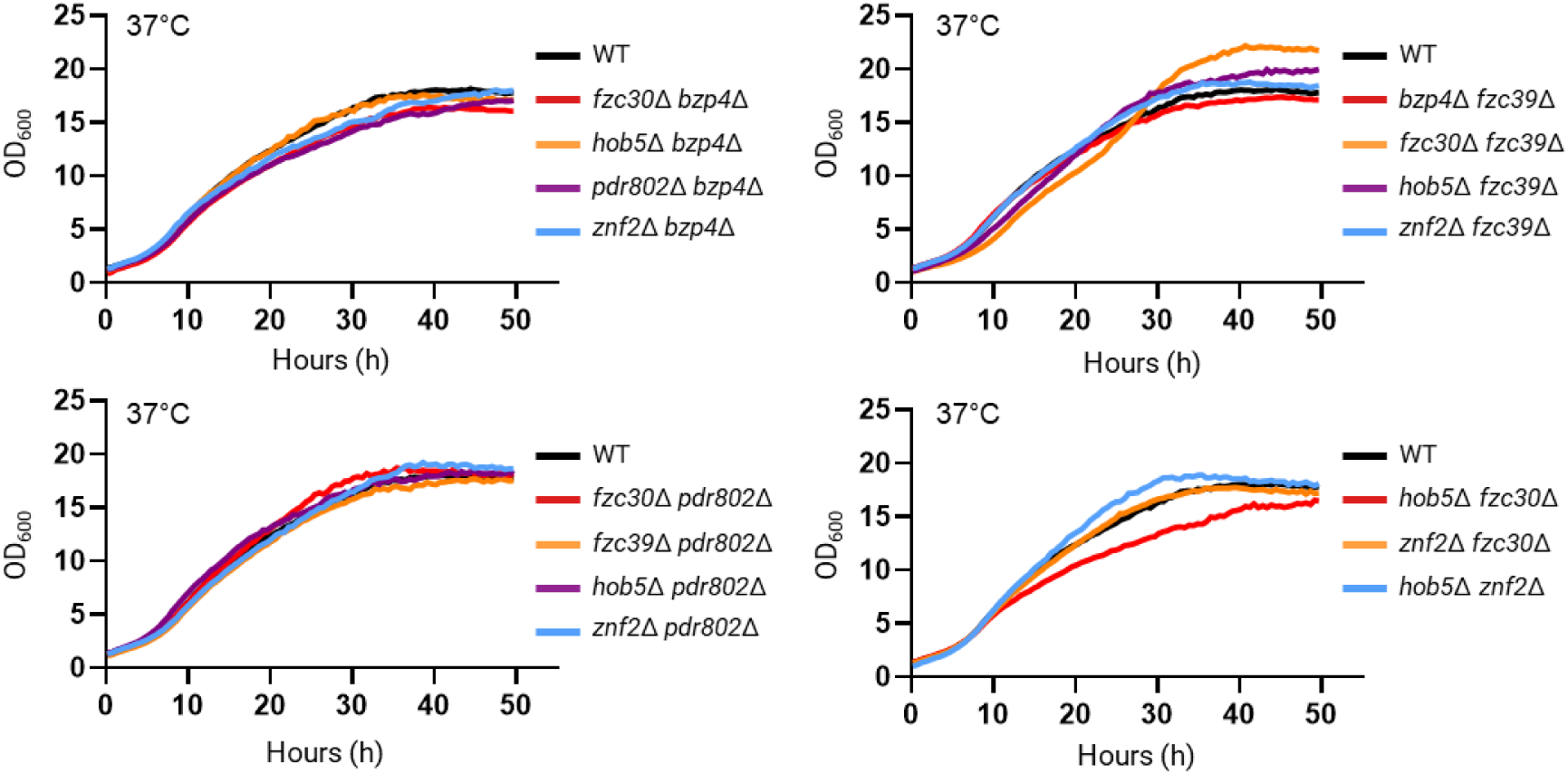
Growth at 37°C is maintained despite pairwise deletion of temperature-responsive transcription factors. Representative isolates of all 15 pairwise double-deletion combinations of *BZP4*, *FZC30*, *FZC39*, *HOB5*, *PDR802*, and *ZNF2* were cultured together with WT at 37°C, and growth was continuously monitored by measuring OD_600_ in a separate 16-channel bioreactor experiment for each temperature. The strains were YSB7671 (*fzc30*Δ *bzp4*Δ), YSB7928 (*hob5*Δ *bzp4*Δ), YSB7650 (*pdr802*Δ *bzp4*Δ), YSB8264 (*znf2*Δ *bzp4*Δ), YSB7931 (*bzp4*Δ *fzc39*Δ), YSB8274 (*fzc30*Δ *fzc39*Δ), YSB7761 (*hob5*Δ *fzc39*Δ), YSB7759 (*znf2*Δ *fzc39*Δ), YSB7598 (*fzc39*Δ *pdr802*Δ), YSB7673 (*fzc30*Δ *pdr802*Δ), YSB7077 (*hob5*Δ *pdr802*Δ), YSB7656 (*znf2*Δ *pdr802*Δ), YSB7922 (*hob5*Δ *fzc30*Δ), YSB7676 (*znf2*Δ *fzc30*Δ), and YSB7731 (*hob5*Δ *znf2*Δ). For clarity, the 15 double-mutant growth curves were divided among four panels at each temperature to avoid excessive overlap. Because all strains were analyzed simultaneously within each experiment, the identical WT growth curve is reproduced in each panel as a common reference.

**Figure S6.**
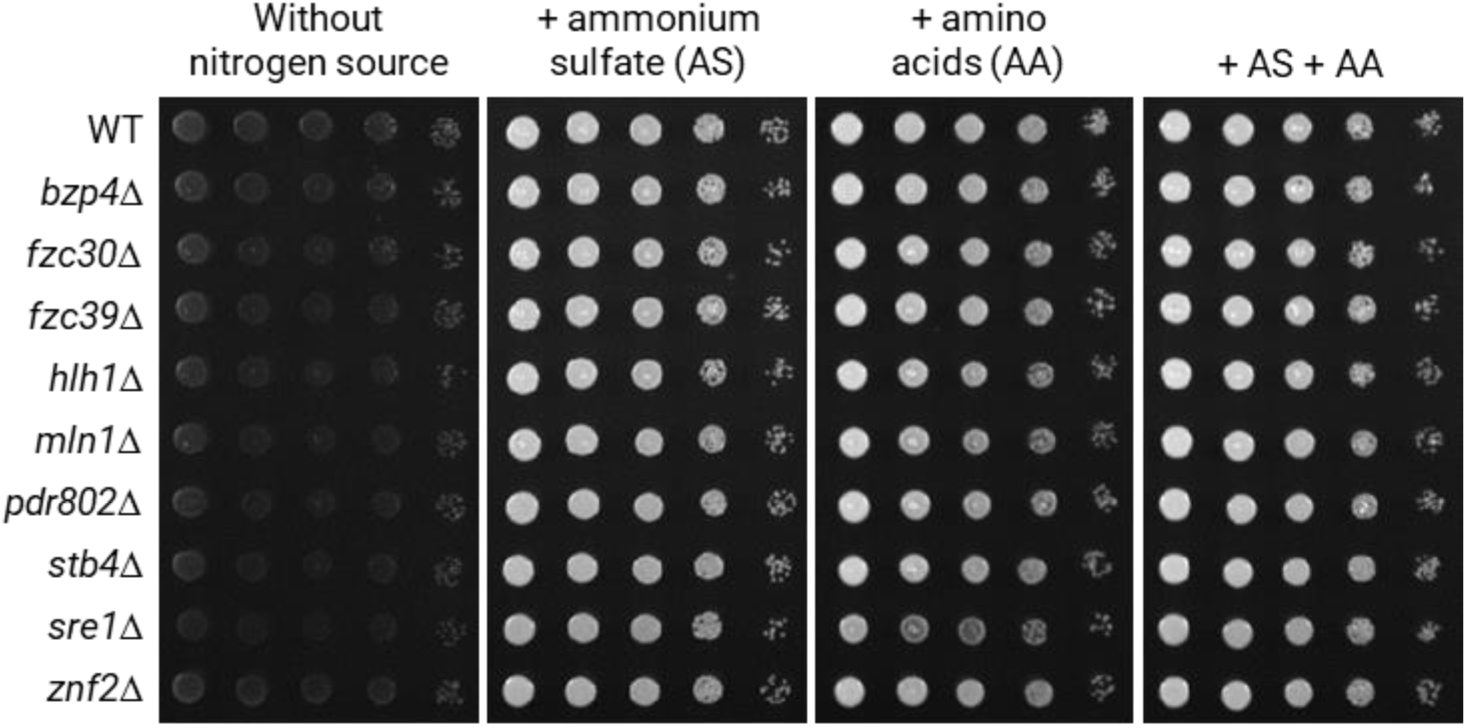
Nitrogen source utilization of nitrogen-responsive transcription factor mutants. Growth of WT (H99) and the nine nitrogen-responsive transcription factor deletion mutants, *bzp4*Δ (YSB1894), *fzc30*Δ (YSB2447), *fzc39*Δ (YSB1820), *hlh1*Δ (YSB1175), *mln1*Δ (YSB1172), *pdr802*Δ (YSB2387), *stb4*Δ (YSB1013), *sre1*Δ (YSB2494), and *znf2*Δ (YSB2740), was assessed on solid media containing different nitrogen sources. Cells were grown overnight in YPD at 30°C, washed twice with distilled water, and resuspended in sterile water. Ten-fold serial dilutions were spotted onto YNB agar containing 2% glucose but lacking amino acids and ammonium sulfate, or onto the same medium supplemented with ammonium sulfate (AS), an amino acid mixture (AA), or both (AS + AA). Plates were incubated at 30°C and imaged after 2 days. Images are representative of three independent biological replicates.

**Figure S7.**
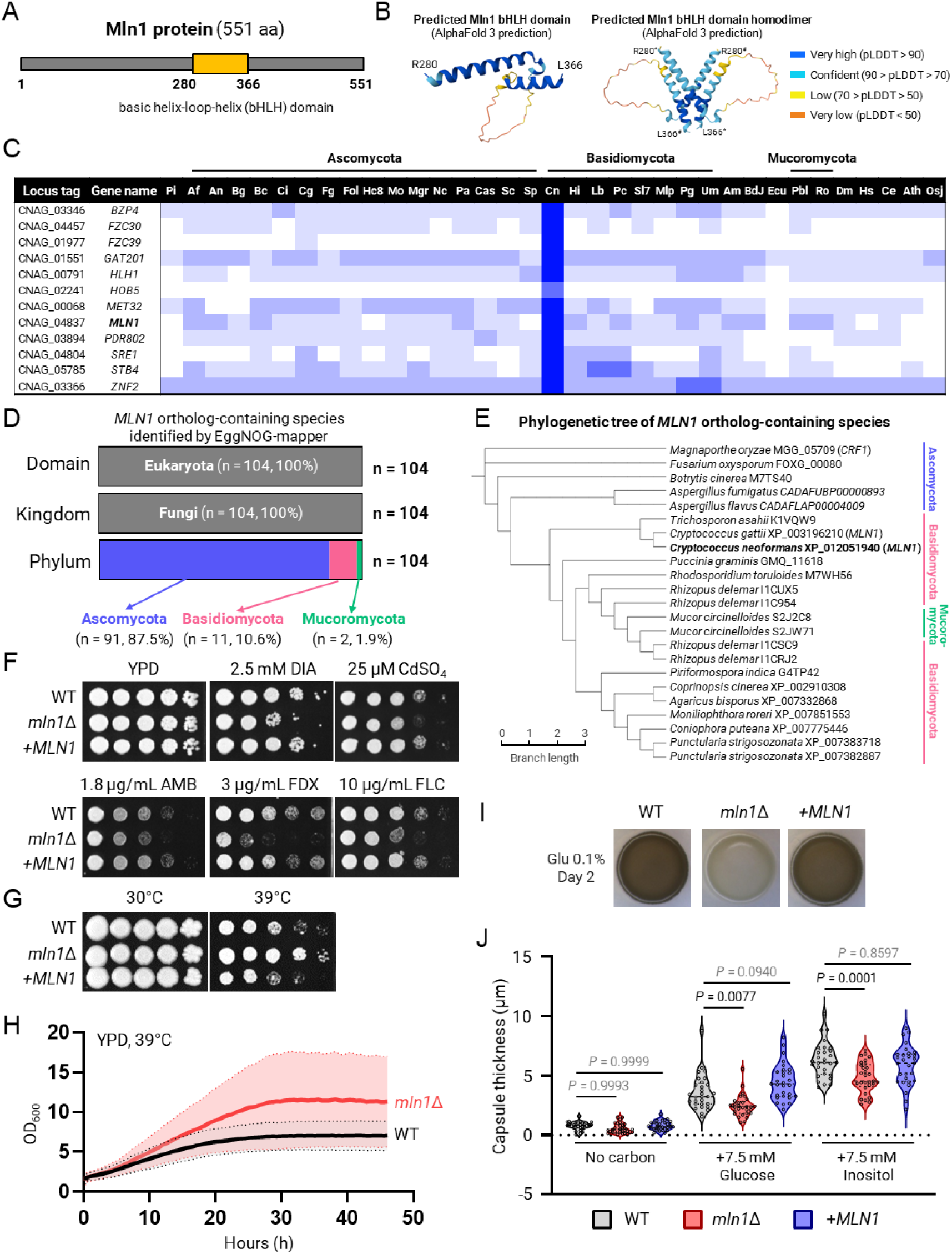
Conservation and phenotypic characterization of Mln1 in *C. neoformans*. (A) Schematic representation of the 551-amino-acid Mln1 protein showing the predicted basic helix-loop-helix (bHLH) domain spanning residues R280–L366. (B) AlphaFold 3-predicted structure of the isolated Mln1 bHLH domain and homodimeric model generated using two identical copies of the Mln1 bHLH domain sequence. The predicted homodimeric model complex showed moderate inter-chain confidence (ipTM = 0.56; pTM = 0.57). Structural models are colored according to prediction confidence (Dark blue, pLDDT >90; light blue, 70–90; yellow, 50–70; orange, <50). The two Mln1 bHLH monomers within the homodimer are distinguished by asterisks (*) and hash symbols (#), respectively. (C) Conservation of host-cue-responsive candidate transcription factors (TFs) across fungi and other eukaryotes. A BLAST-based similarity matrix was used to compare protein sequences of *C. neoformans* var. *grubii* H99 TFs with homologous sequences from representative fungal and non-fungal eukaryotic species. Sequence similarity is shown by a blue color gradient based on BLAST E-values, with darker blue indicating stronger similarity. Species abbreviations, phylogenetic color coding, and the BLAST similarity matrix were adapted from Jung et al [3]. (D) Taxonomic distribution of *MLN1* ortholog-containing species identified by EggNOG-mapper. *MLN1* orthologs were identified using a DIAMOND-based search against the EggNOG v5 database. A total of 104 species containing *MLN1* orthologs were identified and classified based on NCBI taxonomy. Ortholog-containing species were summarized at the phylum level. (E) Maximum-likelihood phylogenetic tree of representative Mln1 orthologs. Protein sequences were aligned using MUSCLE, and the phylogenetic tree was constructed in MEGA11 using the maximum-likelihood method with the best-fit amino acid substitution model. Branch lengths indicate evolutionary distance, and species are colored according to fungal phylum. (F) Spot dilution assays assessing stress resistance of WT (H99), *mln1*Δ (YSB1172), and the complemented strain *mln1*Δ+*MLN1* (YSB11310). Cells were grown overnight in YPD at 30°C, serially diluted 10-fold, and spotted onto YPD plates containing the indicated stress agents: diamide (DIA), CdSO_4_, amphotericin B (AMB), fludioxonil (FDX), and fluconazole (FLC). Plates were imaged after 3 days of incubation at 30°C. (G) Thermotolerance of WT, *mln1*Δ, and *mln1*Δ+*MLN1* strains. Cells were serially diluted and spotted onto YPD agar plates, followed by incubation at 30°C or 39°C. Plates were imaged after 6 days of incubation. (H) Growth kinetics of WT and *mln1*Δ strains at 39°C. Cells were cultured in YPD medium at 39°C, and growth was continuously monitored by measuring OD_600_ in a multi-channel bioreactor. Solid lines represent mean growth curves, and shaded regions indicate variation among 3 biological replicates. (I) Melanin production of WT, *mln1*Δ, and *mln1*Δ+*MLN1* strains. Cells were spotted onto L-DOPA agar supplemented with 0.1% glucose and incubated at 37°C. Plates were imaged after 2 days of incubation. (J) Capsule formation of WT, *mln1*Δ, and *mln1*Δ+*MLN1* strains under capsule-inducing minimal-medium conditions. Cells were incubated in minimal medium containing 10.0 mM MgSO_4_, 29.4 mM KH_2_PO_4_, 13.0 mM glycine, and 3.0 μM thiamine, adjusted to pH 5.5, under the indicated carbon-source conditions. Capsule size was quantified from microscopic images. Violin plots show the distribution of capsule measurements from individual cells. Statistical significance was assessed by ordinary one-way ANOVA followed by Dunnett’s multiple-comparisons test, comparing each strain with WT. *P* values are shown in the panel; statistically significant values (*P* < 0.05) are shown in black, whereas nonsignificant values (*P* ≥ 0.05) are shown in gray.

**Figure S8.**
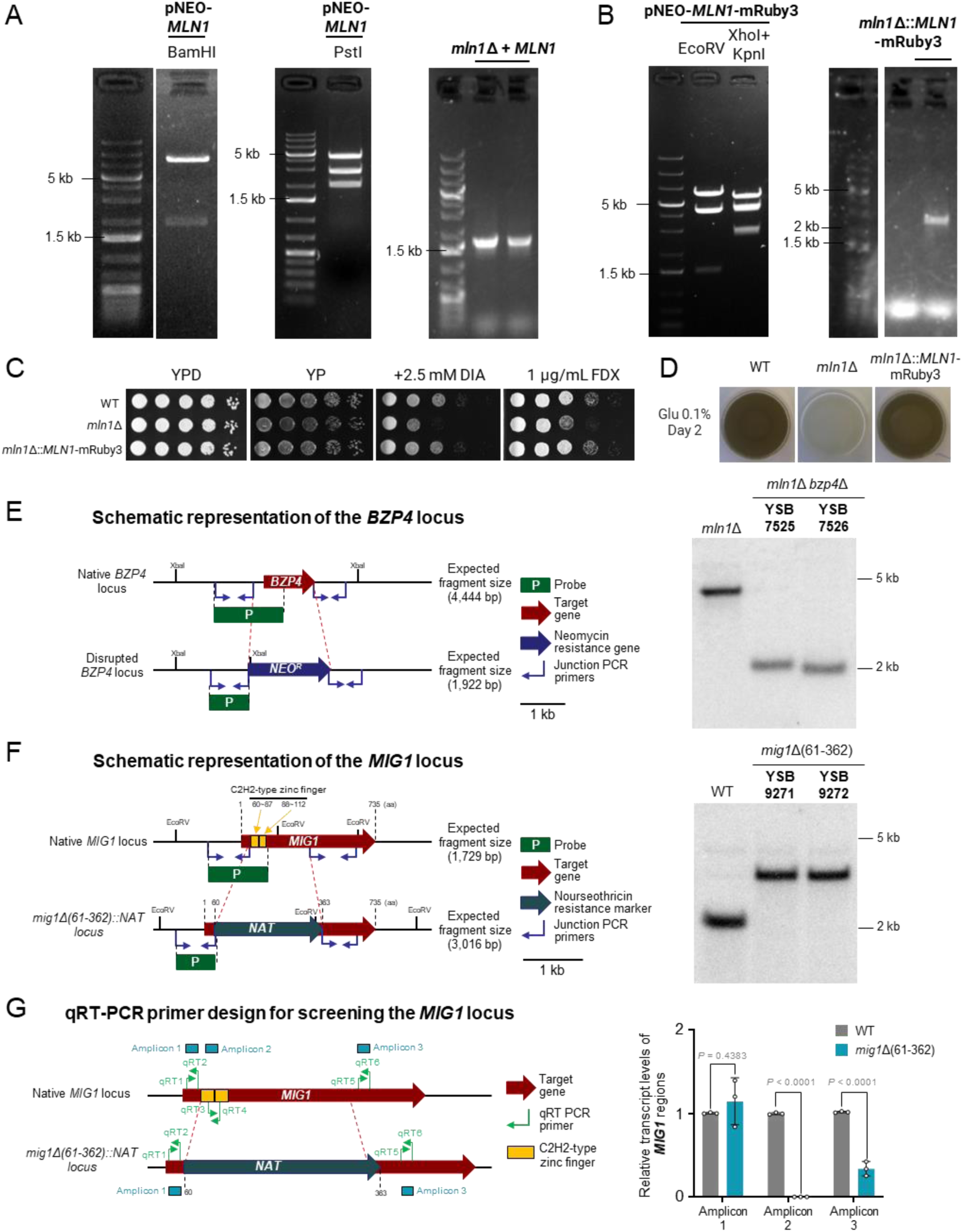
Construction and validation of complemented strains and selected mutant alleles. (A) Construction and verification of the neomycin-resistance (*NEO*)-marked *MLN1* complementation plasmid. The plasmid was confirmed by restriction digestion with BamHI, yielding the expected fragments of 7,689 and 1,789 bp, and with PstI, yielding fragments of 4,580, 3,022, and 1,876 bp. The verified construct was introduced into the *mln1*Δ strain by biolistic transformation, and the presence of the introduced *MLN1* complementation cassette was confirmed by PCR using primers L1 (B1758) and PO (B1764), which produced the expected 1,756-bp amplicon. (B) Construction and verification of the *NEO*-marked *MLN1*-mRuby3 complementation plasmid. The plasmid was confirmed by restriction digestion with EcoRV, yielding the expected fragments of 5,853, 4,526, and 1,462 bp, and with XhoI plus KpnI, yielding fragments of 5,215, 3,997, and 2,629 bp. The verified construct was introduced into the *mln1*Δ strain by biolistic transformation, and targeted integration was confirmed by PCR using primers CSO (B11420) and PO (B1764), which produced the expected 2,298-bp amplicon. (C) Phenotypic complementation of the *mln1*Δ mutant by the *MLN1*-*mRuby3* construct. WT, *mln1*Δ, and the complemented strain were serially diluted and spotted onto the indicated media. Plates were photographed after 2 days of incubation. (D) Restoration of melanization by *MLN1*-mRuby3 complementation. WT, *mln1*Δ, and the complemented strain were cultured on L-DOPA agar containing 0.1% glucose for 2 days under the indicated conditions. (E) Construction and validation of the *mln1*Δ *bzp4*Δ double-deletion mutant. The schematic illustrates replacement of the native *BZP4* locus with the *NEO* marker by homologous recombination in the *mln1*Δ background. Green bars indicate the Southern blot probe, blue arrows indicate junction PCR primers, and red dashed lines indicate the homologous recombination regions. Correct disruption of *BZP4* was confirmed by Southern blot analysis. (F) Genetic organization and validation of the *mig1*Δ(61–362) allele. The schematic illustrates replacement of the region of *MIG1* encoding amino acids 61-362, including the C2H2-type zinc-finger DNA-binding domain, with the nourseothricin-resistance (*NAT*) marker by homologous recombination. Green bars indicate the Southern blot probe, blue arrows indicate junction PCR primers, and red dashed lines indicate the homologous recombination regions. Correct disruption of the *MIG1* locus was confirmed by Southern blot analysis following EcoRV digestion, yielding the expected 1,729-bp fragment in WT and 3,016-bp fragments in two independent mutants. (G) qRT-PCR validation of the *mig1*Δ allele using three primer pairs targeting different regions of the *MIG1* locus. The same three biological replicates of basal samples used for the qRT-PCR analysis in Fig 3D were analyzed using the primer pairs listed in Supporting Information, Table S2. Amplicon 1 targets the 5′ region upstream of the deleted sequence, amplicon 2 targets the C2H2 zinc-finger region removed in *mig1*Δ, and amplicon 3 targets the 3′ region downstream of the deletion. Transcript levels were normalized to *ACT1* and expressed relative to WT for each amplicon. Bars represent the mean ± SD of three biological replicates, and dots indicate individual biological replicates. Statistical significance was assessed by two-way repeated-measures ANOVA followed by Šídák’s multiple-comparisons test comparing WT and *mig1*Δ within each amplicon.

**Figure S9.**
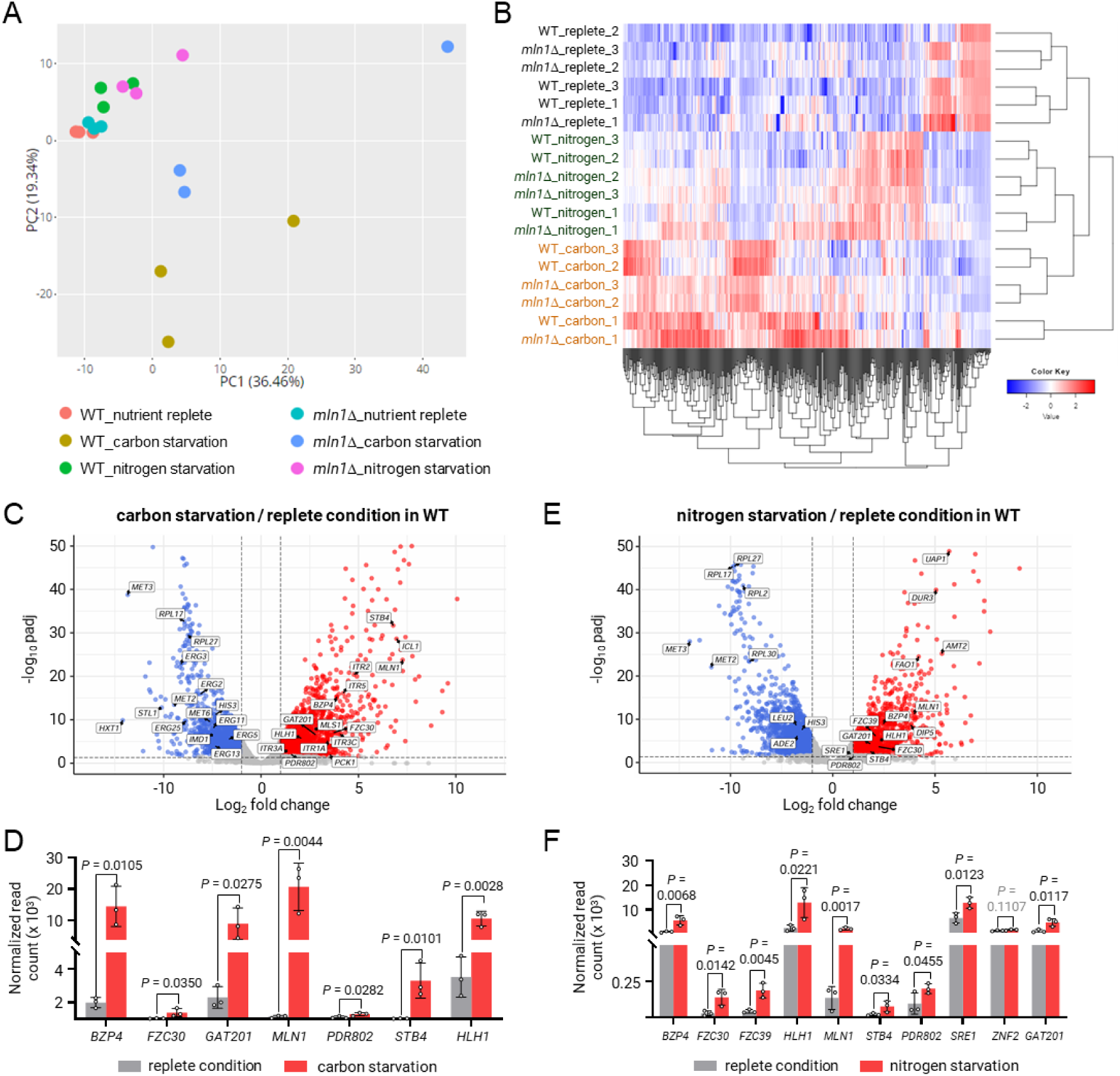
Quality assessment and stress-response validation of the RNA-seq dataset. (A) Principal-component analysis (PCA) of RNA-seq samples from WT and *mln1*Δ cells under nutrient-replete, carbon-starved, and nitrogen-starved conditions. Each dot represents one biological replicate. Samples separate primarily according to nutrient condition and genotype. (B) Hierarchical clustering heatmap of RNA-seq samples based on global transcript abundance. Samples showed nutrient-condition-associated clustering, with intermingling of WT and *mln1*Δ replicates within conditions. (C) Volcano plot showing transcriptional changes in WT cells under carbon starvation relative to nutrient replete conditions. Genes corresponding to carbon-responsive transcription factors (TFs) identified in the host-cue expression screen are labeled. (D) RNA-seq-based expression levels of carbon-starvation-responsive TFs identified in the qRT-PCR-based host-cue screen, together with *HLH1*, which was also induced under carbon starvation in the RNA-seq dataset. Raw read counts were normalized using the DESeq2 median-of-ratios method. Bars represent the mean ± SD, and dots indicate individual biological replicates. Statistical significance was determined using an unpaired one-tailed Student’s t-test comparing nutrient-replete and carbon-starved conditions for each gene. (E) Volcano plot showing transcriptional changes in WT cells under nitrogen starvation relative to nutrient replete conditions. Genes corresponding to nitrogen-responsive TFs identified in the host-cue expression screen are labeled. (F) RNA-seq-based expression levels of nitrogen-starvation-responsive TFs identified in the qRT-PCR-based host-cue screen, together with *GAT201*, which was also induced under nitrogen starvation in the RNA-seq dataset. Raw read counts were normalized using the DESeq2 median-of-ratios method. Bars represent the mean ± SD, and dots indicate individual biological replicates. Statistical significance was determined using an unpaired one-tailed Student’s t-test comparing nutrient-replete and nitrogen-starved conditions for each gene.

**Figure S10.**
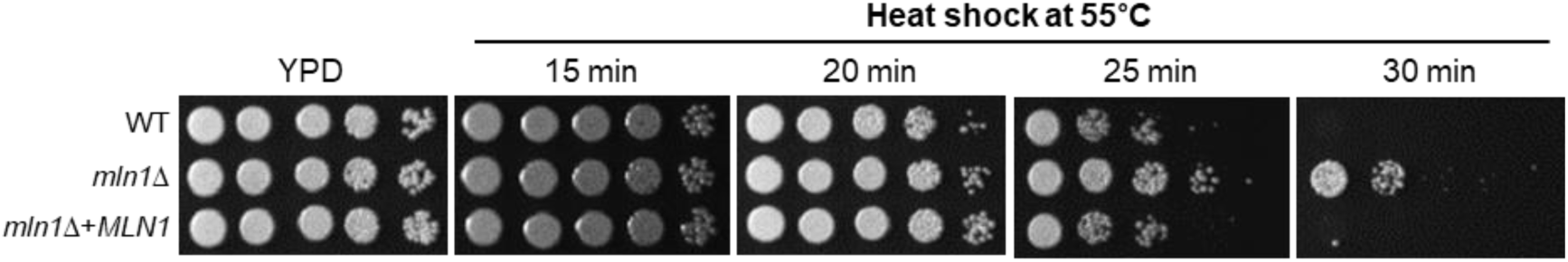
Loss of *MLN1* enhances acute heat-shock resistance. Acute heat-shock resistance of WT, *mln1*Δ, and the complemented strain. Serially diluted cells were spotted onto YPD agar, exposed to 55°C for 15, 20, 25, and 30 min, and subsequently incubated at 30°C for 3 days.

**Table S1.** List of strains used in this study.

| **Strains (YSB #)** | **Genotype** | Parent | Reference |
|---|---|---|---|
| H99 | *MAT*α |  | [19] |
| YSB1894 | *MAT*α *bzp4*Δ::*NAT*-STM #295 | H99 | [3] |
| YSB2447 | *MAT*α *fzc30*Δ::*NAT*-STM #230 | H99 | [3] |
| YSB1820 | *MAT*α *fzc39*Δ::*NAT*-STM #231 | H99 | [3] |
| YSB3300 | *MAT*α *gat201*Δ::*NAT*-STM #273 | H99 | [3] |
| YSB1175 | *MAT*α *hlh1*Δ::*NAT*-STM #146 | H99 | [3] |
| YSB1585 | *MAT*α *hob5*Δ::*NAT*-STM #219 | H99 | [3] |
| YSB1172 | *MAT*α *mln1*Δ::*NAT*-STM #146 | H99 | [3] |
| YSB1173 | *MAT*α *mln1*Δ::*NAT*-STM #146 | H99 | [3] |
| YSB2387 | *MAT*α *pdr802*Δ::*NAT*-STM #220 | H99 | [3] |
| YSB2494 | *MAT*α *sre1*Δ::*NAT*-STM #240 | H99 | [3] |
| YSB1013 | *MAT*α *stb4*Δ::*NAT*-STM #125 | H99 | [3] |
| YSB2740 | *MAT*α *znf2*Δ::*NAT*-STM #211 | H99 | [3] |
| YSB2372 | *MAT*α *snf1*Δ::*NAT*-STM #204 | H99 | [20] |
| YSB9271 | *MAT*α *mig1Δ*(61-362)::*NAT*-STM #210 | H99 | This study |
| YSB9272 | *MAT*α *mig1Δ*(61-362)::*NAT*-STM #210 | H99 | This study |
| YSB7671 | *MAT*α *fzc30*Δ::*NAT bzp4*Δ::*NEO* | YSB2447 | This study |
| YSB7672 | *MAT*α *fzc30*Δ::*NAT bzp4*Δ::*NEO* | YSB2447 | This study |
| YSB7928 | *MAT*α *hob5*Δ::*NAT bzp4*Δ::*NEO* | YSB1585 | This study |
| YSB7716 | *MAT*α *hob5*Δ::*NAT bzp4*Δ::*NEO* | YSB1585 | This study |
| YSB7650 | *MAT*α *pdr802*Δ::*NAT bzp*4Δ::*NEO* | YSB2387 | This study |
| YSB7651 | *MAT*α *pdr802*Δ::*NAT bzp4*Δ::*NEO* | YSB2387 | This study |
| YSB8264 | *MAT*α *znf2*Δ::*NAT bzp4*Δ::*NEO* | YSB2740 | This study |
| YSB8265 | *MAT*α *znf2*Δ::*NAT bzp*4Δ::*NEO* | YSB2740 | This study |
| YSB7931 | *MAT*α *bzp4*Δ::*NAT fzc39*Δ::*NEO* | YSB1894 | This study |
| YSB7932 | *MAT*α *bzp4*Δ::*NAT fzc39*Δ::*NEO* | YSB1894 | This study |
| YSB8274 | *MAT*α *fzc30*Δ::*NAT fzc39*Δ::*NEO* | YSB2447 | This study |
| YSB8275 | *MAT*α *fzc30*Δ::*NAT fzc39*Δ::*NEO* | YSB2447 | This study |
| YSB7761 | *MAT*α *hob5*Δ::*NAT fzc39*Δ::*NEO* | YSB1585 | This study |
| YSB7763 | *MAT*α *hob5*Δ::*NAT fzc39*Δ::*NEO* | YSB1585 | This study |
| YSB7759 | *MAT*α *znf2*Δ::*NAT fzc39*Δ::*NEO* | YSB2740 | This study |
| YSB7929 | *MAT*α *znf2*Δ::*NAT fzc39*Δ::*NEO* | YSB2740 | This study |
| YSB7673 | *MAT*α *fzc30*Δ::*NAT pdr802*Δ::*NEO* | YSB2447 | This study |
| YSB7675 | *MAT*α *fzc30*Δ::*NAT pdr802*Δ::*NEO* | YSB2447 | This study |
| YSB7598 | *MAT*α *fzc39*Δ::*NAT pdr802*Δ::*NEO* | YSB1820 | This study |
| YSB7599 | *MAT*α *fzc39*Δ::*NAT pdr802*Δ::*NEO* | YSB1820 | This study |
| YSB7077 | *MAT*α *hob5*Δ::*NAT pdr802*Δ::*NEO* | YSB1585 | This study |
| YSB7078 | *MAT*α *hob5*Δ::*NAT pdr802*Δ::*NEO* | YSB1585 | This study |
| YSB7656 | *MAT*α *znf2*Δ::*NAT pdr802*Δ::*NEO* | YSB2740 | This study |
| YSB7783 | *MAT*α *znf2*Δ::*NAT pdr802*Δ::*NEO* | YSB2740 | This study |
| YSB7922 | *MAT*α *hob5*Δ::*NAT fzc30*Δ::*NEO* | YSB1585 | This study |
| YSB7923 | *MAT*α *hob5*Δ::*NAT fzc*30Δ::*NEO* | YSB1585 | This study |
| YSB7676 | *MAT*α *znf2*Δ::*NAT fzc30*Δ::*NEO* | YSB2740 | This study |
| YSB7677 | *MAT*α *znf2*Δ::*NAT fzc30*Δ::*NEO* | YSB2740 | This study |
| YSB7731 | *MAT*α *hob5*Δ::*NAT znf2*Δ::*NEO* | YSB1585 | This study |
| YSB7760 | *MAT*α *hob5*Δ::*NAT znf2*Δ::*NEO* | YSB1585 | This study |
| YSB7525 | *MAT*α *mln1*Δ::*NAT bzp4*Δ::*NEO* | YSB1172 | This study |
| YSB7526 | *MAT*α *mln1*Δ::*NAT bzp4*Δ::*NEO* | YSB1172 | This study |
| YSB10818 | *MAT*α *mln1*Δ::*MLN1*-mRuby3::*NEO* | YSB1172 | This study |
| YSB11310 | *MAT*α *mln1*Δ::*NAT*+*MLN1*-*NEO* | YSB1172 | This study |
| YSB42 | *MAT*α *cac1*Δ::*NAT*-STM #159 | H99 | [21] |

**Table S2.**
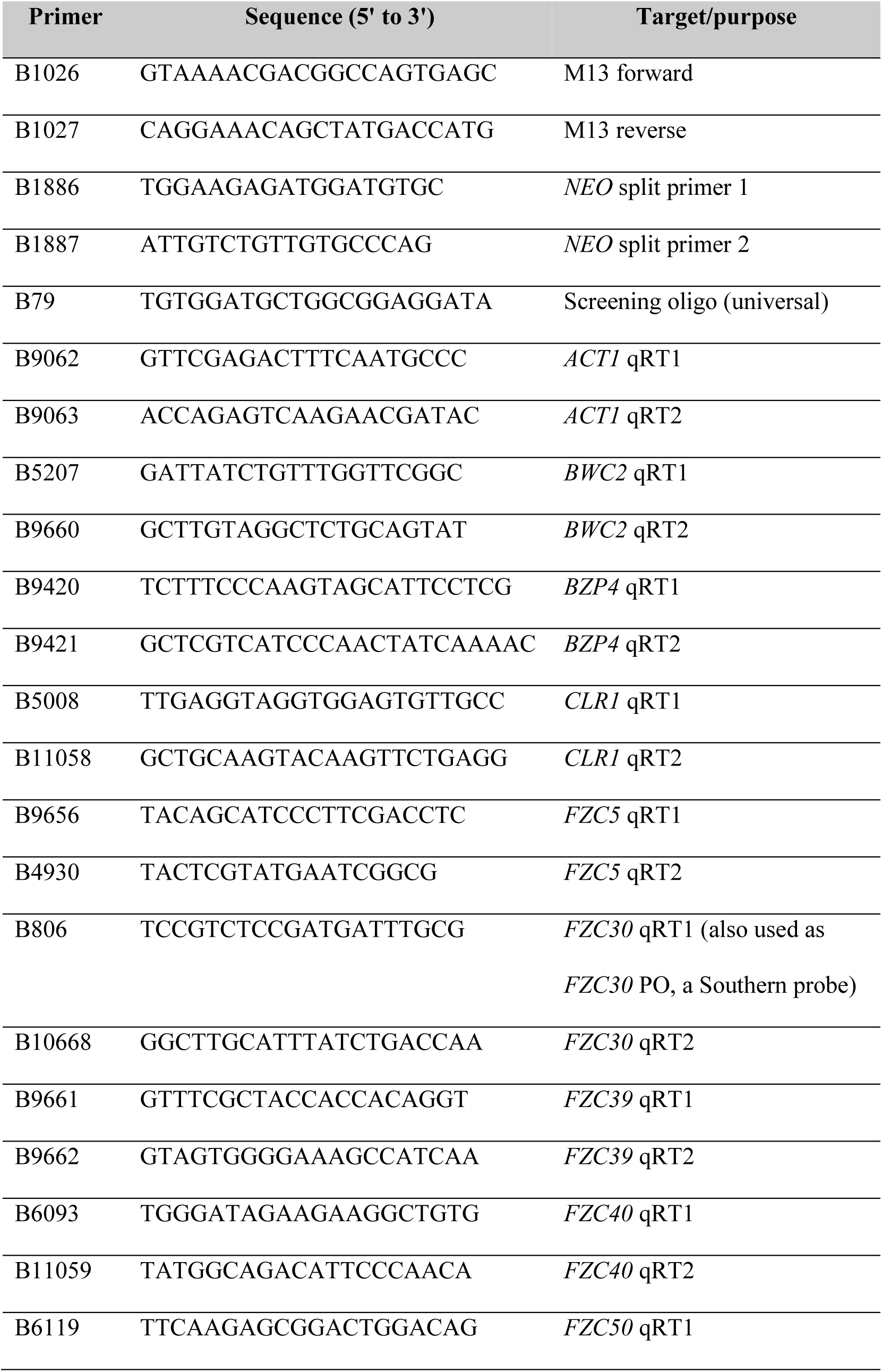
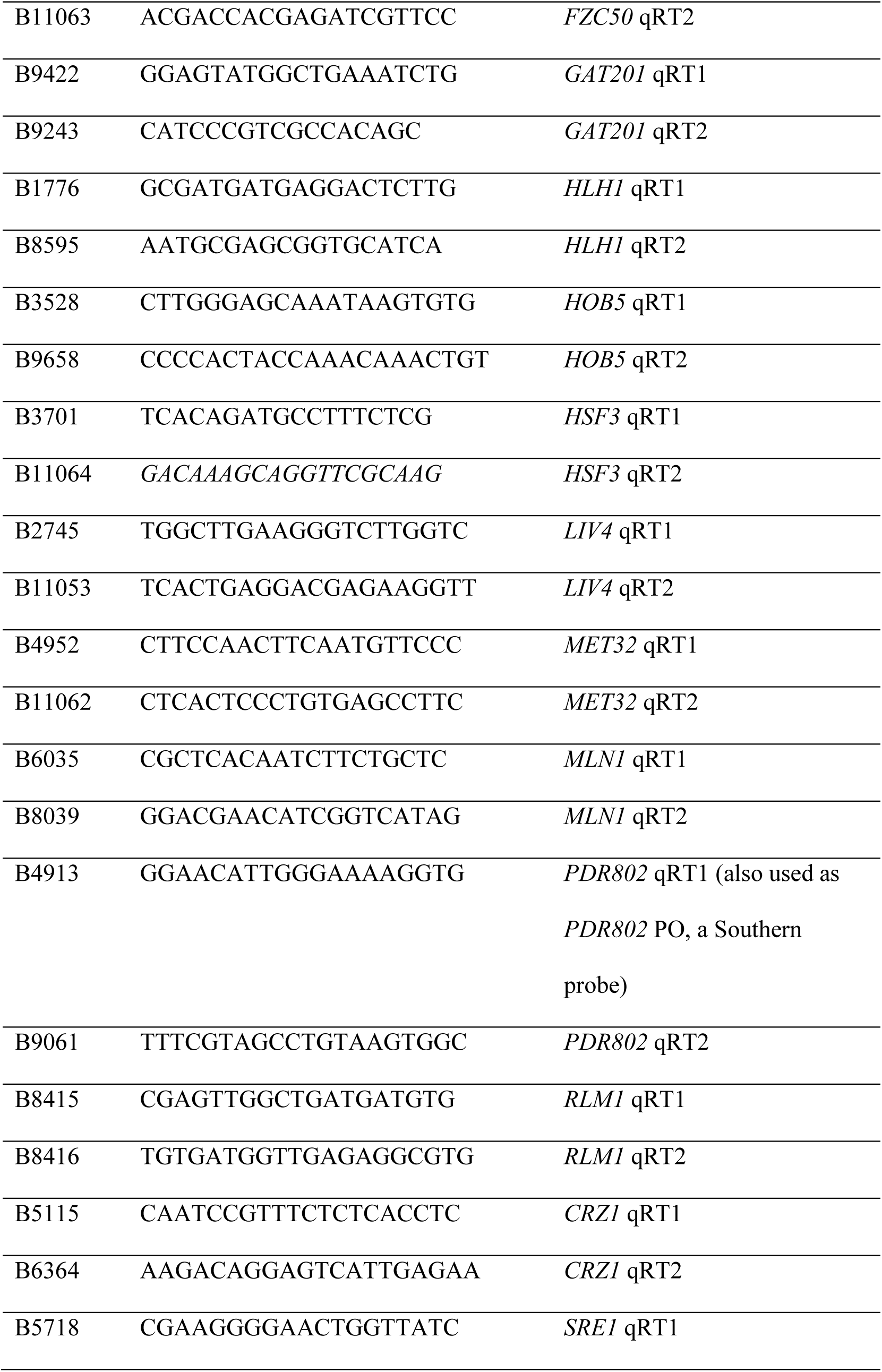
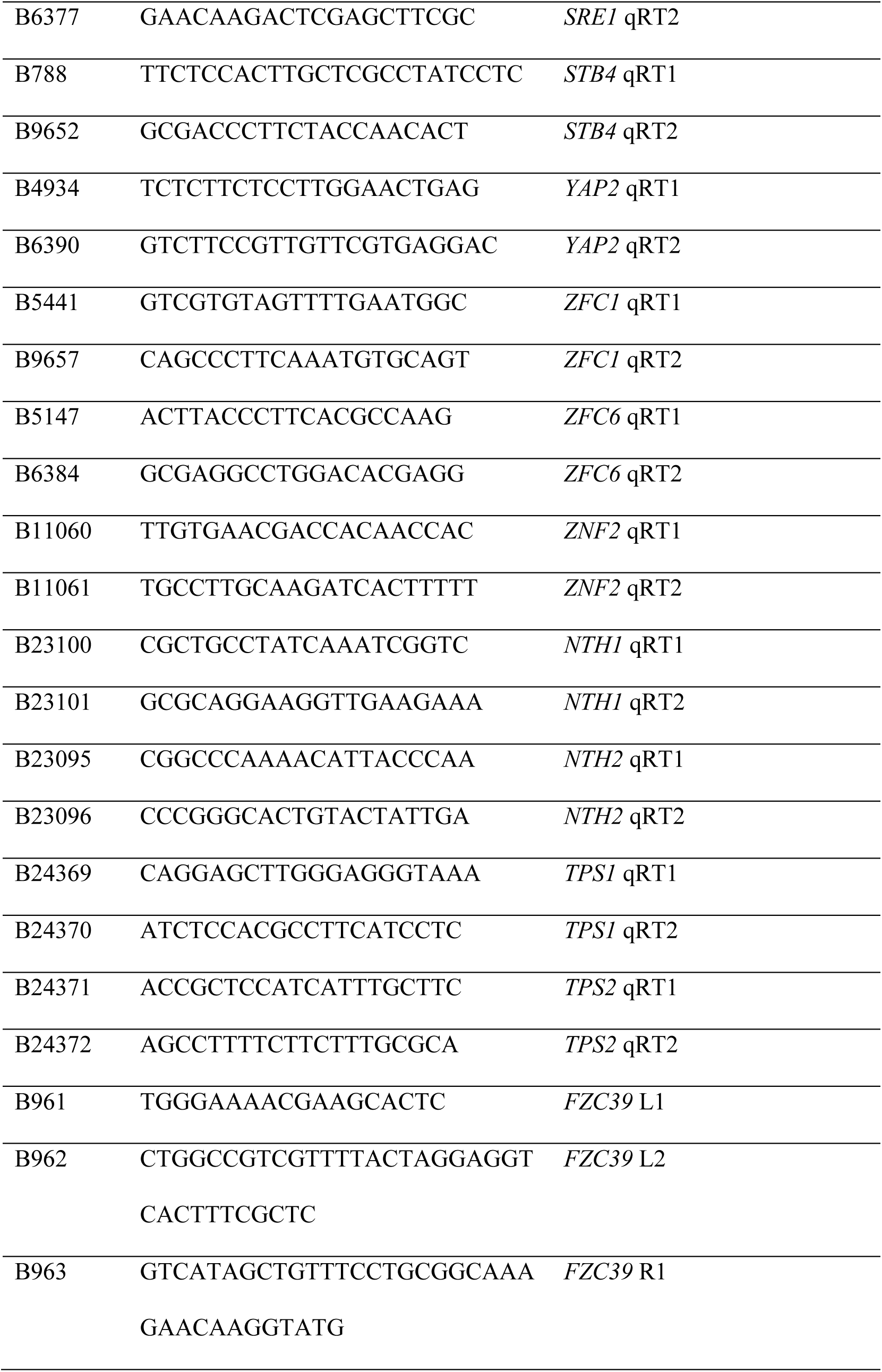
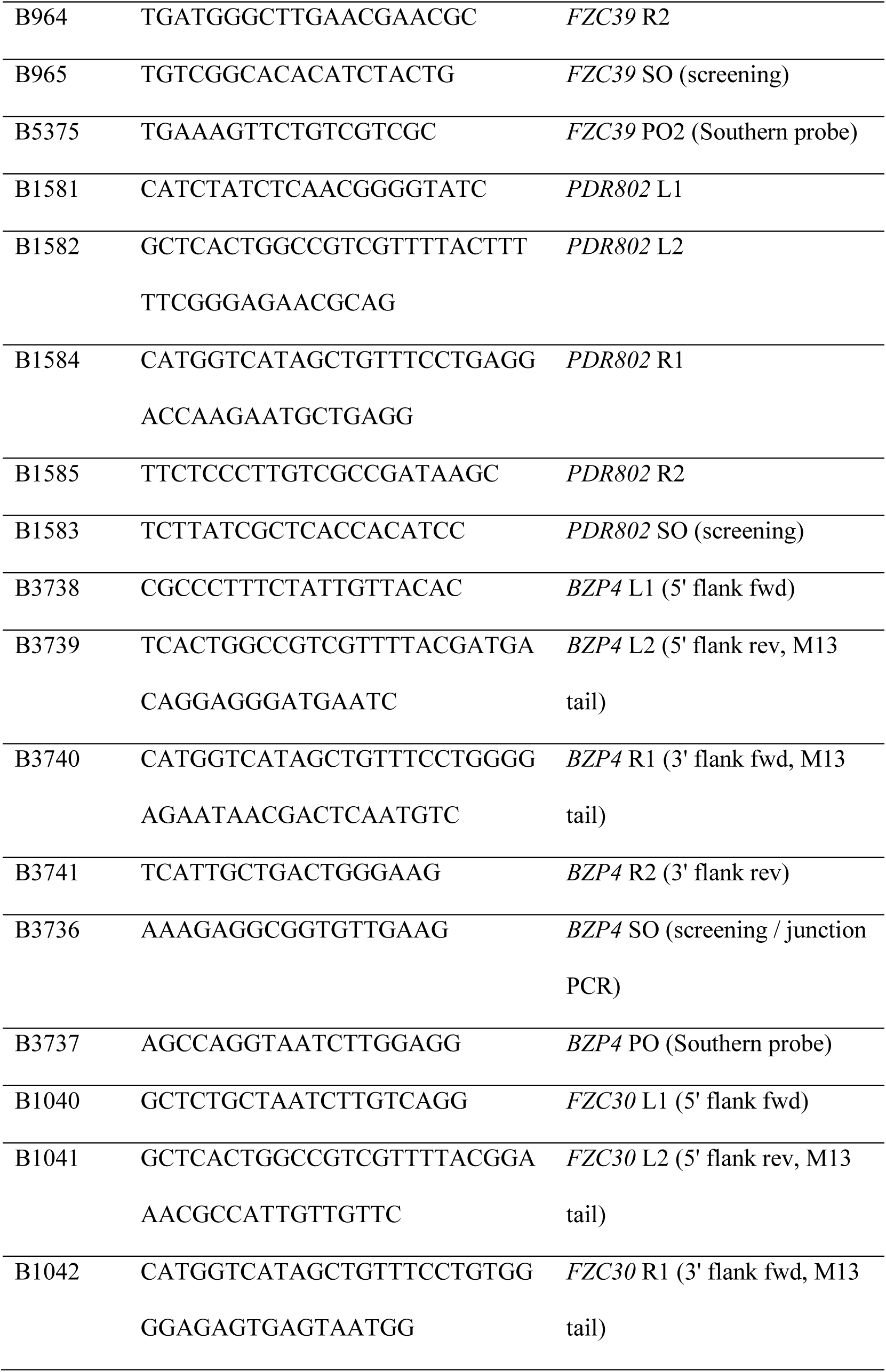
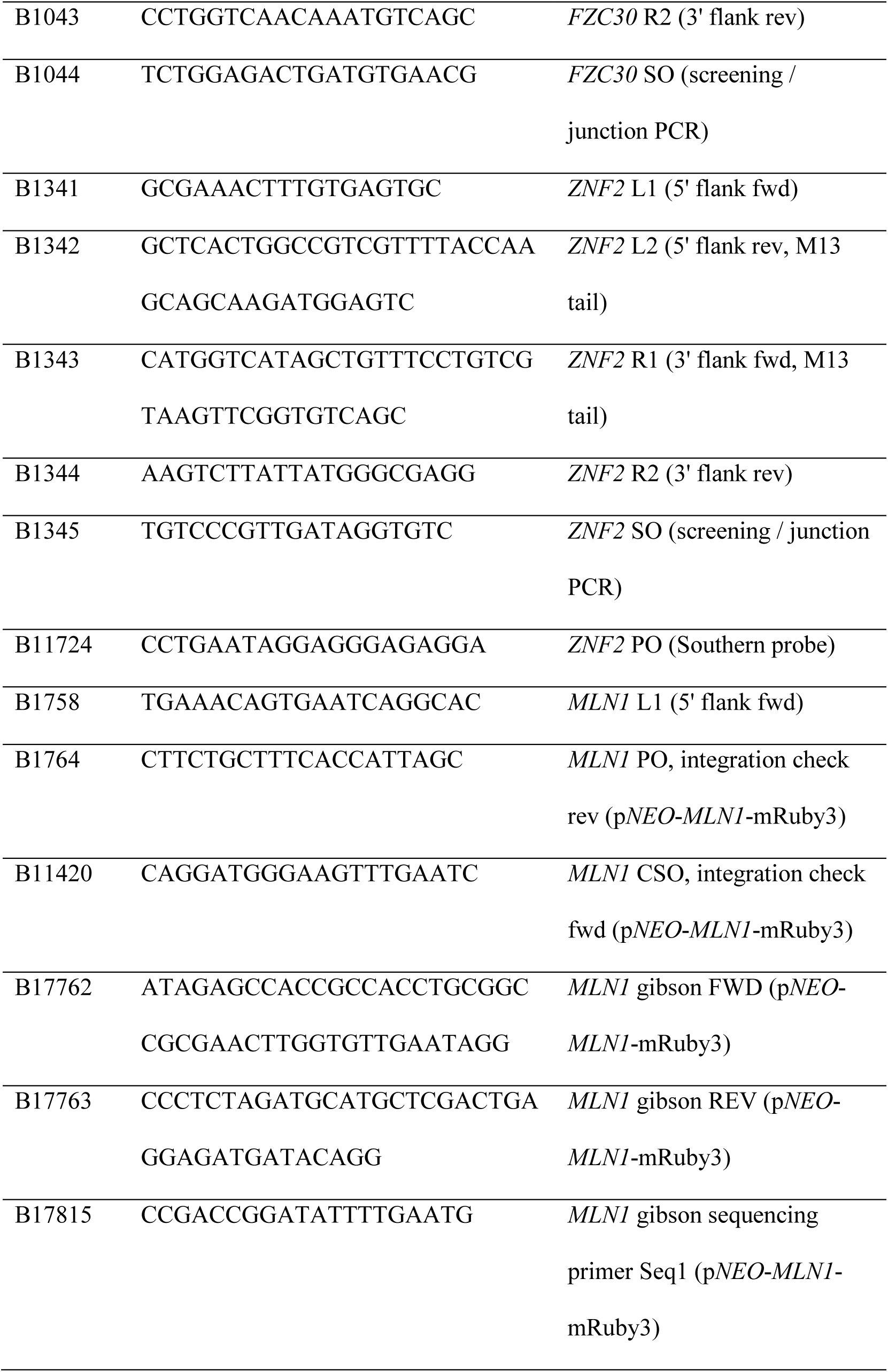
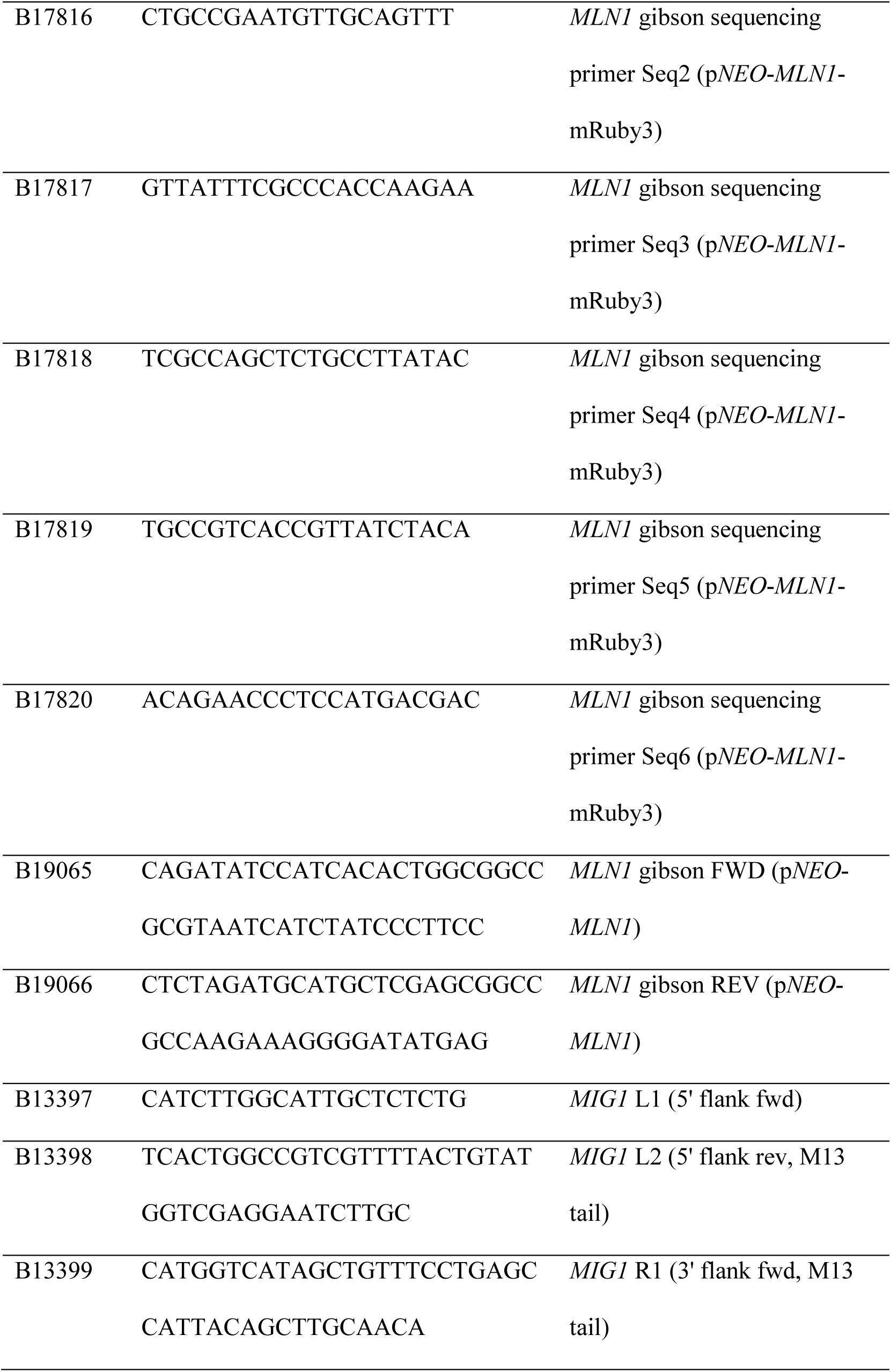
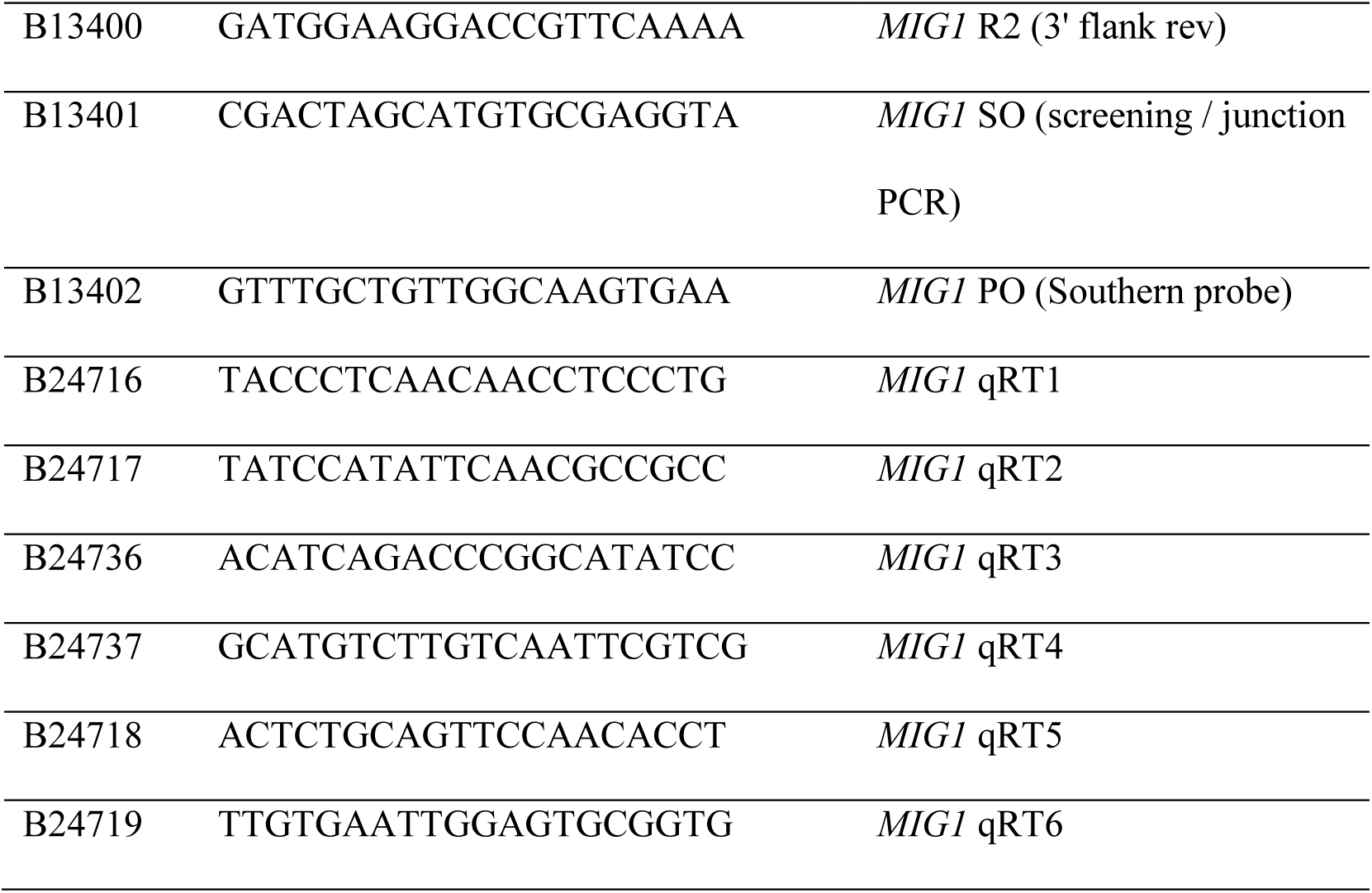
Primers used in this study.

**Table S3.** Media compositions used in this study.

| **Condition** | **Media composition** |
|---|---|
| RPMI condition | RPMI1640 only |
|  | RPMI1640 + 10% FBS |
|  | RPMI1640 + 2% glucose |
|  | RPMI1640 + 10% FBS + 2% glucose |
| YPD condition | Yeast extract-peptone |
|  | Yeast extract-peptone + 2% glucose |
|  | Yeast extract-peptone + 2% glucose + 10% FBS |
| YNB condition | Yeast-nitrogen base (YNB) without amino acids and with ammonium sulfate (AS) + 2% glucose |
|  | YNB without amino acids (AA) and ammonium sulfate (AS) + 2% glucose |
| MM | Yeast-nitrogen base (YNB) without amino acids |
| MMG | Yeast-nitrogen base (YNB) without amino acids + 2% glucose |

**Dataset S1 (separate file). Integrated transcriptomic and infectivity data used for the stepwise selection of pathogenicity-related transcription factors in *C. neoformans*.**

**Dataset S2 (separate file). RNA-seq analysis of Mln1-dependent gene expression under basal, carbon-starved, and nitrogen-starved conditions.**

